# Partial inhibition of mitochondrial complex I attenuates neurodegeneration and restores energy homeostasis and synaptic function in a symptomatic Alzheimer’s mouse model

**DOI:** 10.1101/2020.07.01.182428

**Authors:** Andrea Stojakovic, Sergey Trushin, Anthony Sheu, Layla Khalili, Su-Youne Chang, Xing Li, Trace Christensen, Jeffrey L. Salisbury, Rachel E. Geroux, Benjamin Gateno, Padraig J. Flannery, Mrunal Dehankar, Cory C. Funk, Jordan Wilkins, Anna Stepanova, Tara O’Hagan, Alexander Galkin, Jarred Nesbitt, Xiujuan Zhu, Utkarsh Tripathi, Slobodan Macura, Tamar Tchkonia, Tamar Pirtskhalava, James L. Kirkland, Rachel A. Kudgus, Renee A. Schoon, Joel M. Reid, Yu Yamazaki, Takahisa Kanekiyo, Song Zhang, Emirhan Nemutlu, Petras Dzeja, Adam Jaspersen, Christopher Ye In Kwon, Michael K. Lee, Eugenia Trushina

## Abstract

We demonstrate that mitochondrial respiratory chain complex I is an important small molecule druggable target in Alzheimer’s Disease (AD). Partial inhibition of complex I triggers the AMP-activated protein kinase-dependent signaling network leading to neuroprotection in symptomatic APP/PS1 mice, a translational model of AD. Treatment of APP/PS1 mice with complex I inhibitor after the onset of AD-like neuropathology improved energy homeostasis, synaptic activity, long-term potentiation, dendritic spine maturation, cognitive function and proteostasis, and reduced oxidative stress and inflammation in brain and periphery, ultimately blocking the ongoing neurodegeneration. Therapeutic efficacy *in vivo* was monitored using translational biomarkers FDG-PET, ^31^P NMR, and metabolomics. Cross-validation of the mouse and the human AMP-AD transcriptomic data demonstrated that pathways improved by the treatment in APP/PS1 mice, including the immune system response and neurotransmission, represent mechanisms essential for therapeutic efficacy in AD patients.

---

Alzheimer’s Disease (AD) is a multifactorial disorder without a cure. It is characterized by progressive accumulation of aggregated amyloid β (Aβ) peptides and hyperphosphorylated Tau protein, memory decline, and neurodegeneration. The consistent failure of clinical trials focused on reducing Aβ levels and aggregation suggests that such therapies may not work in AD patients regardless of disease stage, underscoring the need to discover novel targets and therapies for AD^1,2^. Recent studies demonstrated that altered energy homeostasis associated with reduced cerebral glucose uptake and utilization, altered mitochondrial function and microglia and astrocyte activation might underlie neuronal dysfunction in AD^3–7^. Intriguingly, accumulating evidence suggests that non-pharmacological approaches, such as diet and exercise, reduce major AD hallmarks by engaging an adaptive stress response that leads to improved metabolic state, reduced oxidative stress and inflammation, and improved proteostasis^8^. While mechanisms of the stress response are complex, AMPK-mediated signaling has been directly linked to the regulation of cell metabolism, mitochondrial dynamics and function, inflammation, oxidative stress, protein turnover, Tau phosphorylation, and amyloidogenesis^9^. Combined analysis performed using multiple types of genome-wide data identified a predominant role for metabolism-associated biological processes in the course of AD, including autophagy and insulin and fatty acid metabolism, with a focus on AMPK as a key modulator and therapeutic target^10^. However, the development of direct pharmacological AMPK activators to elicit beneficial effects has presented multiple challenges^11^. We recently demonstrated that mild inhibition of mitochondrial complex I (MCI) with the small molecule tricyclic pyrone compound, CP2 blocked cognitive decline in transgenic mouse models of AD when treatment was started *in utero* through life or at a pre-symptomatic stage of the disease^12,13^. Moreover, in neurons, CP2 restored mitochondrial dynamics and function and cellular energetics. However, it was unclear whether MCI inhibition would elicit similar benefits if administered at the advanced stage of the disease, after the development of prominent Aβ accumulation, brain hypometabolism, cognitive dysfunction, and progressive neurodegeneration. As a proof of concept, we demonstrate that partial inhibition of MCI triggers stress-induced AMPK-dependent signaling cascade leading to neuroprotection and a reversal of behavior changes in symptomatic APP/PS1 mice, a translational model of AD.

## Results

### CP2 activates AMPK-dependent neuroprotective pathways and restores cognitive and motor function in symptomatic APP/PS1 mice

The tricyclic pyrone, CP2, specifically inhibits the activity of MCI in human and mouse brain mitochondria^12^ (Extended Data Fig. 1a-c). CP2 penetrates the blood-brain barrier (BBB) and accumulates in mitochondria, mildly decreasing MCI activity, which leads to an increase in AMP/ATP ratio and AMPK activation^12^. CP2 was effective in blocking cognitive dysfunction when treatment was administered to pre-symptomatic mice carrying familial mutations in the APP(K670N/M671L) and PS1(M146L) genes (APP/PS1)^12,13^. To determine whether CP2 could engage AMPK-dependent neuroprotective mechanisms (Extended Data Fig. 1d) in symptomatic mice, we administered a single oral dose to 9 −10-month-old APP/PS1 mice and examined the expression of key proteins in each pathway after 4, 24, 48, and 72 h (Extended Data Fig. 1e-l, Extended Data Fig. 2a, Supplementary Fig. 1). An independent cohort of CP2-treated APP/PS1 mice was assayed using *in vivo* 18F-fluorodeoxyglucose positron emission tomography (FDG-PET) and compared to non-transgenic (NTG) untreated littermates (Extended Data Fig. 1h, 2b). Consistent with previous observations, CP2 robustly activated AMPK after 24 h, when increased phosphorylation of acetyl-CoA carboxylase 1 (ACC1), a biomarker of AMPK target engagement associated with increased fatty acid oxidation^14^, was evident 4 h after CP2 administration (Extended Data Fig. 2a). Remarkably, a significant increase in the glucose transporters, Glut 3 and 4, was observed as early as 4 h after CP2 treatment that persisted for 24 and 48 h, consistent with the established role of AMPK in the maintenance of glucose uptake in the brain^15–17^ (Extended Data Fig. 2a). Improved glucose uptake in APP/PS1 mice was independently established using *in vivo* FDG-PET (Extended Data Fig. 2b). A decreased ratio of phosphorylated *vs.* total pyruvate dehydrogenase (PDH) at 24 and 48 h confirmed that augmented glucose uptake was associated with improved glucose utilization (Extended Data Fig. 1g, 2a), since increased PDH activity leads to acetyl-CoA production from pyruvate in the glucose catabolism pathway, promoting energy production in mitochondria^18^. Furthermore, increased expression of the transcriptional coactivator, peroxisome proliferator-activated receptor-γ coactivator-1α (PGC-1α), mitochondrial transcription factor A (TFAM), and mitochondria-specific neuroprotective Sirtuin 3 (Sirt3), supports the view that there was improved mitochondrial biogenesis and function (Extended Data Fig. 1i)^19,20^. We further confirmed that CP2 induces an anti-inflammatory mechanisms evident by increased levels of IκBα and decreased phosphorylation of p65, which together block NF-κB transcription factor activation (Extended Data Fig. 1j)^21^. Additionally, CP2 activates nuclear factor E2-related factor 2 (Nrf 2)-dependent expression of antioxidants including heme oxygenase (HO1), superoxide dismutase (SOD1), and catalases (Extended Data Fig. 1k)^22^. Finally, CP2 treatment increased autophagy based on changes in the autophagy activating kinase ULK, Beclin and LC3B proteins essential for autophagosome formation (Extended Data Fig. 1l)^23^. These data demonstrate that oral CP2 administration activates the neuroprotective AMPK signaling network in symptomatic APP/PS1 mice at therapeutic doses.

To determine the therapeutic efficacy of chronic CP2 administration, we treated symptomatic APP/PS1 female mice with CP2 (25 mg/kg/day in drinking water *ad lib*) from 9 until 23 months of age (Fig. 1a). NTG female age-matched littermates were controls. All treated mice tolerated CP2 well; they did not manifest side effects and gained weight throughout the duration of the study with an increase in lean and fat mass (Fig. 1b, Extended Data Fig. 3). Results of behavioral tests confirmed that CP2-treated APP/PS1 mice had improved spatial memory and learning (Morris Water Maze, Fig. 1c,d), restoration of attention and non-spatial declarative memory (Novel Object Recognition, Fig. 1e), reduced hyperactivity in open field testing (Fig. 1f), and increased strength and motor coordination (rotarod, hanging bar, Fig. 1g,h). CP2 penetrates the BBB^12,13^ and has oral bioavailability of 65% (Extended Data Fig. 4). CP2 concentrations measured in the brain at the end of the study averaged at ~62 nM, (Supplementary Table 1), consistent with earlier bioavailability studies^12,13^. To establish CP2 selectivity and specificity, we conducted *in vitro* pharmacological profiling against 44 human targets, including G-protein-coupled receptors, ion channels, enzymes, neurotransmitter transporters, and 250 kinases (Supplementary Tables 2-4). At 1 and 10 μM, concentrations higher than those found in the brain tissue of chronically treated mice (Supplementary Table 1), CP2 had minimal off-target activities demonstrating its selectivity at therapeutic doses. These observations reveal that chronic treatment with MCI inhibitor CP2 improves cognitive and motor function in APP/PS1 mice to that of NTG mice after the onset of significant accumulation of Aβ plaques^24^, and development of behavioral^25^ and mitochondrial dysfunction^26^.

**Fig. 1.**
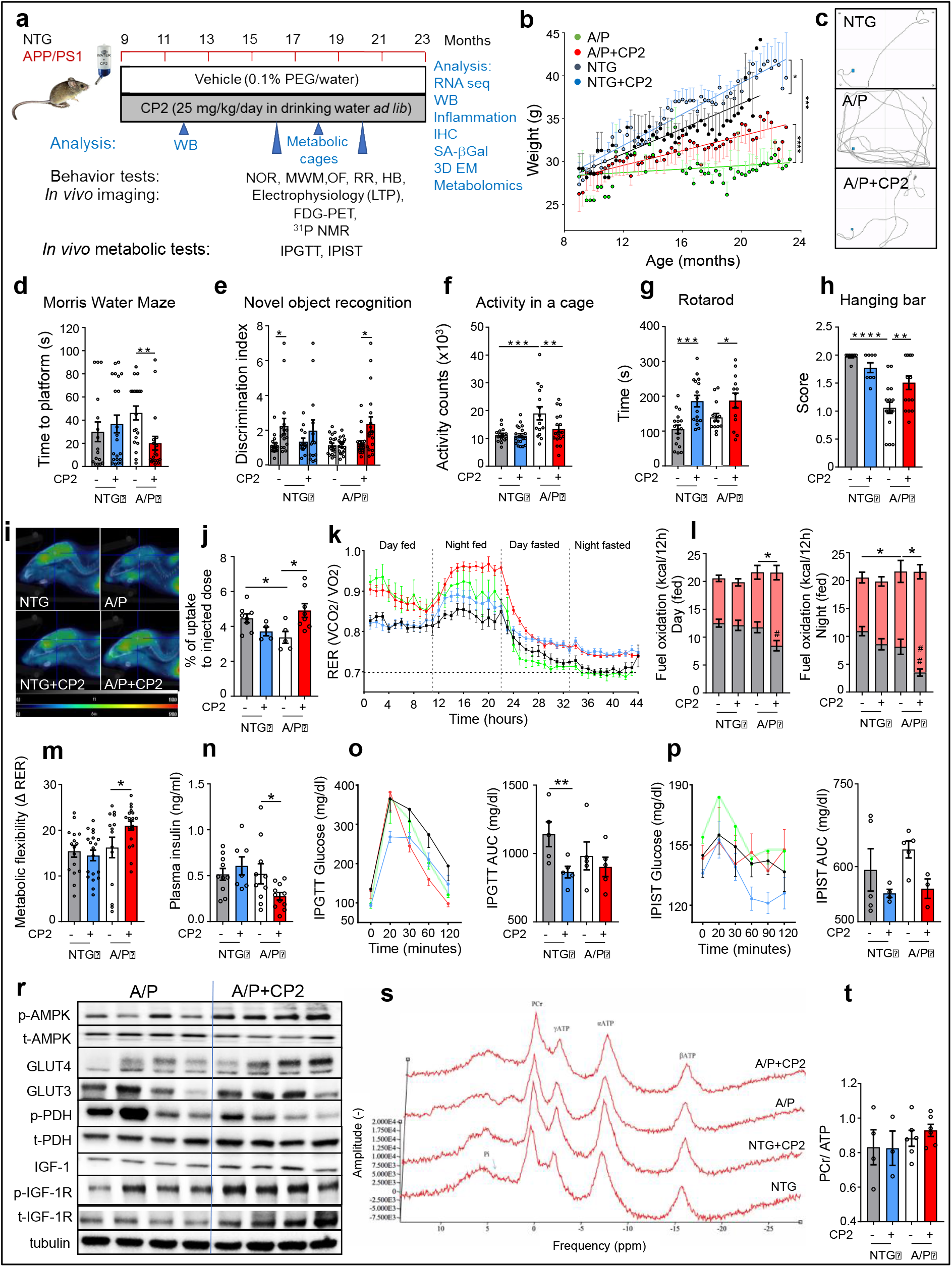
CP2 treatment restores cognitive function and increases glucose uptake and utilization in symptomatic APP/PS1 mice. **a,** Timeline of chronic CP2 treatment. **b**, Weight of NTG and APP/PS1 mice treated with vehicle or CP2 through the duration of the study. **c-h**, CP2 treatment improves performance in the Morris Water Maze (**c, d**) and in the Novel Object Recognition test (**e**), reduces hyperactivity of APP/PS1 mice in open field test (**f**), and increases motor strength and coordination on the rotating rod (**g**) and hanging bar (**h**). *n* = 17 - 20 mice *per* group. **i**, Glucose uptake was increased in the brain of CP2-treated APP/PS1 mice measured using FDG-PET after 9 months of treatment. **j**, Quantification of glucose uptake by FDG-PET imaging from (**i**). *n* = 5 - 8 mice *per* group. **k**, Changes in respiratory exchange ratio (RER) recorded in all treatment groups over 44 h during *ad lib* fed and fasting states. **l**, Glucose oxidation was increased in CP2-treated APP/PS1 mice fed *ad lib* based on CLAMS data from (**k**). Grey bars indicate fat consumption; orange bars indicate carbohydrate and protein oxidation. **m**, Metabolic flexibility is increased in CP2-treated APP/PS1 mice based on their ability to switch from carbohydrates to fat between feeding and fasting states. **k-m**, *n* = 15 - 20 mice *per* group. **n-p**, CP2 treatment reduces fasting insulin levels in plasma of APP/PS1 mice (**n**); increases glucose tolerance in NTG mice measured by intraperitoneal glucose tolerance test (IPGTT) (**o**); and displays tendency to improve intraperitoneal insulin sensitivity test (IPIST) in NTG and APP/PS1 mice (**p**) after 9 - 10 months of treatment. *n* = 5 - 10 mice *per* group. **r,** Western blot analysis conducted in the brain tissue of APP/PS1 mice treated with CP2 for 13 months indicates increased IGF-1signaling, expression of Glut 3 and 4 transporters and changes in pyruvate dehydrogenase (PDH) activation associated with glucose utilization in the TCA cycle. **s,** Representative ^31^P NMR spectra with peaks corresponding to energy metabolites, including inorganic phosphate (Pi), phosphocreatine (PCr), and three phosphate group peaks for ATP generated in living NTG and APP/PS1 mice after 9 months of vehicle or CP2 treatment. **t**, Phosphocreatine/ATP ratio calculated based on the ^31^P NMR *in vivo* spectra from (**s**). *n* = 4 - 6 mice *per* group. Data are presented as mean ± S.E.M. Data were analyzed by two-way ANOVA with Fisher`s LSD *post-hoc* test. A paired Student *t*-test was used for statistical analysis of NOR test. *P < 0.05, **P < 0.01, ***P < 0.001, ****P < 0.0001; # < 0.05, ## < 0.01, for comparison of fat oxidation between vehicle and CP2-treated groups among same genotype. Body weight data in (**b**) were analyzed by linear regression analysis: differences between slopes for NTG *vs*. NTG+CP2 (P < 0.05); A/P *vs*. A/P+CP2 (P < 0.0001); NTG *vs*. A/P+CP2 (P < 0.0001). In all graphs: A/P, APP/PS1, green; NTG, non-transgenic littermates, black; NTG+CP2, blue; APP/PS1+CP2, red.

### CP2 treatment improves glucose uptake and utilization and metabolic flexibility in symptomatic APP/PS1 mice

We next determined effects of chronic CP2 treatment on brain energy homeostasis. Similarly to AD patients, glucose utilization in the brain of APP/PS1 mice measured by FDG-PET imaging was significantly reduced (Fig. 1i,j). Consistent with results of the acute administration (Extended Data Fig. 1h), chronic CP2 treatment over 9 months alleviated pronounced brain glucose hypometabolism in APP/PS1 mice (Fig. 1i,j). Data generated using indirect calorimetry (CLAMS) provided further evidence that, compared to untreated counterparts, CP2 treatment in APP/PS1 mice increased carbohydrate oxidation and metabolic flexibility, an essential ability to switch between lipid and carbohydrate oxidation that is affected in metabolic diseases and aging (Fig. 1k-m). Consistent with the improved regulation of glucose metabolism, CP2-treated APP/PS1 and NTG mice had decreased fasting plasma insulin levels and better insulin sensitivity and glucose tolerance (Fig. 1n-p). Western blot analysis in brain tissue revealed increased expression of Glut 3 and 4 and a decreased ratio of pPDH/PDH indicative of improved glucose uptake and utilization together with enhanced signaling through the IGF pathway (Fig. 1r, Extended Data Fig. 5, Supplementary Fig. 2).

Since CP2 inhibits MCI, we examined if chronic treatment affects ATP levels in the brain using ^31^P nuclear magnetic resonance (^31^P NMR) spectroscopy (Fig. 1s,t). This method allows non-invasive measurement of *in vivo* energy metabolite concentrations including phosphocreatine (PCr), inorganic phosphates (Pi), and the α, β and γ phosphate groups of ATP (Fig. 1s). Chronic CP2 treatment over 10 months did not decrease the PCr /ATP ratio in APP/PS1 or NTG mice (Fig. 1t) consistent with improved glucose uptake/utilization. As a second measure, we performed metabolomic profiling in brain of vehicle- and CP2-treated APP/PS1 and NTG mice treated with CP2 for 6 months (Supplementary Tables 5). Consistent with target engagement, CP2 treatment increased levels of AMP in APP/PS1 mice but did not reduce brain levels of ATP. Treatment resulted in increased levels of citrate and N-acetyl aspartate (NAA), markers of improved mitochondrial and neuronal function. Importantly, levels of 2-hydroxyglutarate, a marker of detrimental mitochondrial stress^27^, were not elevated, suggesting inhibition of MCI with CP2 does not induce adverse mitochondrial stress that could negatively regulate neuronal survival and function. Changes in levels of the amino acids β-alanine, serine, alanine, valine, and glycine were increased in the brain of CP2-treated APP/PS1 mice, suggesting improved anabolism. Increased levels of 4-aminobutyrate, a metabolite involved in the gamma-aminobutyric acid (GABA) neurotransmitter system, indicate an improvement in neurotransmission and potential reversal of AD-related neurodegeneration. Increased levels of ascorbic/dehydroascorbic acids imply an improvement in vitamin C status and redox balance in the brain, consistent with CP2-induced activation of neuroprotective mechanisms (Extended Data Fig. 1 d-l).

### CP2 treatment reduces Aβ-related pathology, inflammation, and oxidative stress and improves proteostasis in brain and periphery

Histological examination of the hippocampus and cortex of CP2-treated APP/PS1 mice using 4G8 antibody revealed a significant reduction in Aβ plaques compared to untreated littermates (Fig. 2a,b). Biochemical analysis conducted using sequential extraction of brain tissue showed that CP2 treatment reduced total Aβ levels (Fig. 2c). While soluble Aβ was increased (Fig. 2d,e), levels of insoluble peptides were significantly decreased (Fig. 2f). Improved proteostasis could be promoted by autophagic degradation associated with AMPK activation and inhibition of the activity of glycogen synthase kinase-3 (GSK3β whose hyperactivation in AD is directly linked to Aβ pathology^28,29^. Indeed, CP2-dependent AMPK activation increased inhibitory phosphorylation of GSK3β and promoted the expression of proteins associated with lysosomal biogenesis and autophagy, including the transcription factor EB (TFEB), lysosomal-associated membrane protein 1 (LAMP-1), and microtubule-associated protein light chain 3 (LC3B) (Fig. 2g, Extended Data Fig. 6, Supplementary Fig. 3). These data further support the contention that CP2-induced autophagy is one of the neuroprotective pathways essential for Aβ clearance (Extended Data Fig. 1d).

**Fig. 2.**
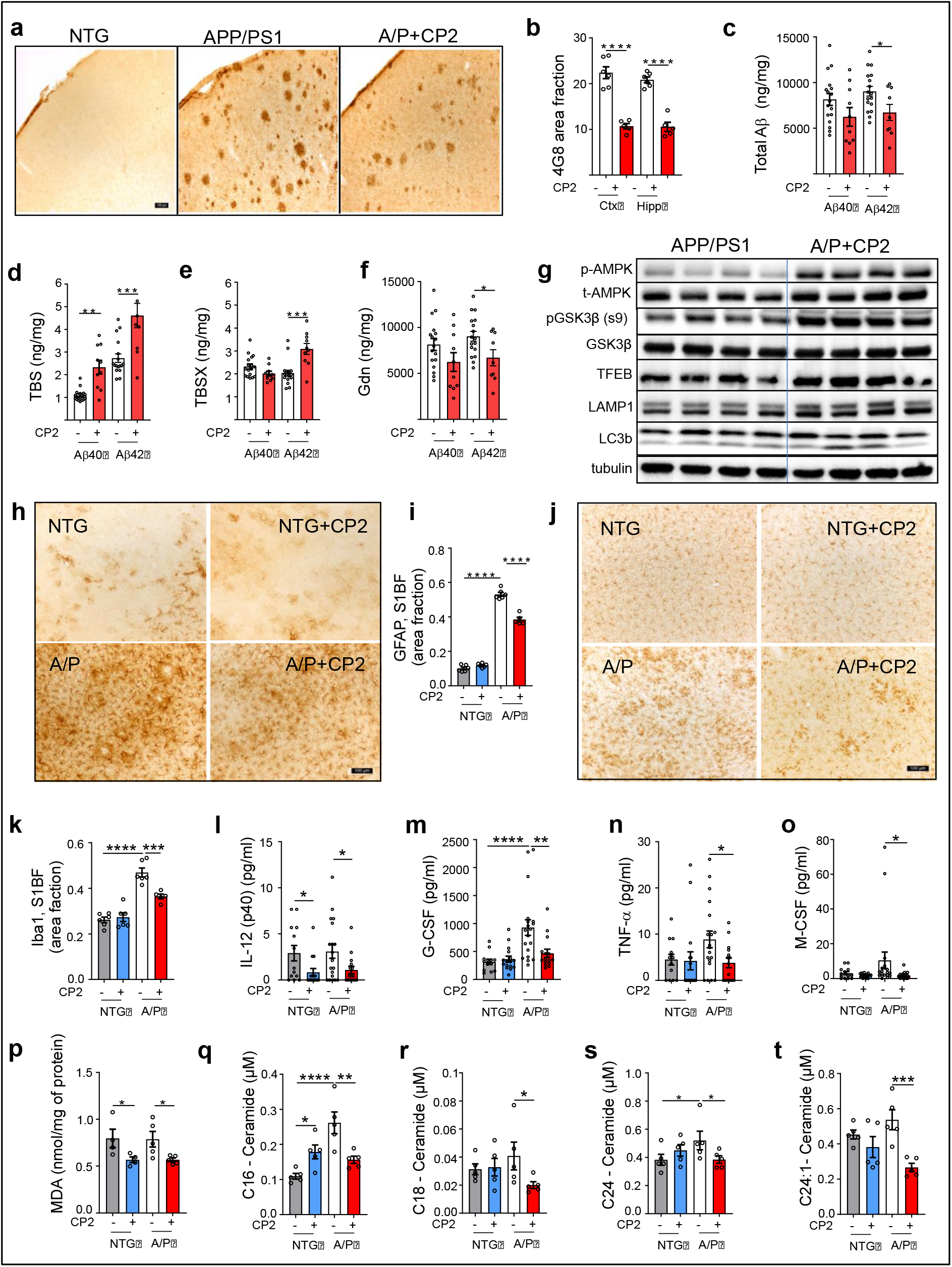
CP2 treatment reduces levels of Aβ, inflammation and oxidative stress in symptomatic APP/PS1 mice. **a,** Representative images of Aβ plaques visualized using 4G8 antibody in primary somatosensory barrel field (*S1BF*) cortex of NTG and APP/PS1 mice. Scale bar, 100 μm. **b**, Levels of Aβ plaques are significantly reduced in *S1BF* and hippocampus in CP2-treated APP/PS1 mice estimated using 4G8 antibody shown in (**a**). **c-f**, Differential centrifugation and ELISA revealed decreased levels of total Aβ42 in brain homogenates from CP2-treated APP/PS1 mice (**c**). Levels of soluble Aβ40 and 42 obtained using TBS (**d**) and TBSX (**e**) fractions were increased, while concentrations of the least soluble Aβ40 and 42 were decreased in brain fractions obtained using guanidine (Gdn) (**f**). *n* = 10 - 17 mice *per* group. **g**, CP2 treatment induces AMPK activation, reduces the activity of GSK3β, and activates autophagy in brain tissue of APP/PS1 mice (*n* = 6 - 8 mice *per* group). **h,j**, Representative images of GFAP (**h**) and Iba1 (**j**) staining in the *S1BF* in vehicle and CP2-treated NTG and APP/PS1 mice. Scale bar, 100 μm. **i,k**, Quantification of GFAP (**i**) and Iba1 (**k**) staining from (**h**) and (**j**), respectively. **l-o**, CP2 reduces pro-inflammatory markers in plasma of NTG and APP/PS1 mice. *n* = 15 - 20 mice *per* group. **p**, Levels of lipid peroxidation measured using malondialdehyde (MDA) were significantly reduced in brain tissue of CP2-treated NTG and APP/PS1 mice. *n* = 4 - 6 mice *per* group. **q-t**, Concentrations of ceramides (C16, C18, C24, C24-1) were significantly reduced in blood collected from CP2-treated APP/PS1 mice and measured using targeted metabolomics. *n* = 5 mice *per* group. All mice were 23-month-old. Data are presented as mean ± S.E.M. A two-way ANOVA with Fisher`s LSD *post-hoc* test was used for data analysis. For the comparison between vehicle and CP2-treated APP/PS1 groups (Fig. 2 b-f), an unpaired Student *t*-test was used for statistical analysis. *P < 0.05; **P < 0.01; ***P < 0.001; ****P < 0.0001.

Since AMPK activation is known to reduce inflammation and promote anti-oxidant response^30^, we examined levels of glial fibrillary acidic protein (GFAP) and the ionized calcium-binding adaptor molecule 1 (Iba1), a well-established markers of glial activation and inflammation, in the brain of CP2- and vehicle-treated APP/PS1 mice (Fig. 2h-k). Blood from the same mice was profiled for cytokines and chemokines (Fig. 2l-o, Extended Data Fig. 7a). We found that treatment significantly reduced inflammation in the brain and periphery, decreasing glial activation (Fig. 2h-k) and pro-inflammatory markers (*e.g*., IL-12, TNFα, G-CSF, Fig. 2l-o). We next measured lipid peroxidation in the brain, a well-established marker of oxidative stress prominent in the hippocampus and cortex of AD patients, which correlates with the extent of neurodegeneration and Aβ deposition^31^. After 13 months of CP2 treatment, levels of malondialdehyde (MDA) were significantly reduced in both APP/PS1 and NTG mice (Fig. 2p).

Oxidative stress and inflammation induce cell proliferation arrest and the cell senescence phenotype, contributing to age-related diseases. Therefore, we examined levels of senescent cells in inguinal (ING) and periovarian (POV) adipose tissue from NTG and APP/PS1 mice using β-Galactosidase (β-Gal) staining^32^ (Extended Data Fig. 7b-d). CP2 treatment significantly reduced abundance of senescent cells in both APP/PS1 and NTG mice. In AD patients, increased levels of ceramides, especially Cer16, Cer18, Cer20, and Cer24, were directly linked to oxidative stress and Aβ pathology^33^. Targeted metabolomic profiling conducted in plasma of CP2-treated mice revealed a significant decrease in concentrations of Cer16, Cer18, and Cer24, specifically in APP/PS1 mice (Fig. 2q-t). These data suggest that CP2 treatment induces multiple protective mechanisms including autophagy, anti-inflammatory and anti-oxidant responses, which contribute to improved proteostasis, reducing Aβ pathology, which in turn could be monitored using a translational metabolomic approach.

### CP2 treatment improves synaptic function, long term potentiation (LTP), dendritic spine maturation, and mitochondrial dynamics

Synaptic loss is the best correlate of cognitive dysfunction in AD^34^. To determine whether augmented cognitive performance after CP2 treatment was associated with improved synaptic function, we analyzed excitatory postsynaptic potential (fEPSP) in the CA1 region in acute hippocampal slices of APP/PS1 and NTG mice measuring local field potential (Fig. 3a-f)^35^. We initially recorded basal synaptic transmission and strength of post-synaptic responses to electrical stimulation of Schaffer collaterals (Fig. 3a). Activation of Schaffer collaterals revealed reduction of fiber volley amplitudes in APP/PS1 mice compared to NTG mice (Fig. 3b,c), which was partially restored by CP2 treatment (Fig. 3c).

**Fig. 3.**
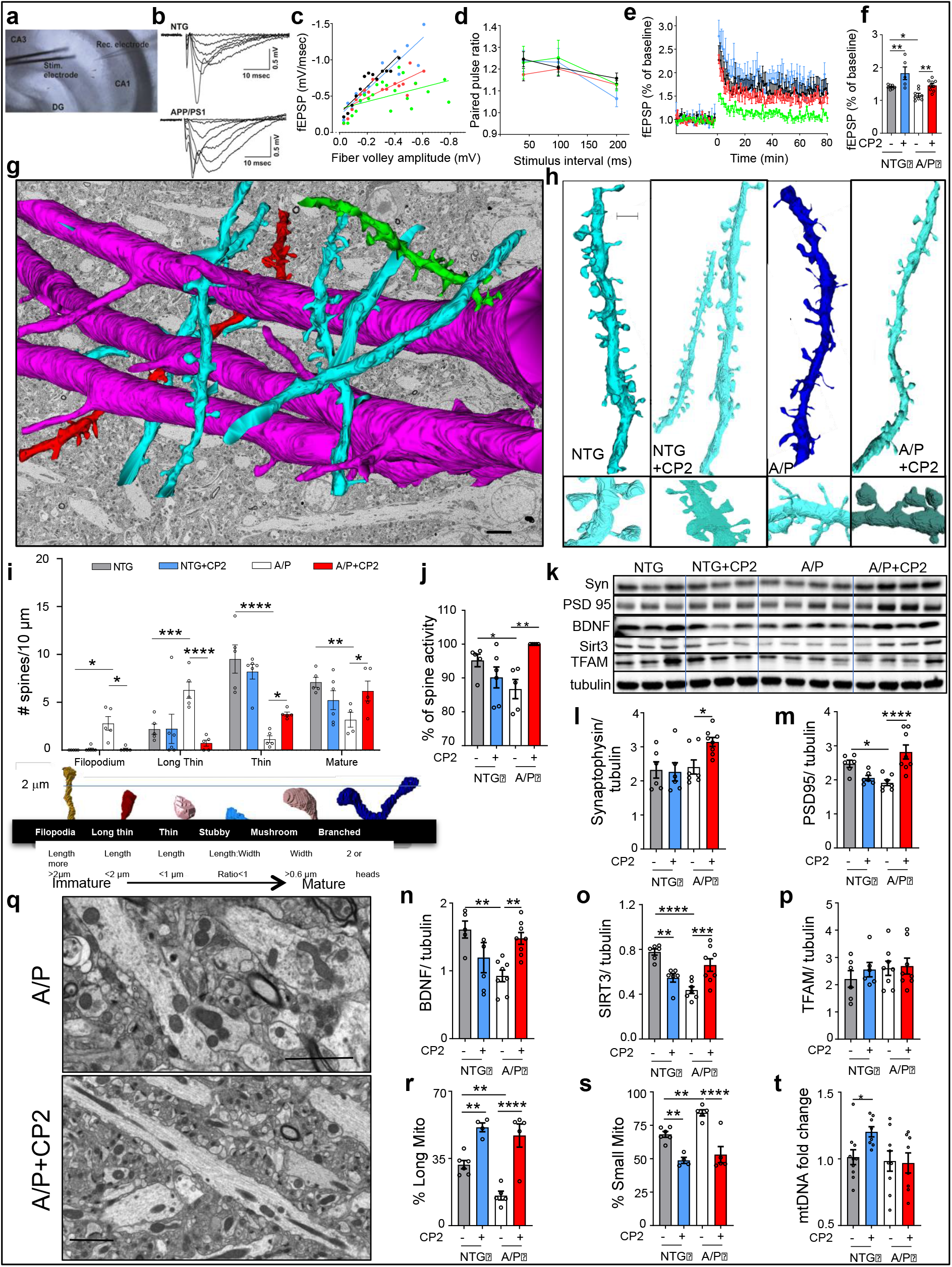
Chronic CP2 treatment improves synaptic activity, LTP, dendritic spine morphology, and mitochondrial dynamics in hippocampus of APP/PS1 mice. **a,** Experimental setting for stimulation-evoked local field potential (LFP) measurement in the hippocampal slice. The stimulation (Stim) electrode was placed at the Schaffer collaterals and the recording (Rec) electrode was placed at the *striatum radiatum* in the CA1. **b**, Representative raw traces of field excitatory post-synaptic potentials (fEPSP) from NTG and APP/PS1 mice. Various stimulation intensities (10 - 300 μA) were applied to evoke fEPSP. The stimulation pulse width and intervals were fixed at 60 μsec and 30 sec, respectively. As the stimulation intensity was increased, the initial slope of fEPSP was increased. **c**, CP2 effect on basal synaptic strength. To examine the pre-post synaptic relationships, initial slopes of fEPSP were plotted against amplitudes of presynaptic fiber volleys. Pre-post synaptic relationship in the CP2-treated APP/PS1 group was improved compared to the APP/PS1 group. **d**, Paired-pulse facilitation did not differ between experimental groups. Two stimulations were applied with a short interval to determine presynaptic involvement in synaptic plasticity. **e**, CP2 treatment improves LTP formation. Average traces for fEPSPs in hippocampal slices from each experimental group (*n* = 2-3 slices from 3–5 mice *per* group). Traces represent mean ± S.E.M. *per* time point. To induce LTP, three tetanic stimulations (100 Hz, 60 μsec-pulse width for 1 sec) were applied with 3-second intervals. In APP/PS1 hippocampus, the tetanic stimulation induced early phase post-tetanic potentiation; however, long lasting potentiation was not observed. In the slice from APP/PS1 CP2-treated mice, LTP was induced and maintained over 60 minutes. NTG, black; APP/PS1, green; NTG+CP2, blue; APP/PS1+CP2, red. **f**, LTP intensities among groups were compared at 60 min (*n* = 2-3 slices from 3–5 mice *per* group). **g**, 3DEM reconstruction of axons (red) and dendrites (green, blue) from the CA1 hippocampal region of an APP/PS1 mouse. Reconstruction is superimposed on 2DEM from the same brain. Scale bar, 5 μm. **h**, Representative 3DEM reconstructions of dendrites from CA1 region of NTG and APP/PS1 mice. Scale bar, 1 μm. **i**, Quantification of dendritic spine morphology in vehicle and CP2-treated NTG and APP/PS1 mice. Mature dendritic spines included stubby, mushroom and branched. **j**, Quantification of active synapses visualized using 3DEM. **k**, Western blot analysis in the hippocampal tissue assaying levels of synaptophysin (Syn), BDNF, post synaptic density 95 (PSD95), TFAM, and Sirt3. l**-p**, Quantification of proteins from (**k**). **q**, Representative 2DEM micrographs of mitochondria in the hippocampus of CP2- and vehicle-treated APP/PS1 mice are used for quantifying mitochondrial morphology. **r,s**, CP2 treatment increases the number of elongated mitochondria and decreases the number of small organelles. **t**, CP2 increased mitochondrial DNA copy number in NTG mice. Data are presented as mean ± S.E.M. A two-way ANOVA with Fisher`s LSD *post-hoc* test was used. *n* = 5 *per* group. *P < 0.05, **P < 0.01, ***P < 0.001, ****P < 0.0001.

Since short-term plasticity plays a crucial role in neuronal information processing relevant to cognitive function, we next investigated the effect of CP2 on Schaffer collaterals-CA1 short-term plasticity utilizing a paired-pulse stimulation protocol. Paired-pulse facilitation (PPF) measures the ability of synapses to increase transmitter release upon the second of two closely spaced afferent stimuli, which depends on residual calcium levels in the presynaptic terminal^36^. If LTP is mediated presynaptically, an increase in transmitter release is accompanied by a change in short-term plasticity. We found that the PPF was not different between groups (Fig. 3d), suggesting that CP2-dependent improvement in LTP in APP/PS1 mice was not associated with the pre-synaptic release of neurotransmitters. We further applied tetanic stimulation to Schaffer collaterals-CA1 to induce and record LTP over 60 minutes to determine EPSP. Significant currents associated with strong LTP were recorded in NTG and CP2-treated NTG and APP/PS1 mice, while vehicle-treated APP/PS1 mice did not exhibit significant LTP formation (Fig. 3e,f). These data indicate that cognitive impairment in APP/PS1 mice could be associated with the inability to form and maintain LTP in the hippocampus, while CP2 treatement corrected this defect.

LTP critically depends on the morphology of dendritic spines, which determines synaptic strength and plasticity^37,38^. We examined dendritic spine morphology in the CA1 hippocampal region of vehicle- and CP2-treated APP/PS1 and NTG mice using three-dimensional electron microscopy (3D EM) reconstruction (Fig. 3g,h)^39^. In NTG mice, the majority of spines were mature (thin, stubby, mushroom, and branched), while immature filopodia and long thin spines were prevalent in APP/PS1 mice (Fig. 3h,i). CP2 treatment promoted maturation of dendritic spines in APP/PS1 mice (Fig. 3h,i) and markedly improved spine geometry (the length and width of spine necks and heads, and compartmentalization factor) in NTG and APP/PS1 mice, indicating greater ability to maintain LTP^40^ (Extended Data Fig. 8). In APP/PS1 mice, CP2 treatment restored the mushroom spine volume, length, and head width to the dimensions observed in NTG mice (Extended Data Fig. 8a-e). In NTG and APP/PS1 mice, CP2 significantly increased the compartmentalization factor (Extended Data Fig. 8f), a measure of the spine head depolarization during synaptic transmission, which is regulated by the length of the spine neck and indicates greater synaptic plasticity^41^. Increased spine maturation resulted in more active synapses in CP2-treated APP/PS1 mice, bringing synaptic activity to the level of NTG mice (Fig. 3j). Improved synaptic function in CP2-treated APP/PS1 mice was associated with increased levels of synaptophysin, the postsynaptic density protein PSD95, and BDNF (Fig. 3k-n, Supplementary Fig. 4). Thus, CP2-dependent cognitive protection is associated with improved morphology of dendritic spines, LTP, and synaptic transmission in the hippocampus.

Since synaptic activity requires energy, we examined mitochondrial integrity in the brain tissue of the same APP/PS1 and NTG mice utilized in the study of dendritic spines (Fig. 3q-t). Consistent with reports on mitochondrial fragmentation in AD, we observed increased numbers of round-shaped organelles in vehicle-treated APP/PS1 mice. CP2 treatment resulted in a higher number of elongated organelles and increased levels of neuroprotective Sirt3 in APP/PS1 mice (Fig. 3k,o,q,r), which could additionally contribute to reduced inflammation and neuroprotection^19,42,43^. Interestingly, the increased mitochondrial mass was observed only in CP2-treated NTG mice (Fig. 3t). Together with our previous reports of CP2-dependent restoration of axonal trafficking and enhanced bioenergetics^12^, these data demonstrate an improvement in mitochondrial dynamics and function in symptomatic APP/PS1 mice, which is essential for synaptic function and improved energy homeostasis

### CP2 treatment attenuates the ongoing neurodegeneration

Neurons of the *locus coeruleus* (LC) provide norepinephrine to the hippocampus, mediating memory and attention^44^. AD patients exhibit early neurodegeneration in the LC where severity of neuronal loss correlates with the duration of illness^45^. Neurodegeneration in the LC has been shown to affect Aβ and Tau aggregation, inflammation, synaptic function, neuronal metabolism, and the BBB permeability^46^. Previously, we demonstrated that the degeneration of LC neurons in human AD is recapitulated in mouse models of cerebral amyloid^47,48^. In the current APP/PS1 mice, progressive loss of TH+ cortical afferents starts at 6 months of age (Fig. 4a,b; Extended Data Fig. 9a,c), followed by the loss of noradrenergic (TH^+^) neurons in the LC (Fig. 4c,d, Extended Data Fig. 9b,d). Consistent with the progressive loss of afferents, there was a reduction in the volume of TH^+^ neurons in APP/PS1 mice starting at 12 months of age (Fig. 4e,f). Analysis of 12 month old APP/PS1 mice that received CP2 treatment for 2 months (from 10 months of age) showed that CP2 did not impact neurodegeneration, since the cortical TH^+^ axon density, the number of TH^+^ neurons, and neuronal volumes were similar between vehicle- and CP2-treated subjects (Fig. 4b,d,f; 12-month-old group). In mice receiving CP2 for 10 months, further progression of neurodegeneration was completely halted by the CP2 treatment. Thus, cortical TH^+^ axon density, TH^+^ neuron number in LC, and TH^+^ neuronal volume in 20 month old CP2-treated APP/PS1 mice were comparable to those in 12 month old APP/PS1 mice (Fig, 4b,d,f; 20-month-old group). These data demonstrate that CP2 specifically protects the neuronal network in APP/PS1 mice, which might be associated with the reduction of Aβ accumulation and toxicity.

**Fig. 4.**
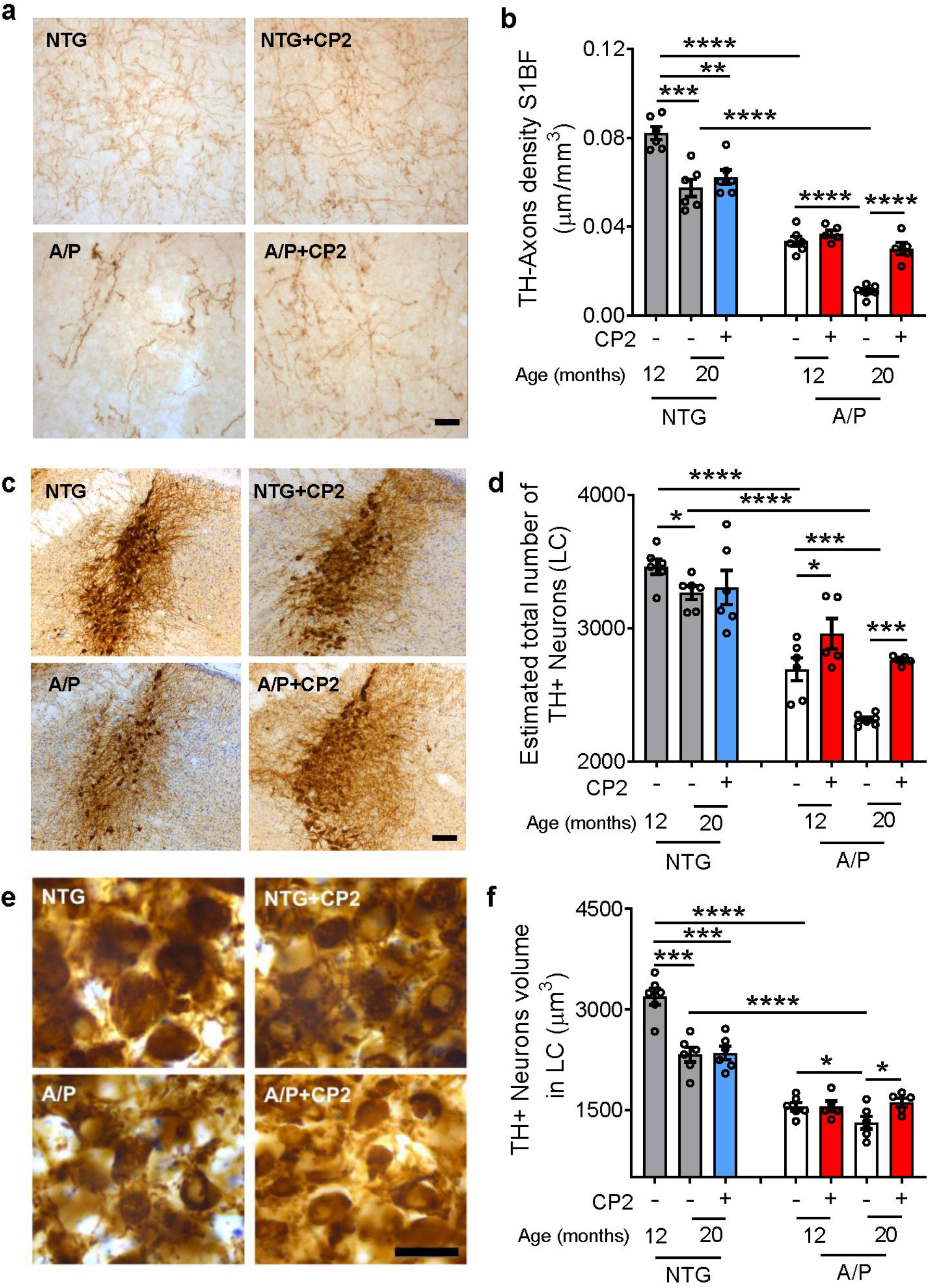
CP2 treatment after the onset of AD-like neuropathology halts progressive degeneration of LC neurons in APP/PS1 mice. APP/PS1 (A/P) and NTG mice were treated with CP2 or vehicle starting at 10 months of age. The animal brains were harvested at 12 (2 months treatment) or 20 months of age (10 months treatment) and evaluated for the integrity of the NAergic neurotransmitter system. **a**, CP2 stops the progressive loss of TH^+^ axons in the cortex of 20-month-old mice A/P mice. Representative images of TH^+^ axonal projections in Sensory Barrel Cortex (S1BF). Scale bar, 20 μm. **b**, The density (μm/mm^3^) of TH^+^ axons in S1BF was determined using stereological length estimation using spherical probe and images presented in (**a**). Compared to NTG mice, A/P mice exhibit a significant progressive loss of TH^+^ axons at 12 and 20 months of age. CP2 treatment prevented loss of TH^+^ axons in APP/PS1 mice occurs between 12 and 20 months of age. **c**, Representative image of TH^+^ LC neurons in 20-month-old mice. Scale bar, 100 μm. **d**, CP2 stops the progressive loss of TH^+^ neurons in A/P mice in LC. **e**, Higher magnification images from (**c**) were used to evaluate the relative sizes of neurons. Scale bar, 50 μm. **f**, CP2 stops the progressive loss of TH^+^ neuronal volume in A/P mice at 20 months of age. *n* = 5 – 7 female mice *per* group. Data are presented as mean ± S.E.M. A two-way ANOVA with Fisher`s LSD *post-hoc* test was used to analyze the differences between A/P mice, and between untreated groups of NTG and A/P mice. A Student *t*-test was used to analyze the differences between untreated and CP2-treated NTG mice. *P < 0.05, **P < 0.01, ***P < 0.001, ****P < 0.0001.

### CP2 treatment activates translational neuroprotective mechanisms essential for human AD

To investigate further mechanisms associated with CP2 efficacy, we performed next-generation RNA sequencing (RNA-seq) using brain tissue from vehicle- or CP2-treated NTG and APP/PS1 mice (Fig. 5). Principal component analysis (PCA) revealed a good separation among all groups (Extended Data Fig. 10a). The comparison between vehicle-treated NTG and APP/PS1 mice identified 3320 differentially expressed genes (DEGs) (Fig. 5a, Supplementary Table 6). The top functional changes associated with these DEGs are listed in Supplementary Tables 7,8 and Extended Data Fig. 10b. Processes affected by the disease in APP/PS1 mice overlap with pathways well-established in AD patients including ATP metabolism, ion transport, nervous system development, synaptic transmission, and inflammation^49–54^ (Extended Data Fig. 10b). Comparison of CP2- and vehicle-treated APP/PS1 mice revealed changes in 1262 DEGs (Fig. 5a, Supplementary Table 9). The top biological processes associated with these DEGs included inflammatory response, redox signaling, nervous system development, and regulation of axonal guidance (Extended Data Fig. 10c-e, Supplementary Table 10,11). Out of 3320 genes differentially affected in vehicle-treated APP/PS1 *vs.* NTG mice, the expression of 567 genes was reverted by CP2 treatment to the levels detected in NTG mice (Fig. 5b, Supplementary Table 12). A heatmap of changes in these 567 DEGs shows two clusters with a subset of genes that were either down- or up-regulated by CP2 treatment (Fig. 5b, Clusters 1 and 2, Supplementary Tables 13,14). Gene function enrichment analysis showed that pathways down-regulated by CP2 in APP/PS1 mice included oxidative stress and immune response (Fig. 5b-d), consistent with data generated in the brain and periphery of CP2-treated APP/PS1 mice (Fig. 2).

**Fig. 5.**
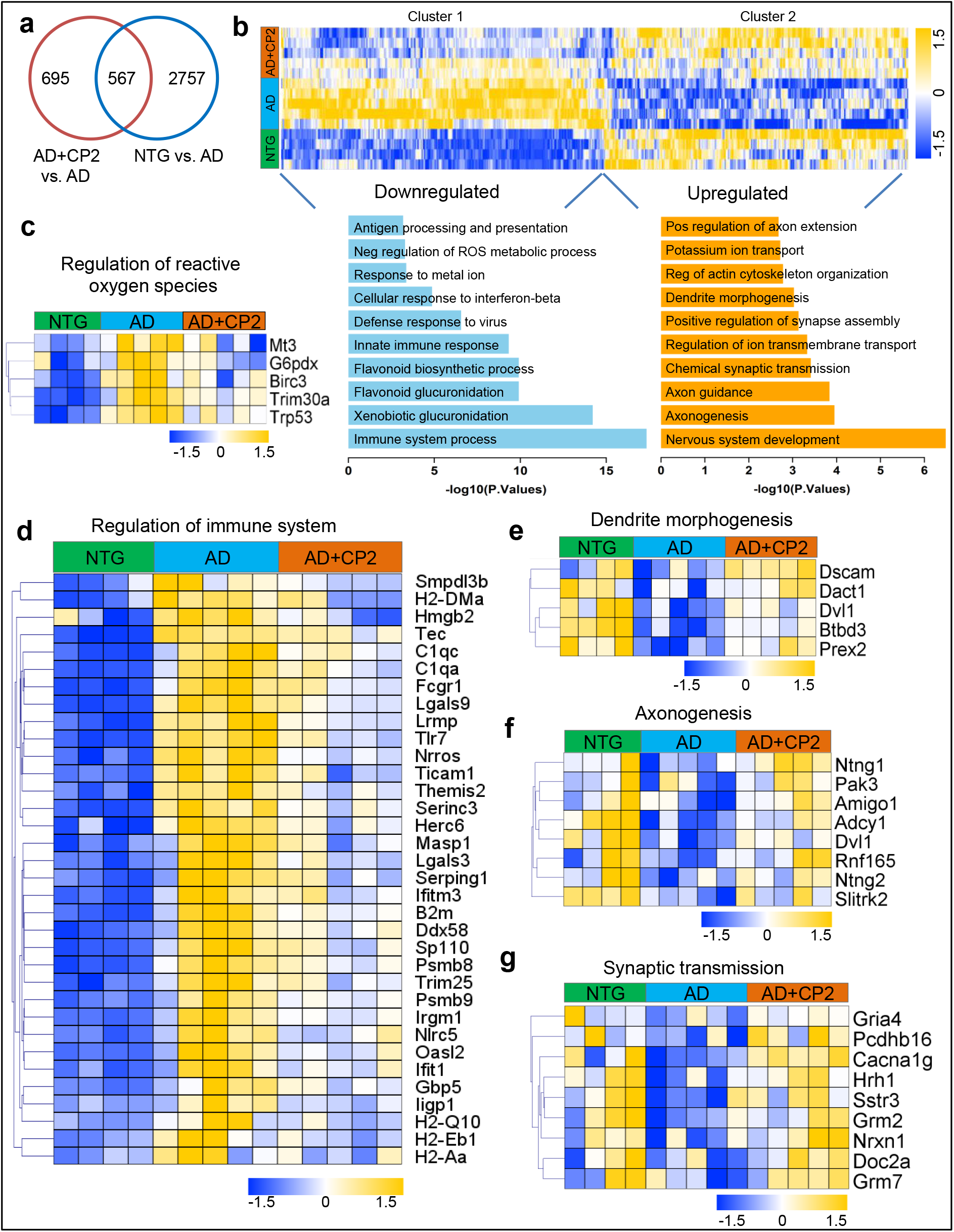
Global gene expression patterns in brain tissue of APP/PS1 mice treated with CP2 relative to NTG and APP/PS1 mice. **a,** Venn diagram of differentially expressed genes (P < 0.05) in the brain tissue of vehicle- and CP2-treated NTG and APP/PS1 mice. Overlapped DEGs (567) represent specific gene pools affected by CP2 treatment. **b**, A heatmap of the overlapped 567 genes shows two clusters where CP2 treatment reversed (up- or down-regulated) the expression of a subset of genes in APP/PS1 mice to the levels observed in NTG littermates. Gene function enrichment analysis shows pathways associated with down-regulated (Cluster 1) or up-regulated (Cluster 2) genes after CP2 treatment in APP/PS1 mice. **c, d**, Heatmaps of changes in genes associated with reactive oxygen species (**c**) and the immune system (**d**) that were down-regulated after CP2 treatment in APP/PS1 mice. **e-g**, Heatmaps of changes in genes associated with dendrite morphogenesis (**e**), axonogenesis (**f**), and synaptic transmission (**g**) that were up-regulated after CP2 treatment in APP/PS1 mice. The weight of the edges corresponds to the confidence scores of gene integration. All mice were 23-month-old treated with CP2 or vehicle for 13-14 months. *n* = 4 - 5 mice *per* group.

Among genes involved in the regulation of oxidative stress and apoptosis were G6pdx, BIRC3, TRIM30a, Trp53, and Mt3, all known to play a role in human disease (Fig. 5c). Other significant changes included global down-regulation of the immune response by CP2 including the acute phase response, such as up-regulation of SERPING1, interferon signaling (DDX58, FCGR1A, TRIM25, GBP5, TLR7, IFITM3, IFIT1, NLRC5, OASL2, IRGM1, IIGP1), and major histocompatibility complex (MHC) class II presentation (H2-DMa, H2-Q10, H2-Eb1, H2-Aa, PSMB8, PSMB9) (Fig. 5d). Pathways up-regulated by CP2 included dendritic spine maturation, axonal extension and guidance, and synaptic transmission (Fig. 5b,e-g). The identified up-regulated genes in dendrite morphogenesis pathways included the Down syndrome cell-adhesion molecule (DSCAM) (Fig. 5e), which is involved in governing neurite arborization, mosaic tiling, and dendrite self-avoidance and BTB Domain Containing 3 (BTBD3), which has a role in dendritic guidance toward active axon terminals. Genes that mediate axonogenesis, including Ntng1 and Ntng2, and axonal guidance, were also up-regulated in CP2-treated APP/PS1 mice (Fig. 5f, Extended Data Fig. 10d). Additional genes up-regulated by CP2 in APP/PS1 mice included those involved in synaptic transmission and synapse assembly and that are known to be down-regulated in AD patients, including glutamate receptor 4 (GRIA4), which mediates fast synaptic excitatory neurotransmission; metabotropic glutamate receptor 7 and 21 (GRMN7 and GRM2), which facilitates the formation of LTP; Double C2 protein (Doc2a), which contributes to spontaneous excitatory and inhibitory release; and Neurexins1 (NRXN1), which facilitates formation of functional synaptic structures (Fig. 5g, Extended Data Fig. 10e). These data are consistent with the improved synaptic function in CP2-treated APP/PS1 mice (Fig. 3).

To provide further evidence for the translational potential of our findings, we cross-validated transcriptomic data from our study with the human brain transcriptome available through coexpression meta-analysis across the Accelerating Medicines Partnership in Alzheimer’s Disease Target Discovery and Preclinical Validation Project (AMP-AD – ampadportal.org)^55^. RNA-seq AMP-AD data were generated across three large scale but distinct human *postmortem* brain studies collected from 2114 samples across 7 brain regions and 3 research studies^52–54^.

Since in our study we used only female mice, we restricted human data to females only. We first correlated genetic changes found in our comparison of NTG *vs*. APP/PS1 mice to significant DEGs identified in comparison between the AD *vs*. control female cohort in the AMP-AD set. Out of 3114 down-regulated DEGs in the human AD cohort (Fig. 6a, Supplementary Table 15), we identified 294 mouse DEGs that matched the human gene set (Supplementary Table 16). Functional enrichment analysis showed that the most down-regulated pathways in both human and mouse AD were involved in synaptic transmission, nervous system development, histone deacetylation, and axonogenesis (Fig. 6b, Supplementary Table 17). Down-regulated shared genes in mouse and human AD included genes involved in pyruvate metabolism such as MCP2, DLAT and PDHB (Supplementary Table 17). Among 518 up-regulated DEGs shared between human and mouse AD, top functions were enriched for the innate immune response (Fig. 6a,c, Supplementary Tables 18,19). Thus, APP/PS1 mice recapitulate major pathways affected in human AD.

**Fig. 6.**
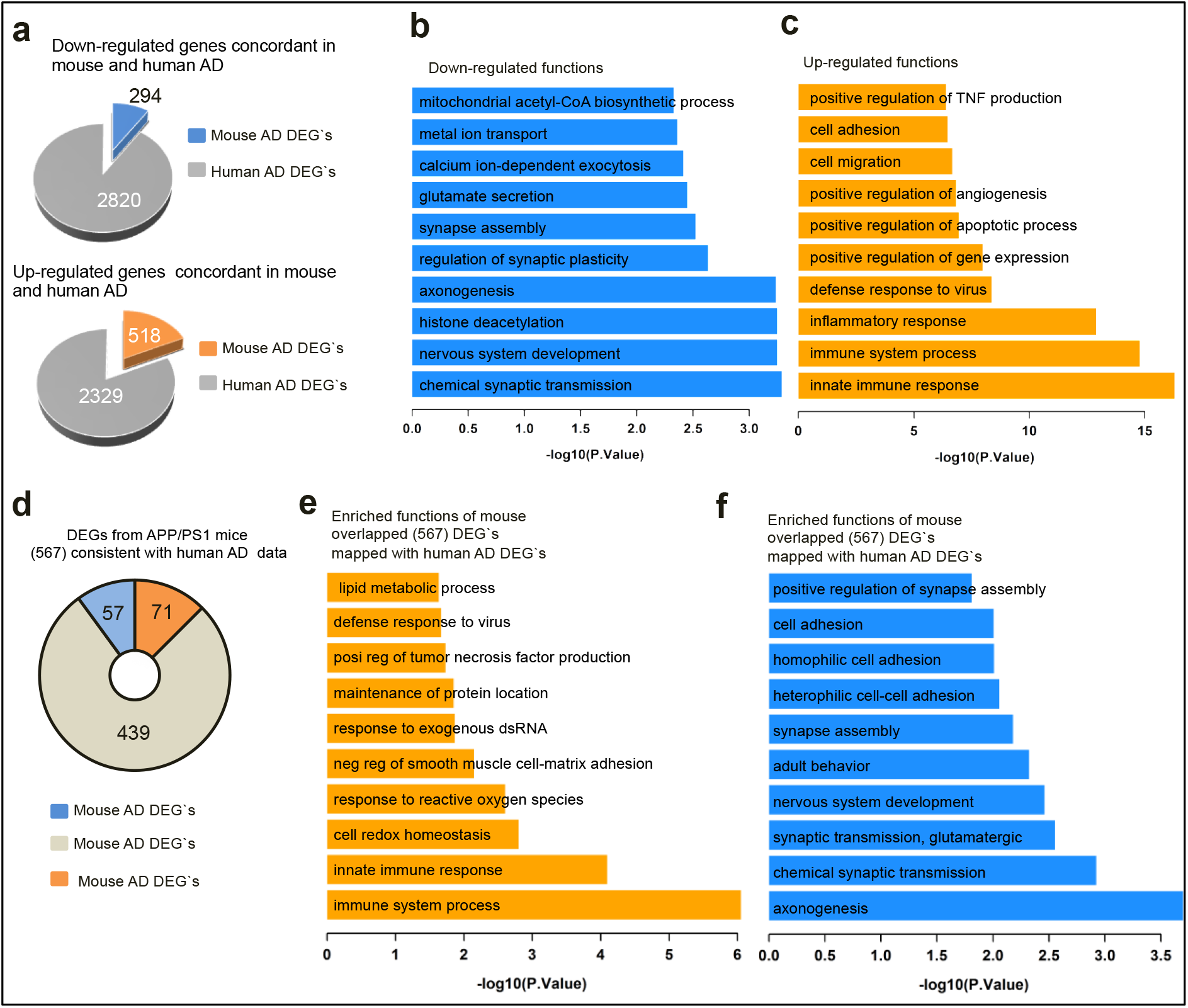
CP2 treatment affects pathways relevant to AD patients based on the cross-validation of mouse and human AMP-AD transcriptomic data. **a,** Pie diagrams showing the number of down-regulated (upper graph) and up-regulated (lower graph) DEGs in vehicle-treated APP/PS1 mice that correlate with corresponding down- or up-regulated DEGs in the human AMP-AD dataset, respectively. **b**. Enriched functions of down-regulated DEGs in vehicle-treated APP/PS1 mice that correlate with the identified down-regulated DEGs in the human AMP-AD dataset. **c**. Enriched functions of up-regulated DEGs in vehicle-treated APP/PS1 mice that correlate with the identified up-regulated DEGs in the human AMP-AD dataset. **d**. Comparison of human DEGs from the AMP-AD set with 567 DEGs shown in Fig. 5a that are specifically affected by CP2 treatment in APP/PS1 mice. **e**, Functions identified by the enrichment analysis associated with the up-regulated DEGs (567) in APP/PS1 mice mapped against human up-regulated AD DEGs. These functions were reversed (down-regulated) by CP2 treatment in APP/PS1 mice. **f**, Functions identified by the enrichment analysis associated with the down-regulated DEGs (567) in APP/PS1 mice mapped with human down-regulated AD DEGs. These functions were reversed (up-regulated) by CP2 treatment in APP/PS1 mice. All mice were 23-month-old treated with CP2 or vehicle for 13-14 months. *n* = 4 - 5 mice *per* group.

We next compared the 567 DEGs associated with CP2 treatment in APP/PS1 mice (Fig. 5a) with DEGs from females in the AMP-AD RNA-seq data collection. We found that 128 out of the total 567 overlapping mouse AD DEGs corresponded to human AD genes (Fig. 6d). CP2 treatment in APP/PS1 mice reversed the expression of 71 genes that were up-regulated in both mouse and human AD (Supplementary Table 20). Functional enrichment analysis showed that these 71 genes were involved in the regulation of the immune processes, inflammation, response to reactive oxygen species, and TNF production (Fig. 6e, Supplementary Table 21). In contrast, CP2 reversed the expression of 57 deregulated genes that are involved in axonogenesis, glutamatergic synaptic transmission, nervous system development, and synapse assembly (Fig. 6d,f and Supplementary Tables 22,23). Taken together, these data demonstrate that AD-associated transcriptional and functional changes observed in our mouse model of AD are counteracted by CP2 treatment. CP2-treated APP/PS1 mice showed attenuated expression of a significant number of genes involved in neuroinflammatory processes, consistent with our observation of a decreased number of activated astrocytes and microglia. Moreover, CP2 treatment of APP/PS1 mice restored expression of genes involved in neurotransmission, dendritic morphology and axonal guidance and extension, which were linked to the restoration of hippocampal LTP and increased number of mature dendritic spines (Fig. 3). These data demonstrate that pathways improved by CP2 treatment in APP/PS1 mice comprise major pathways essential for therapeutic efficacy in AD patients.

## Discussion

AD is associated with early energy hypometabolism, synaptic and mitochondrial dysfunction, oxidative stress, inflammation, abnormal proteostasis and progressive neurodegeneration. Here, we demonstrate that mild energetic stress associated with partial inhibition of MCI induces activation of integrated stress-response mechanisms that attenuate effects of pathological pathways such as abnormal energy homeostasis, synaptic dysfunction, and inflammation, ultimately blocking neurodegeneration in a translational mouse model of AD, the APP/PS1 mouse. The therapeutic efficacy achieved has translational relevance, as the intervention was started after the onset of Aβ neuropathology^24^, cognitive symptoms^25^, bioenergetic dysfunction^26^, and progressive neurodegeneration. Beneficial mechanisms affected by CP2 treatment in APP/PS1 mice overlap with signatures established in AD patients, females in particular, supporting the high translational potential of this approach. Major translational targets affected by CP2 treatment included the immune system response and multiple pathways involved in synaptic function and neurotransmission, which are underlie early pathology in AD patients^7^. Since CP2 improved axonogenesis and dendritic spine morphology and function, it is feasible that this treatment could also induce neuronal regeneration.

The strength of our study is in the utilization of multiple early and late disease outcome measures in a mouse model of AD that closely mimics neuropathological mechanisms of human disease. In particular, we hypothesized that the general failure of the preclinical studies in mouse models of AD to predict outcomes of human clinical trials is related to the reliance on treatments of younger mice and a very limited set of neuropathological outcome measures. Thus, it is important that CP2 treatment of APP/PS1 mice was conducted when AD-like pathology, including progressive neurodegeneration, was well established and the broad array of measures, including advanced imaging techniques and translational biomarkers, were applied *in vivo* and in tissue, further supporting the ability to monitor therapeutic efficacy of this approach in humans.

While the details and the hierarchy of molecular mechanisms involved in neuroprotective stress response require further evaluation, AMPK activation appears to play a central role. Indirect activation of AMPK by inhibition of MCI has been shown to increase life span, rejuvenate the transcriptome, and protect from neurodegeneration^56–60^. Paradoxically, these manipulations improved MCI assembly, increased complex I-linked state 3 respiration, and decreased ROS production^57^. Studies in a cohort of 2200 ultranonagenarians revealed that mutations in subunits of MCI that resulted in its partial inhibition had beneficial effect on longevity^61^. The compelling support for safety of the application of MCI inhibitors in humans comes from metformin, an FDA approved drug to treat diabetes. Among other targets, metformin inhibits MCI. It is prescribed to the elderly population and has relatively safe profile even after chronic treatment^62^. Recent study conducted in a large Finish population of older people with diabetes demonstrated that long-term and high-dose metformin use do not increase incidences of AD and is associated with a lower risk of developing AD^63^. Resveratrol is another MCI modulator where its effect on MCI (activation or inhibition) depends on the concentration. It also inhibits mitochondrial Complex V. Resveratrol is currently in clinical trials for multiple human conditions^64–66^. Compared to CP2, these compounds have limitations associated with the lack of selectivity, specificity, and bioavailability.

CP2 treatment effectively reduced Aβ accumulation. This might be explained by an additional ability of CP2 to bind Aβ peptides reducing their toxicity, which together with the AMPK-dependent mechanisms, could produce a synergistic effect^67^. Our work also provides evidence that CP2 treatment improves mitochondrial dynamics and function. While counterintuitive, given that progressive mitochondrial dysfunction is well characterized in AD patients and APP/PS1 mice, mild MCI inhibition in dysfunctional mitochondria could decrease ROS production, while in functional mitochondria could improve energetics, as we demonstrated previously^67^, and through activation of stress responses, could promote biogenesis and mitophagy, contributing to a healthier mitochondrial pool and more effective energy production. Positive effects on mitochondrial function and neuroprotection are further supported by an increase in Sirt3 levels. In AD patients and AD models, reduction of Sirt3 was directly linked to the loss of synaptic function, Aβ and Tau pathology, and neurodegeneration^19,20,42^. While APP/PS1 mice utilized in our study do not have pTau accumulation, increased levels of Sirt3 and decreased activity of GSK3β suggest that CP2 treatment could also be effective in reducing Tau toxicity. While not as pronounced as in APP/PS1 mice, CP2 treatment improved health parameters in aged NTG mice, reducing oxidative stress and cellular senescence, key pathways that accelerate aging, and improving body mass, glucose tolerance, mitochondrial function, and physical strength. Taken together, the results of our studies suggest that partial inhibition of MCI represents a novel treatment strategy for blocking neurodegeneration and cognitive impairment, even after the development of AD-like symptoms, and also could enhance healthspan delaying the onset of multiple age-related diseases, including AD.

## Methods

All experiments with mice were approved by the Mayo Clinic Institutional Animal Care and Use Committee in accordance with the National Institutes of Health’s *Guide for the Care and Use of Laboratory Animals*.

### CP2 synthesis

CP2 was synthesized by the Nanosyn, Inc biotech company (http://www.nanosyn.com) as described^68^ and purified using HPLC. Authentication was through NMR spectra to ensure lack of batch-to-batch variation in purity. Standard operating procedures (SOP) were developed and followed for CP2 preparation, storage, cell treatment, and administration to animals.

### Mice

The following female mice were used in the study: double transgenic APP/PS1^69^ and their non-transgenic (NTG) littermates; and C57BL/6J wild-type mice. Genotypes were determined by PCR as described in^69^. All animals were kept on a 12 h–12 h light-dark cycle, with a regular feeding and cage-cleaning schedule. Mice were randomly selected to study groups based on their age and genotype. Mice were housed 5 *per* cage, water consumption and weight were monitored weekly. CP2 concentration was adjusted based on mouse weight/water consumption weekly. The number of mice in each group was determined based on the 95% of chance to detect changes in 30 - 50% of animals. The following exclusion criteria were established: significant (15%) weight loss, changing in the grooming habits (hair loss), pronounced motor dysfunction (paralyses), or other visible signs of distress (unhealed wounds).

### Human brain tissue

Experiments with *post-mortem* human brain tissue were approved by the Mayo Clinic IRB (#12-007847) and were carried out in accordance with the approved guidelines. Informed consent was obtained from all subjects involved in the study. One brain specimen with a postmortem interval of 6 h from a cognitively normal female 99 years old was obtained from Mayo Clinic. Cortex was used for mitochondrial isolation to determine CP2-dependent inhibition of MCI.

### Mitochondrial isolation and measurements of ETC complex activity

Intact brain mitochondria were isolated from *postmortem* human or mouse brain tissue using differential centrifugation with digitonin treatment^70^. Brain tissue was immersed into ice-cold isolation medium (225 mM mannitol, 75 mM sucrose, 20 mM HEPES-Tris, 1mM EGTA, pH 7.4), supplemented with 1 mg/ml BSA. Tissue was homogenized with 40 strokes by pestle “B” (tight) of a Dounce homogenizer in 10 ml of isolation medium, diluted two-fold, and transferred into centrifuge tubes. The homogenate was centrifuged at 5,900 g for 4 minutes in a refrigerated (4 °C) Beckman centrifuge. The supernatant was centrifuged at 12,000 g for 10 minutes and pellets were resuspended in the same buffer, and 0.02% digitonin was added. The suspension was homogenized briefly with five strokes in a loosely fitted Potter homogenizer and centrifuged again at 12,000 g for 10 minutes, then gently resuspended in the isolation buffer without BSA and washed once by centrifuging at 12,000 g for 10 minutes. The final mitochondrial pellet was resuspended in 0.1 ml of washing buffer and stored on ice. The respiratory activities were measured in Oroboros high-resolution respirometer as previously described^71^.

The activity of ETC complexes was measured spectrophotometrically using a plate reader (SpectraMax M5, Molecular Devices, USA) in 0.2 ml of standard respiration buffer composed of 125 mM sucrose, 25 mM Tris-HCl (pH=7.5), 0.01 mM EGTA, and 20 mM KCl at 25 °C. NADH-dependent activity of complex I was assayed as oxidation of 0.15 mM NADH at 340 nm (ε340 nm = 6.22 mM^−1^cm^−1^) in the assay buffer supplemented with 10 μM cytochrome *c*, 40 μg/ml alamethicin, 1 mM MgCl_2_ (NADH media). NADH:Q reductase was measured in NADH media containing 2 mg/ml BSA, 60 μM decylubiquinone, 1 mM cyanide and 5-15 μg protein *per* well. Only the rotenone (1 μM)-sensitive part of the activity was used for calculations. NADH:HAR reductase was assayed in NADH media containing 1 mM HAR and 2-5 μg protein *per* well. Complex II succinate:DCIP reductase activity was recorded at 600 nm (ε600nm = 21 mM^−1^cm^−1^) in the KCl assay buffer (125 mM KCl, 20 mM HEPES-Tris, 0.02 mM EGTA, pH 7.6) containing 15 mM succinate, 40 μM decylubiquinone, 0.1 mM DCIP, 1 mM KCN, and 5-10 μg protein *per* well. Complex IV ferrocytochrome *c* oxidase activity was measured as oxidation of 50 μM ferrocytochrome *c* at 550 nm (ε550 nm = 21.5 L∙mM^−1^cm^−1^) in KCl assay buffer supplemented with 0.025% dodecylmaltoside and 1-3 μg protein *per* well. To assess the effect of CP2 on the activity, we pre-incubated 20-40 μg/ml mitochondria with various concentrations of CP2 for 10 min at 25 °C in the absence of substrates and then measured the residual activity as described above.

### *In vitro* Safety Pharmacology Studies

The *in vitro* Safety Pharmacology assays including CP2 binding and enzyme inhibition and uptake measurements, were conducted by the Contract Research Organization (CRO) Eurofins Cerep (France). CP2 was tested at 10 μM. The compound binding was calculated as a % inhibition of the binding of a radioactively labeled ligand specific for each target. The compound enzyme inhibition effect was calculated as a % inhibition of control enzyme activity. In each experiment and if applicable, the respective reference compound was tested concurrently with the test compound, and the data were compared with historical values determined at Eurofins. The experiment was accepted in accordance with Eurofins validation SOP. Results showing an inhibition (or stimulation for assays run in basal conditions) higher than 50% are considered to represent significant effects of the test compounds. Results showing an inhibition (or stimulation) between 25% and 50% are indicative of weak to moderate effects. Results showing an inhibition (or stimulation) lower than 25% are not considered significant and mostly attributable to the variability of the signal around the control level. Low to moderate negative values have no real meaning and are attributable to the variability of the signal around the control level. High negative values (≥ 50%), which are sometimes obtained with high concentrations of test compounds, are generally attributable to non-specific effects of the test compounds in the assays.

### 250 Kinase Panel

CRO Nanosyn, Inc. (Santa Barbara, CA) was contracted to conduct Kinome Wide Panel (KWP) screening against 250 kinases using the established SOP. CP2 was tested at 1 and 10 μM concentrations.

### *In vivo* CP2 Pharmacokinetic

CP2 bioavailability was determined using C57BL/6J female mice ordered from the Jackson Laboratory. Mice were acclimatized for one week to the new environment prior to initiation of experiments. For evaluating CP2 concentrations in plasma, mice were injected with a single intravenous dose of CP2 (3 mg/kg in DMSO) in a lateral tail vein using a 0.5 cc tuberculin syringe. An independent cohort of mice was administered CP2 by oral gavage (25 mg/kg in 20% PEG 400 and 5% dextrose water in PEG) with a 1 cc tuberculin syringe with a stainless steel 22 gauge straight feeding needle. At 0, 0.08, 0.25, 0.5, 1, 2, 4, 8 and 24 h after treatment, mice were anesthetized, and 200 μl of blood was collected from the retro-orbital sinus through a K2EDTA coated capillary into a K2EDTA coated microtainer tube. Plasma was separated by centrifugation at 4 °C (10,000 rpm x 3 min), transferred to a microcentrifuge tube, immediately frozen on dry ice, and stored at - 80 °C until the analysis. Pharmacokinetic parameters for CP2 were estimated with standard non-compartmental analysis.

### CP2 quantification using LC-MS/MS

Paclitaxel (Sigma, St. Louis, MO) was used as an internal standard (IS). Ultra-pure water was generated using a Barnstead nanopure diamond system (Thermo, Marietta, OH). Optima LC/MS-grade methanol (MeOH, Fisher Scientific, Waltham, MA), analytical grade formic acid (Fisher Scientific, Waltham, MA), 96 well protein crash plates and 1 ml 96 well polypropylene collection plates (Chromtech, Apple Valley, MN), Kunststoff-Kapillaren end-to-end K_2_EDTA coated plastic 30 μl capillary tubes (Fisher Scientific, Waltham, MA) and K_2_EDTA presprayedmicrotainer collection tubes (500 μl, Becton, Dickinson and Company, Franklin Lakes, NJ) were utilized in this study. For controls, drug-free mouse plasma was obtained from healthy CD-1 mice containing 0.1% K_2_EDTA that was purchased from Valley Biomedical and stored at −20 °C for later use. The LC-MS/MS consisted of a Waters Acquity H class ultra-performance liquid chromatography (UPLC) system containing a quaternary solvent manager and sample manager-FTN coupled to a Xevo TQ-S mass spectrometer equipped with an electrospray ionization (ESI) source. Data were acquired and analyzed using Waters MassLynx v4.1 software. Detection of CP2 and paclitaxel (internal standard, IS) was accomplished by multiple reaction monitoring (MRM) using the mass spectrometer in positive ESI mode with capillary voltage, 2.5 kV; source temperature 150 °C; desolvation temperature 400 °C; cone gas flow 150 L/h; desolvation gas flow 800 L/h. The cone voltages and collision energies for CP2 and paclitaxel were determined by MassLynx-Intellistart v4.1 software and were 12 and 18 V (cone) and 30 and 66 eV (collision), respectively. MRM precursor and product ions were monitored at m/z 394.41 > 139.19 for CP2 and 854.29 > 105.08 for paclitaxel. Data were collected from 2 - 5.5 for both CP2 and paclitaxel. The separation of CP2 and paclitaxel was achieved using an Agilent Poroshell 120 EC-C18 (2.7 μ, 2.1 x 100 mm) with an Agilent EC-C18 pre-column (2.7 μ, 2.1 x 5 mm) and a gradient elution program containing ultra-pure water and MeOH, both with 0.1% formic acid. The gradient begins with 70% aqueous, decreases to 10% aqueous over 3 min and holds there for 2 min, then returns to baseline over 0.1 min and holds to equilibrate for 2.9 min. The flow rate was 0.4 ml/min, the total run time was 8 min, injection volume was 5 μl, and the column and autosampler temperatures were 40 °C and 20°C, respectively. Stock solutions of CP2 (5 mg/mL, dissolved in DMSO) and paclitaxel (200 ng/mL, dissolved in acetonitrile, ACN) were prepared in salinized glass vials and stored at −20 °C. 20X working stock solutions were prepared daily and diluted in 1:1 MeOH:H_2_O and stored at −20 °C. Plasma standards containing CP2 (0.2 - 100 ng/mL) were prepared by adding (5 μl) aliquots of 20 X CP2 to plasma (95 μl) in 1.5 mL slick microfuge tubes, 50 μl of this plasma dilution were transferred to a 96 well protein crash plate. Analytes were isolated using protein precipitation with 150 μl ACN containing IS (200 ng/ml). The plate was capped and shaken for 20 min at 1100 rpm. The sample was then vacuum filtered into a Chromtech (Apple Valley, MN) 1 mL 96 well polypropylene collection plate, evaporated to dryness under a gentle stream of nitrogen, and reconstituted with 200 μl H_2_O/ACN (1:1). The collection plate was capped and shaken at 900 rpm for 20 min, and 5 μl aliquots were injected into the LC/MS/MS.

Mass Spectra and ion chromatograms of CP2 and IS were processed using MassLynx v4.1 software with TargetLynx. Standard curves for CP2 and paclitaxel were analyzed using the peak area ratio of CP2 *vs*. IS. CP2 standard curve concentrations were 0.2, 0.5, 1, 5, 10, 20, 50, and 100 ng/ml. The concentration of CP2 in the brain was determined using the same approach.

### Acute CP2 treatment in symptomatic APP/PS1 mice

A cohort of 9-10-month-old female APP/PS1 mice was treated with CP2 (25 mg/kg in 20% PEG 400 and 5% dextrose water in PEG) by oral gavage. Mice were sacrificed at 0, 4, 24, 48 and 72 h (*n* = 1 *per* time point), and Western blot analysis was conducted in the hippocampal tissue to determine the engagement of neuroprotective mechanisms. In an independent cohort of 9 – 10-month-old female and male APP/PS1 mice and their NTG controls (*n* = 3 mice *per* group), baseline glucose uptake was measured at 0 h and after CP2 gavage (25 mg/kg in 20% PEG 400 and 5% dextrose water in PEG) at 24, 48, and 72 h using *in vivo* FDG-PET as discussed below.

### *In vivo* FDG-PET

Mice were fasted one hour prior to i.p. injection of 270 uCi of fludeoxyglucose F18 (18FDG) in 200 uL injection volume prepared the same day at the Mayo Clinic Nuclear Medicine Animal Imaging Resource. Imaging was conducted 30 minutes post injection. Prior to imaging, mice were individually placed in an anesthesia machine (Summit Medical Equipment Company, Bend, Oregon) and anesthetized with 4% isoflurane with 1-2 LPM oxygen. Anesthesia was further maintained with 2% isoflurane delivered by a nose cone. Mice were placed in MicroPET/CT scanner (Scanner Inveon Multiple Modality PET/CT scanner, Siemens Medical Solutions USA, Inc.). Mouse shoulders were positioned in the center of the field of view (FOV), and PET acquisition was performed for 10 min. CT scanning parameters were as following: 360 degree rotation; 180 projections; Medium magnification; Bin 4; Effective pixel size 68.57; Tranaxial FOV 68 mm; Axial FOV 68 mm; 1 bed position; Voltage 80 keV; Current 500 uA; Exposure 210 ms. CT reconstruction parameters: Alogorithm: Feldkamp, Downsample 2, Slight noise reduction with application of Shepp-Logan filter. A final analysis was done using PMOD Biomedical Image Quantification and Kinetic Modeling Software, (PMOD Technologies, Switzerland). The volume of interest (VOI) was created on the CT image of the entire brain. The VOI was applied to the corresponding registered PET scan. The volume statistics were recorded. The percentage of brain glucose uptake was calculated by correcting the measured concentration of uCi recorded after 30 min to the amount of injected dose.

### Chronic CP2 treatment in NTG and symptomatic APP/PS1 mice

NTG and APP/PS1 female mice (*n* = 16 - 21 *per* group) were given either CP2 (25 mg/kg/day in 0.1% PEG dissolved in drinking water *ad lib*) or vehicle-containing water (0.1% PEG) starting at 9 months of age as we described in^13^. Mice were housed 5 *per* cage, water consumption and weight were monitored weekly. CP2 concentration was adjusted based on mouse weight/water consumption weekly. Independent groups of mice were continuously treated for 14 months until the age of 23 months. Seven to eight months after CP2 treatment, mice were subjected to the battery of behavior tests, metabolic cages (CLAMS), FDG-PET, ^31^P NMR spectroscopy, and electrophysiology. After mice were sacrificed, tissue and blood were subjected to Western blot analysis, profiling for cytokines/chemokines, next-generation RNA sequencing, immunohistochemistry, electron microscopy examination, and metabolomics as described below.

### Behavior battery

Behavioral tests were carried out in the light phase of the circadian cycle with at least 24 h between each assessment as we described previously^12^. More than one paradigm ran within 1 week. However, no more than two separate tests were run on the same day. Behavioral and metabolic tests were performed in the order described in the experimental timeline.

#### Open field test

Spontaneous locomotor activity was measured in brightly lit (500 lux) Plexiglas chambers (41 cm × 41 cm) that automatically recorded the activity by photo beam breaks (Med Associates, Lafayette, IN). The chambers were located in sound-attenuating cubicles and were equipped with two sets of 16 pulse-modulated infrared photo beams to automatically record X–Y ambulatory movements at a 100 ms resolution. Data was collected over a 100-minute trial at 30-second intervals.

#### Hanging bar

Balance and general motor function were assessed using the hanging bar. Mice were lowered onto a parallel rod (D < 0.25 cm) placed 30 cm above a padded surface. Mice were allowed to grab the rod with their forelimbs, after which they were released and scored for success (pass or failure) in holding onto the bar for 30 seconds. Mice were allowed three attempts to pass the test. Any one successful attempt was scored as a pass. The final score was presented as an average of three trials *per* animal.

#### Rotarod test

The accelerating rotarod (UgoBasile, Varese, Italy) was used to test balance and coordination. It comprised of a rotating drum that accelerated from 5 to 40 rpm over a 5-minute periods. The latency of each animal to fall was recorded and averaged across three consecutive trials.

#### Novel Object Recognition test (NOR)

was used to estimate memory deficit. All trials were conducted in an isolated room with dim light in Plexiglas boxes (40 cm x 30 cm). A mouse was placed in a box for 5 minutes for acclimatization. Thereafter, a mouse was removed, and two similar objects were placed in the box. Objects with various geometric shape and color were used in the study. Mice were returned to the box, and the number of interrogations of each object was automatically recorded by a camera placed above the box for 10 minutes. Mice were removed from the box for 5 minutes, and one familiar object was replaced with a novel object. Mice were returned to the box, and the number of interrogations of novel and familiar objects was recorded for 10 minutes. Experiments were analyzed using NoldusЕthoVision software. The number of interrogations of the novel object was divided by the number of investigations of the familiar object to generate a discrimination index. Intact recognition memory produces a discrimination index of 1 for the training session and a discrimination index greater than 1 for the test session, consistent with greater interrogation of the novel object.

#### Morris Water Maze

Spatial learning and memory were investigated by measuring the time it took each mouse to locate a platform in opaque water identified with a visual cue above the platform. The path taken to the platform was recorded with a camera attached above the pool. Each mouse was trained to find the platform during four training sessions *per* day for three consecutive days. For each training session, each mouse was placed in the water facing away from the platform and allowed to swim for up to 60 seconds to find the platform. Each training session started with placing a mouse in a different quadrant of the tank. If the mouse found the platform before the 60 seconds have passed, the mouse was left on the platform for 30 seconds before being returned to its cage. If the animal had not found the platform within the 60 seconds, the mouse was manually placed on the platform and left there for 30 seconds before being returned to its cage. A day of rest followed the day of formal testing.

### Aβ ELISA

Levels of Aβ40 and Aβ42 were determined in brain tissue from 23-month-old APP/PS1 mice treated for 14 months with CP2 (*n* = 9) or vehicle (*n* = 9). Differential fractionation was achieved by collecting fractions with most to least soluble Aβ40 and Aβ42 from brain tissue sequentially homogenized in Tris-buffered saline (TBS, most soluble); in TBS containing 1% Triton X-100 (TBS-TX), and 5 M guanidine in 50 mM Tris-HCl (least soluble), pH 8.0 as described in^12^. Levels of human Aβ40 and Aβ42 were determined by ELISA using antibodies produced in-house as previously published^72^.

#### Immunohistochemistry

We followed a protocol described previously^48^. Briefly, for histological analysis, mice were perfused intracardially with 4% paraformaldehyde. Brains were cryoprotected, and serial frozen coronal sections (40 μm) were serially distributed into individual wells of 12-well plates. To facilitate the identification of regions of interest for the quantitative stereological analysis, every 24th section through the entire brain was Nissl stained and compared with the stereotaxic coordinates of the mouse brain^73^. To detect antigens of interest, the sections were incubated in primary antibodies followed by the ABC method (Vector Laboratories) using 3,3’-diaminobenzidine (DAB, Sigma Alrich) as the chromogen for visualization. Antigen retrieval was performed using a Rodent Decloaker for all samples, with an additional 88% formic acid pre-treatment for 4G8-incubated samples. Primary antibodies used are: tyrosine hydroxylase (TH) antibody (rabbit polyclonal, Millipore) for noradrenergic (NAergic) neurites/axons; 4G8 anti-Aβ mouse monoclonal antibody (Biolegend) for amyloid deposits; anti-GFAP rabbit polyclonal antibody (Dako) for astrocytes; and anti-Iba1 rabbit polycloncal antibody (Wako) for microglia. NA and dopaminergic (DA) fibers/neurons were visualized using an anti-tyrosine hydroxylase (TH) antibody (Novus Biologicals). Cresyl violet (CV) was used to stain for nuclei of non-MAergic neurons for neuronal counts.

### Stereological analysis of Aβ deposition and glial reaction

All stereological analysis was performed using the StereoInvestigator software (MicroBrightField, Colchester, VT) as previously described^47^. Tracing of brain regions was based on those defined by *The Mouse Brain in Stereotaxic Coordinates*. Extent of brain area covered by amyloid deposits (4G8 immunostained area), astrocytes (GFAP), or microglia (Iba1) was measured in the S1 barrel cortex (S1BF) and dorsal hippocampal regions. Every 12th section containing the appropriate the cortex/hippocampus was immune-stained and the immunostained areas were determined using the area fractionator protocol, “unstained tissue” was marked with square probes, while the immunostained area was marked with a circular “stained” marker. Because the ratio of untainted and stained markers is proportional to the areas occupied, we calculated the percent of total area that was immunostained.

### Stereological analysis of NAergic afferents and neurons

To determine the length of NAergic axons, we used stereological length estimation with spherical probes (Stereo Investigator; MicroBrightField)^74^. Briefly, virtual spherical probes were placed within a thick section, and the intersection of immunoreactive fibers with the sphere was counted. The lengths were measured at 50 random locations through the reference space. At each focal plane, concentric circles of progressively increasing and decreasing diameters were superimposed, and the intersections with the immunoreactive fibers and circles were counted (*Q*). To minimize surface artifacts, a guard volume of 1 μm was used. This method allows for the simple determination of the total length density (*L*_*V*_) and the total length (*L*). To reduce the effects of variations in the area selection, *L*_*V*_ was routinely used for comparison between groups. Because the densities of NAergic afferents show significant regional variation, we focused our analysis on the selected subregions defined For NAergic afferents. In every eighth section, from the randomly selected starting point within the first eight sections through the entire region of interest, slices were processed for immunocytochemistry. The following regions were examined: barrel field region of primary somatosensory cortex (S1BF; sections between bregma −0.10 to −1.22 mm, posterior to the anterior commissure and anterior to hippocampus) and dorsal hippocampus (dentate, CA1, CA2/3; sections between bregma −1.46 to −2.18 mm). To determine the total number of NAergic neurons in *Locus Coeruleus* (LC), every 4th section of the entire region containing the LC that was immunostained for TH and counterstained with CV. Total TH+ and TH-neuron numbers were estimated using the optical fractionator probe; the LC region of interest was traced and magnified at 100x, and TH+ neurons and TH-nuclei were counted using the counting frame of 40 x 30 μm. For volume measurements, neurons were randomly sampled within the sections used for neuron counts. Using the nucleator feature of the software, we determined neuron volumes by placing four rays through the cell, with the nucleolus serving as the midpoint.

### Inflammatory markers

After 16 h of fasting, blood from 23-month-old APP/PS1 and NTG mice that had been treated for 14 months with CP2 (*n* = 15 - 20 *per* group) or vehicle (n = 15 - 20 *per* group) was collected by orbital bleed and centrifuged for 10 minutes at 2,500 rpm. Collected plasma was sent for 32-plex cytokine array analysis (Discovery Assay, Eve Technologies Corp. https://www.evetechnologies.com/discovery-assay/). The multiplexing analysis was performed using the Luminex 100 system (Luminex). More than 80% of the targets were within the detectable range (signals from all samples were higher than the lowest standard). For the data that were out of range (OOR <), their values were designated as 0 pg/mL. Blood samples were run in duplicates.

### Lipid peroxidation assay

Levels of malondialdehyde (MDA), a product of lipid degradation that occurs as a result of oxidative stress, were measured using an MDA assay kit (#MAK085, Sigma Aldrich) in hippocampal brain tissue isolated from 23 month old APP/PS1 and NTG mice treated for 14 months with CP2 or vehicle (*n* = 5 *per* group), according to the manufacturer’s instructions.

### Levels of senescent cells in adipose tissue

The senescent cell burden was assayed by senescent associated β-galactosidase (SA-β-Gal) staining of freshly isolated fat biopsies from 23-month-old APP/PS1 and NTG mice, treated for 14 months with CP2, or vehicle (n = 5 - 6 *per* group) as previously described^75^. In brief, about 100 mg of periovarian and inguinal fat pads were fixed in PBS containing 2.0% formaldehyde and 0.2% glutaraldehyde for 10 minutes. After fixation, tissue was washed and incubated with SA-β-gal staining solution for 16 −18 h at 37°C. The enzymatic reaction was stopped by washing tissue with ice cold PBS. Tissues were counterstained with DAPI, and ten to twelve random images were taken *per* sample with an EVOS microscope under 20X magnification in bright and fluorescent fields. SA-β-gal positive senescence cells and total cell number (DAPI^+^ nuclei) were quantified *per* field. SA-β-gal positive cell numbers were expressed as a percent of the total cell number *per* image.

### Metabolomics

#### Ceramide panel

Blood was collected from fasting (16 h) 23-month-old APP/PS1 and NTG mice treated for 14 months with CP2 or vehicle (*n* = 5 *per* group) by orbital bleeding. Concentrations of ceramides were established using targeted metabolomics at the Mayo Clinic Metabolomics Core using their established SOP as described previously^76^.

### Metabolomics profiling in brain

Fasting (12 h) APP/PS1 and NTG mice treated for 6 months with CP2 or vehicle (*n* = 5 *per* group) were sacrificed by cervical dislocation; brains were rapidly removed; hippocampal tissue was dissected and immediately flash-frozen in liquid N_2_. Tissue was pulverized under liquid N_2_ and extracted in a solution containing 0.6 M HClO_4_ and 1mM EDTA. Extracts were neutralized with 2M KHCO_3_ and used for metabolomic analysis as described in^77^. An aliquot of 100 μL of extract was transferred into Eppendorf tube and spiked with 5 μL IS, myristic-d27 acid (1 mg/mL) at ambient temperature. Samples were gently vortexed and completely dried in a SpeedVac concentrator. The lyophilized brain samples were methoxiaminated and derivatized same way as for plasma samples. For GC-MS analysis, we used conditions previously optimized for an Agilent 6890 GC oven with Agilent 5973 MS^78^. Nucleotides were separated on a reversed-phase Discovery C18 columns (SIGMA, St. Louis, MO) with Hewlett-Packard series 1100 HPLC system (Agilent, Santa Clara, CA). The Agilent Fiehn GC/MS Metabolomics RTL Library was employed for metabolite identification. GC-MS spectra were deconvoluted using AMDIS software. After that, SpectConnect software was used to create the metabolite peaks matrices.

### Electrophysiology

APP/PS1 and NTG mice 15-18 months of age treated for ~10 months with CP2 or vehicle (*n* = 5 *per* group) were used for electrophysiology analysis. Mice were deeply anesthetized with isoflurane and decapitated. Brains were quickly removed and transferred into a cold slicing solution containing an artificial cerebrospinal fluid (ACSF), where NaCl was substituted with sucrose to avoid excitotoxicity. Transverse slices were made at 300 - 350 μm thickness using a vibratome (VT-100S, Leica). Slices were incubated in ACSF containing 128 mM NaCl, 2.5 mM KCl, 1.25 mM NaH2PO_4_, 26 mM NaHCO_3_, 10 mM glucose, 2 mM CaCl_2_, and 1 mM MgSO_4_, aerated with 95% O_2_/5% CO_2_. Slices were maintained at 32 °C for 13 min, and then maintained at room temperature throughout the entire experiment. For electrophysiological recording, each slice (2 - 3 slices *per* mouse) was transferred to a recording chamber, and ACSF was continuously perfused at the flow rate of 2 - 3 ml/min. A single recording electrode and a single bipolar stimulation electrode were placed on top of the slice. A boron-doped glass capillary (PG10150, World Precision Instruments) was pulled with a horizontal puller (P-1000, Sutter Instrument) and filled with ACSF for extracellular recording. Under the microscope (FN-1, Nikon), the recording electrode was placed in the CA1 area of the hippocampus. The bipolar stimulation electrode (FHC) was placed at the Schaffer collaterals. The distance between two electrodes was over ~200 μm. To define a half response of stimulation, various intensities of electrical stimulation were applied (10 - 500 μA). However, the pulse width was fixed at 60 μsec. Once the stimulation parameter was determined to generate a half maximum of evoked fEPSP, this stimulation intensity was used for the paired pulse and LTP experiments. The paired pulse stimulation protocol was applied with 50, 100 and 200 mseconds intervals. For the LTP experiment, test stimulation was applied every 30 seconds for 30 minutes to achieve a stable baseline. Once the stable baseline was achieved, a tetanic stimulation (100 Hz for 1 seconds) was applied three times at a 30-second intervals. Initial slopes of the fEPSP were used to compare synaptic strength. fEPSP slops were analyzed using pCLAMP v. 10.5 software.

### Mitochondrial DNA (mtDNA) copy number

Genomic DNA was isolated from snap-frozen cortico-hippocampal brain sections from NTG and APP/PS1 mice treated with vehicle or CP2 (*n* = 5 *per* group) for 14 months using a DNeasy Blood & Tissue Kit (QIAGEN, cat. # 69504) according to the manufacturer’s instructions. Quantification of mtDNA copy number was performed in triplicates using 100 ng of isolated DNA. qRT-PCR was performed using a TaqMan One-Step RT-PCR Master Mix Reagents Kit (ThermoFisher, cat. # 4309169) and primers for a genomic locus (*b-actin*) and a mitochondrial gene (*mND5*). Primers for *b-actin* were the following: forward sequence (5`-GAT CGA TGC CGG TGC TAA GA-3`); reverse sequence (5`-GGA AAA GAG CCT CAG GGC AT-3`). Primers for the amplification of *mND5*: forward sequence (5`-TGT AAA ACG ACG GCC AGT AGC CCT TTT TGT CAC ATG AT-3`); reverse sequence (5`-CAG GAA ACA GCT ATG ACC GGC TCC GAG GCA AAG TAT AG-3`). The relative mitochondrial DNA copy number was calculated as a ratio of genomic versus mitochondrial DNA. Delta Ct (ΔCt) equals the sample Ct of the mitochondrial gene (*mND5*) subtracted from the sample Ct of the nuclear reference gene (*B-Actin*).

### Electron Microscopy (EM)

#### Two-dimensional transmission EM (2D TEM)

Hippocampal tissue from APP/PS1 and NTG mice treated with CP2 or vehicle for 12 - 14 months (*n* = 5 *per* group) was dissected, cut into 1 mm thick sections, and fixed in 4% paraformaldehyde + 1% glutaraldehyde in 0.1 M phosphate buffer for 24 h. Fixed tissue was further cut into smaller (2 mm^3^) pieces and placed into 8 ml glass sample vials (Wheaton, cat# 225534). Processing was facilitated by the use of a Biowave^®^ laboratory microwave oven set to 150 W (Ted Pella Inc., Redding, CA) and included the following steps: (1) 0.1 M phosphate buffer (PB), pH 7.0, 40 seconds on, 2 min rest at RT, repeat three times; (2) 2% osmium tetroxide in H_2_O, 40 seconds on, 40 seconds off, 40 sec on, 15 minutes rest at RT; (3) H_2_O rinse, 40 seconds on, 2 minutes rest at RT, repeat three times; (4) 2% aqueous uranyl acetate, 40 seconds on, 40 seconds off, 40 seconds on, 15 minutes rest at RT; (5) H_2_O rinse, 40 seconds on, 2 minutes rest at RT, repeat three times; (6) sequential dehydration in ethanol series 60%, 70%, 80%, 95%, 100%, 100% acetone, 100% acetone, 40 seconds on, 2 minutes rest at RT each. Resin infiltration steps were performed as follows: (1) 1:2 resin:acetone, 40 seconds on, 40 seconds off, 40 seconds on, 15 minutes rest with the vacuum; (2) 1:1 resin:acetone, 40 seconds on, 40 seconds off, 40 seconds on, 15 minutes rest with the vacuum; (3) 3:1 resin:acetone, 40 seconds on, 40 seconds off, 40 seconds on, 30 min rest with the vacuum; (4) 100% resin 40 seconds on, 40 seconds off, 40 seconds on, overnight rest with the vacuum. The next day samples were moved to fresh resin in embedding molds and incubated at 60 °C for 24 h to polymerize. Ultrathin sections (silver interference color, ~0.1 μm) were place on 150 mesh copper grids and stained with lead citrate. Images were acquired using a JEOL 1400+ TEM operating at 80 kV equipped with a Gatan Orius camera. For conventional 2D TEM, approximately 50 images were taken randomly within each section for the analysis. All images used for the analysis were taken at 5000 x magnification. For 2D TEM analysis of mitochondrial morphology, organelles were scored according to their appearance as elongated (> 2 μm long) or small (circular 0.5 μm in diameter). Twenty random areas from each CA1 region were imaged, and only neuropils longer than 3 μm were selected for the analyses.

### Three-dimensional (3D) EM using Serial Block Face Scanning Electron Microscopy (SBFSEM)

Images for 3D EM reconstructions were obtained using an ApreoVolumeScope (Thermo Fisher Scientific) electron microscope that combines an integrated serial block face microtome (SBF) and high-resolution field emission scanning (SEM) imaging. Hippocampal CA1 region was dissected from vehicle and CP2-treated NTG and APP/PS1 mice, cut into 2 mm^3^ pieces, and immersion-fixed in neutral 2.5% glutaraldehyde + 2.5% paraformaldehyde in 0.1 M cacodylate buffer + 2 mM CaCl_2_. Following 24 h fixation, samples were processed using the following protocol based on the serial block-face method developed by the National Center for Microscopy and Imaging Research (La Jolla, CA; https://ncmir.ucsd.edu/sbem-protocol): (1) samples were rinsed 4 x 3 minutes in 0.1 M cacodylate buffer + 2 mM CaCl_2_, (2) incubated in 2% osmium tetroxide in 0.15M cacodylate buffer for 1.5 h rotating at RT, (3) incubated in 2% osmium tetroxide + 2% potassium ferrocyanide in 0.1 M cacodylate for 1.5 h rotating at RT, (4) rinsed in H_2_O 4 x minutes, (5) incubated in 1% thiocarbohydrazide (TCH) 45 minutes at 50 °C, (6) rinsed in H_2_O 4 x 3 minutes, (7) incubated in fresh 2% osmium tetroxide in H_2_O 1.5 h rotating at RT, (8) rinsed in H_2_O 4 x 3 minutes, (9) incubated in 1% aqueous uranyl acetate overnight at 4 °C, (10) further incubation in uranyl acetate 1 h at 50 °C, (11) rinsed in H_2_O 4 x 3 minutes, (12) incubated in lead aspartate 1 h at 50 °C, (13) rinsed in H_2_O 4 x 3 minutes, (14) dehydrated through ethanol series (60, 70, 80, 95, 100, 100%) 10 minutes each, (15) two rinses in 100% acetone 10 minutes each, (16) resin 1:2, 1:1, 3:1 in acetone 0.5 h, 1 h, 2 h, overnight in 100% resin. Samples were embedded into the Durcapan hard resin (EMS, Hatfield, PA), and allowed to polymerize at a minimum of 24 h prior to trimming and mounting. Tissue was trimmed of all surrounding resin and adhered to 8 mm aluminum pins (Ted Pella Inc., Redding, CA) using EpoTek silver epoxy (EMS, Hatfield, PA). A square tower (0.5 mm) was trimmed from the tissue using a Diatome ultratrim knife (EMS, Hatfield, PA) and the entire pin was coated with gold palladium. Following coating, the block was trimmed to a planer surface using a diamond knife, and the pin was mounted in the SEM microtome. Serial block-face images were acquired using a Thermo Fisher Volumescope (Thermo Fisher, Inc., Waltham, MA) at 50 nm section depths with detector acceleration voltage of 1.5 kV. Voxel size ranged between 8 and 14 nm according to magnification. For each region of interest (ROI), 400 sections were obtained, registered, and filtered using non-local means. All images used for the analysis were adjusted to 5,000 x magnification. Segmentation and three-dimensional analysis was performed using *Reconstruct* (SynapseWeb)^79^ and *Amira* 6.4 software (Thermo Fisher, Inc., Waltham, MA).

### Image segmentation and quantitative morphometric analysis of dendritic spines using 3D EM

Aligned and normalized stacks of serial sections were further processed with unsharp masking, Gaussian blur and non-local means filters were applied in order to clearly distinguish cellular membranes. Dendrites, dendritic spines and synapses (the PSD and the opposed presynaptic membranes) were segmented by manually tracing contours in consecutive micrographs. Each segmented dendritic spine or synaptic junction was identified independently. Once the dendrite reconstruction was completed in *Reconstruct*, traces were modified by an absolute intensity and maximal thresholding approach and exported in JPEG format into *Amira* software. The magic wand tool in *Amira’s* segmentation mode was used to pull out the reconstructions from the *Reconstruct* exports. Dendritic spines were cut from their parent dendritic shaft through the base of their neck in a 3D optimal orientation and analyzed using the *label analysis function in Amira’s project view*. The label analysis was customized to measure 3D length, surface area, and volume of the spines. By using the *measure tool* in *Amira’s* project view, length, width of the head and length of the neck of each dendritic spine were established. The following classifications for dendritic spine type were adapted from^38^: branched (2 or more heads); filopodium (length > 2 μm, no bulbous head); mushroom (length 1 < x > 2 μm; head > 0.6 μm); long thin (length 1 < x < 2 μm; head < 0.6 μm); thin (length <1 μm; head < 0.6 μm); and stubby (length < 1 μm; length: width ratio < 1). In addition to the frequency of individual spine types, their activity (based on the presence of synaptic vesicles), volume and size (length and width) of spines were also estimated. To evaluate the impact of nanoscale alterations in spine morphology on the diffusional coupling between spines and dendrites that promote long-term potentiation, we calculated the compartmentalization factor, which is defined as CF=V*L/A where ***V*** is the spine head volume, ***L*** is the spine neck length, and ***A*** is the cross-sectional area of the spine neck^40^.

### Next-generation RNA sequencing

Brain tissue, encompassing the hippocampal and cortical regions, from APP/PS1 and NTG mice treated with vehicle of CP2 for 14 months (*n* = 5 *per* group) were lysed in QIAzol (Qiagen cat. # 79306) followed by RNA isolation using miRNeasy (Qiagen cat. # 217004) according to the manufacturer’s instructions. The quantity and quality of RNA were measured using a NanoDrop spectrophotometer and Agilent 2100 Bioanlyzer, respectively. All RINs (RNA integrity numbers) had a value greater than eight.

#### Library preparation and sequencing

Total RNA (200 ng) was used to generate libraries using TruSeq RNA Library Prep Kit v2 (Illumina). All samples were sequenced at the Mayo Clinic Medical Genome Facility (MGF) Sequencing Core by Illumina HiSeq 4000 with paired end 101-bp read length. Approximately 50 million single fragment reads were acquired *per* sample.

#### Bioinformatics methods

MAP-RSeq v2.1.1, a comprehensive computational pipeline developed by the Mayo Clinic’s Division of Biomedical Statistics and Informatics, was used to analyze RNA-Sequencing data. MAP-RSeq uses a variety of publicly available bioinformatics tools tailored by methods developed in-house. The main outputs of the MAP-RSeq workflow are the gene counts, expressed single nucleotide variants (eSNVs), gene fusion candidates, and quality control plots. The aligning and mapping of reads was performed using TopHat2 against the mm10 reference genome. The gene and exon counts were generated by FeatureCounts using the gene definitions files from Ensembl. ‘-O’ option within FeatureCounts was used to account for expression derived from regions shared by multiple genomic features. FeatureCounts was also executed for quantifying expression on a *per* exon basis by utilizing the ‘-f’ option. RSeqQC^80^ was used to create a variety of quality control plots to ensure the results from each sample were reliable and could be collectively used for differential expression analysis. Upon the completion of all the deliverables from MAP-RSeq, a html document was created tying everything together in one interactive document. The R bioinformatics package DeSeq2 was used for differential gene expression analysis. The criteria for the selection of significant differentially expressed genes was P value < 0.05. Gene function enrichment was determined using the Database for Annotation, Visualization and Integrated Discovery (DAVID v. 6.8)^81^. Mouse transcriptional factors were downloaded from the Riken genome database and mapped to our RNA-seq differential gene list. The clustering heatmaps were generated based on unsupervised hierarchical clustering using Pearson correlation distance. Most graphs were generated using customized R programs. Data and materials availability: RNA seq data from NTG and APP/PS1 mice with and without CP2 treatment generated in this study are available at https://www.ncbi.nlm.nih.gov/geo/query/acc.cgi?acc=GSE149248 (GEO accession ID GSE149248).

### Mapping mouse genes to human AD genes from the AMP-AD study

The differentially expressed genes from our RNA-seq data using mouse model were identified as described in the differential analysis section. The up- and down-regulated genes in comparison of control and AD females were from the AMP-AD data set. The human data and analysis process were described in https://www.biorxiv.org/content/10.1101/510420v1. The up-regulated genes in the RNA-seq data of the mouse NTG *vs.* AD comparison were overlapped with the human up-regulated gene list. The down-regulated genes in mouse data were overlapped with the human down-regulated list. Gene function enrichment analysis was performed on the overlapped list of DEGs using NCBI DAVID function analysis tools with a P value cutoff of less than 0.05. Similar analyses were performed on the 567 genes overlapped in the two differential gene lists of NTG *vs.* AD and AD *vs.* AD+CP2 comparisons. All genes are presented in the Supplemental Tables. 6-23.

### Comprehensive Laboratory Animal Monitoring System (CLAMS)

The CLAMS (Columbus Instruments, Columbus, OH) allows automated, non-invasive, and simultaneous monitoring of horizontal and vertical activity, feeding and drinking, oxygen consumption, and CO_2_ production of an individual mouse. APP/PS1 and NTG mice treated with vehicle or CP2 (16 – 18-month-old, *n* =15 - 20 *per* group) were individually placed in CLAMS cages. Indirect calorimetry was monitored over 2 days, when mice were allowed food for 24 h *ad lib* (Fed state), and for the following 24 h food was removed (Fasting state). Mice were maintained at 20 – 22°C under a 12:12 h light–dark cycle. All mice were acclimatized to CLAMS cages for 3 – 6 h before recording. Sample air was passed through an oxygen sensor for the determination of oxygen content. Oxygen consumption was determined by measuring oxygen concentration in the air entering the chamber compared with air leaving the chamber. The sensor was calibrated against a standard gas mix containing defined quantities of oxygen, carbon dioxide, and nitrogen. Food and water consumption were measured directly. The hourly file displayed measurements for the following parameters: VO_2_ (volume of oxygen consumed, ml/lg/h), VCO_2_ (volume of carbon dioxide produced, ml/kg/h), RER (respiratory exchange ratio), heat (kcal/h), total energy expenditure (TEE, kcal/h/kg of lean mass), activity energy expenditure (AEE, kcal/h/kg of lean mass), resting total consumed food (REE kcal/h/kg of lean mass), food intake (g/kg of body weight/12 h), metabolic rate (kcal/h/kg), total activity (all horizontal beam breaks in counts), ambulatory activity (minimum 3 different, consecutive horizontal beam breaks in counts), and rearing activity (all vertical beam breaks in counts). The RER and atty acid (FA) oxidation were calculated using the following equations: RER = VCO_2_/VO_2_; FA oxidation (kcal/h)= TEE X (1-RER/0.3). Daily FA oxidation was calculated from the average of 12 h of hourly FA oxidation. Daily carbohydrate plus protein oxidation was calculated from average of 12 h of hourly TEE minus daily FA oxidation. Metabolic flexibility was evaluated from the difference in RER between daily fed state and fasted state recorded at night phase, according to the following equations: Δ = 100% * (RER fed-RER fasted)/RER-fed.

### Dual energy X-ray absorptiometry (DEXA)

A LUNAR PIXImus mouse densitometer (GE Lunar, Madison, WI), a dual-energy supply X-ray machine, was used for measuring skeletal and soft tissue mass for the assessment of skeletal and body composition in CP2 or vehicle-treated mice. Live mice were scanned under 1.5 - 2% isoflurane anesthesia. Mice were individually placed on plastic trays, which were then placed onto the exposure platform of the PIXImus machine to measure body composition. The following parameters were generated: lean mass (in grams), fat mass (in grams), and the percentage of fat mass. These parameters were used to normalize data generated in CLAMS, including O_2_, VCO_2_, metabolic rate and energy expenditure.

### In vivo 31P NMR spectroscopy

NMR spectra were obtained in 16 – 18-month-old APP/PS1 and NTG mice treated with CP2 or vehicle for 8 months (*n* = 5 *per* group) using an AVANCE III 300 MHz (7 T) wide bore NMR spectrometer equipped with micro-imaging accessories (Bruker BioSpin, Billerica, MA) with a 25-mm inner diameter dual nucleus (^31^P/^1^H) birdcage coil. For anatomical positioning, a pilot image set of coronal, sagittal, and axial imaging planes were used. For ^31^P NMR spectroscopy studies, a single pulse acquisition with pulse width: P1 200 us (~30 degrees), spectral width: SW 160 ppm, FID size: TD 16 k, FID duration: AQT 0.41 s, waiting time: D1 1second and number of scans: NS 512 was used. The acquisition time was 12 minutes. Spectra were processed using TopSpin v1.5 software (BrukerBiospin MRI, Billerica, MA). Integral areas of spectral peaks, corresponding to inorganic phosphate (Pi), phosphocreatine (PCr), and γ, α, and β phosphates of adenosine triphosphate (αATP, βATP, and γATP), were analyzed using jMRUI software. Since in some mice, the Pi peaks were not detectable, Pi values were uniformly omitted from the analyses. The presence of phosphomonoester (PME) or phosphodiester (PDE) peaks was also recorded. However, the signal-to-noise ratio of these peaks was not always adequate for accurate quantification. Levels of PCr and Pi were normalized to the total ATP levels present in that spectrum or to the amount of βATP. Results were consistent between both of these normalization methods; data are presented as the ratio of each parameter to total ATP.

### Western blot analysis

Levels of proteins were determined from the cortico-hippocampal region of the brain of vehicle and CP2-treated NTG and APP/PS1 mice (*n* = 6 - 8 mice *per* group) by Western blot analysis. Tissue was homogenized and lysed using 1× RIPA Buffer plus inhibitors. Total protein lysates (25 μg) were separated in equal volume on 4 – 20% Mini-PROTEAN TGX™ Precast Protein Gels (Bio-Rad, cat. # 4561096) and transferred to an Immun-Blot polyvinylidene difluoride membrane (PVDF cat. # 1620177). The following primary antibodies were used: phospho-AMPK (Thr 172) (1:1000, Cell Signaling Technology, cat. # 2535), AMPK (1:1000, Cell Signaling Technology, cat. #2532), phospho-Acetyl-CoA Carboxylase (Ser79) (1:1000, Cell Signaling Technology, cat. #11818), Synaptophysin (1:200, Santa Cruz Biotechnology, Santa Cruz, CA, cat. # 17750), BDNF (1:200, Santa Cruz Biotechnology, Santa Cruz, CA, cat. # 546), PSD95 (1:1000, Cell Signaling Technology, cat. # 2507), Sirt3 (1:1000, Cell Signaling Technology, cat. #5490), IGF1 (1:2000, Abcam, cat. # ab223567), Superoxide Dismutase 1 (1:1000, Abcam, cat. # ab16831), phospho-IGF-I Receptor β (Tyr1131)/Insulin Receptor β (Tyr1146) (1:1000, Cell Signaling Technology, cat. # 3021), IGF-I Receptor β (1:1000, Cell Signaling Technology, cat. #3027), phospho-Pyruvate Dehydrogenase α1 (Ser293) (1:1000, Cell Signaling Technology, cat. #31866), Pyruvate Dehydrogenase (1:1000, Cell Signaling Technology, cat. #3205), Glut4 (1:1000, Cell Signaling Technology, cat. #2213), Glut3 (1:1000, Santa Cruz Biotechnology, cat. # sc-74399), phospho-ULK1 (Ser555) (1:1000, Cell Signaling Technology, cat. #5869), phospho-ULK1 (Ser317) (1:1000, Cell Signaling Technology, cat. #12753), phospho-Beclin-1 (Ser15) (1:1000, Cell Signaling Technology, cat. # 84966), PGC1α (1∶1000, Calbiochem, cat. # KP9803);, TFAM (1:1000, Sigma-Aldrich, cat. #AV36993), phospho-NF-κB p65 (Ser536) (1:1000, Cell Signaling Technology, cat. #3033), HO-1 (1:1000, Cell Signaling Technology, cat. #70081), IκBα (1:1000, Cell Signaling Technology, cat. #4812), Nrf2 (1:1000, Abcam, cat. # ab62352), LC3B (1:1000, Novus Biologicals, cat. # NB100-2220), TFEB (1:500, Thermo Fisher, cat. #PA5-75572), LAMP1 (1:1000, Cell Signaling Technology cat. #9091), Catalase (1:1000, Cell Signaling Technology cat. # 14097), Tubulin (1:5000, Biovision, cat. #3708), β-Actin (1:5000, Sigma-Aldrich, cat. # A5316). The following secondary antibodies were used: donkey anti-rabbit IgG conjugated with Horseradish Peroxidase (1:10000 dilution, GE Healthcare UK Limited, UK) and sheep anti-mouse IgG conjugated with Horseradish Peroxidase (1:10000 dilution, GE Healthcare UK Limited, UK). Ban quantification was done using Image Lab^TM^ v. 6.0.

### Statistics

The statistical analyses were performed using the GraphPad Prism (Version 8, GraphPad Software, Inc., La Jolla, Ca). Statistical comparisons among four groups concerning behavioral and metabolic tests, immunoreactivity, metabolomics, plasma cytokine panel, body composition, electron microscopy imaging, FDG-PET, ^31^P NMR, electrophysiology, were analyzed by two-way ANOVA, the two-tailed unpaired and paired Student *t* test, where appropriate. The Fisher’s LSD *post hoc* analysis was used if significant interaction among groups was found. A linear regression analysis was applied to determine differences among the groups in body weights and age-related loss of TH+ axons and neurons. Significant differences between vehicle and CP2-treated groups within the same genotype and differences among NTG, APP/PS1 and APP/PS1+CP2 mice were considered in the final analysis. Data are presented as mean ± S.E.M. for each group of mice.

## Acknowledgements

We thank Mayo Clinic Cores for help with FDG-PET; RNA-seq, metabolomics, EM and CLAMS data acquisition; Dr. A. Leontovich, Ms. R. Schlichte, E. Murray, L. G. Andres-Beck and Mr. I. Trushin for help with illustrations. This research was supported by grants from the National Institutes of Health NIA RF1AG55549 (to ET), NINDS R01NS107265 (to ET), RO1AG062135 (to ET and MKL), ADDF 291204 (to ET), MN Partnership for Biotechnology and Medical Genomics #15.08 (to ET and MKL), NIH RO1NS88260 (to SYC), National Cancer Institute Grant P30 CA015083 (to JMR), NIH grants R37AG013925 and P01AG062413 (to JLK and TT), the Alzheimer’s Association Part the Cloud Program, Robert and Arlene Kogod, the Connor Group, Robert J. and Theresa W. Ryan, and the Noaber Foundation. Its contents are solely the responsibility of the authors and do not necessarily represent the official view of the NIH. The funders had no role in study design, data collection and analysis, decision to publish, or preparation of the manuscript.

## Author Contributions

All authors listed have made a substantial, direct, and intellectual contribution to the work, and approved its publication.

## Data availability statement

RNA-seq data generated in this study using mice that support the findings are under the deposition to the public repository and will be available before publication.

---

**Extended Data Fig. 1.**
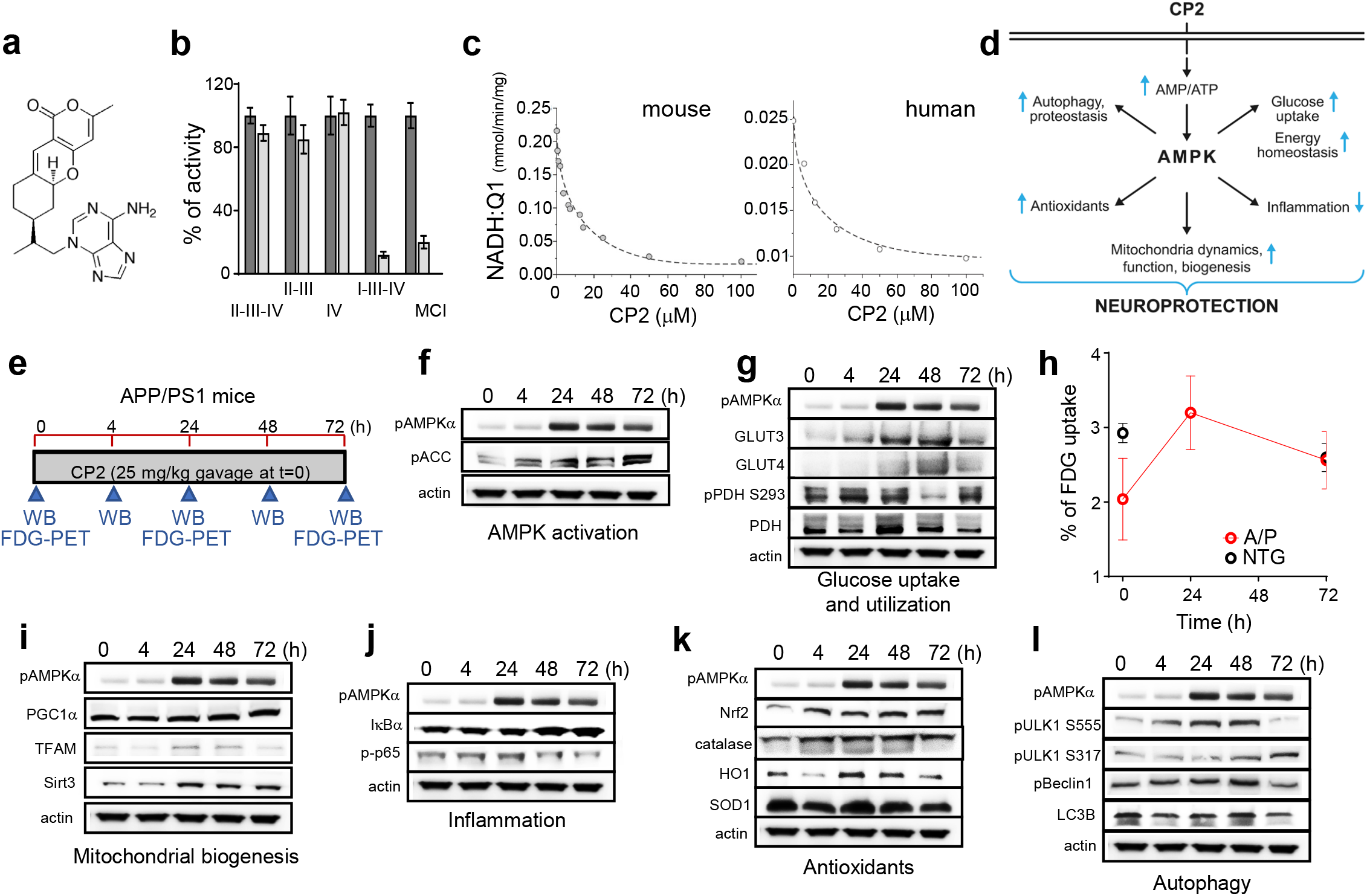
Acute CP2 treatment activates multiple AMPK-dependent neuroprotective mechanisms in symptomatic APP/PS1 mice improving energy homeostasis in the brain. **a,** CP2 structure. **b**, CP2 (light grey bars, 50 mM) does not affect the activity of succinate oxidase (complexes II-III-IV), succinate:cytochrome *c* reductase (complexes II-III), and ferrocytochrome *c* oxidase (complex IV only), but significantly inhibits MCI affecting NADH oxidase (complexes I-III-IV) and NADH:ubiquinone (complex I only) in mouse brain mitochondria. Vehicle, dark grey bars. **c**, CP2 inhibits MCI in mitochondria isolated from the mouse and postmortem human cortical tissue. **d**, Neuroprotective pathways activated by CP2 in the brain converge on AMPK activation. **e**, Timeline of acute CP2 administration via gavage (25 mg/kg) to s APP/PS1 mice 9 −10-month-old. Brain tissues from 1 mouse *per* each time point were examined using western blot analysis. Independent cohort of mice was subjected to the *in vivo* FDG-PET. **f-l**, Western blot analysis of brain tissue from acutely gavaged APP/PS1 mice from (**e**) confirms that CP2 activates multiple AMPK-dependent neuroprotective mechanisms in symptomatic APP/PS1 mice. **h**, Untreated symptomatic APP/PS1 mice 9 – 10-month-old have reduced glucose uptake in the brain determined using *in vivo* FDG-PET compared to NTG littermates (time 0 h). Acute CP2 oral administration improves glucose uptake brining glucose levels in APP/PS1 mice to the same level as in untreated age- and sex-matched NTG mice (time 24 and 72 h); *n* = 3 mice *per* group.

**Extended Data Fig. 2.**
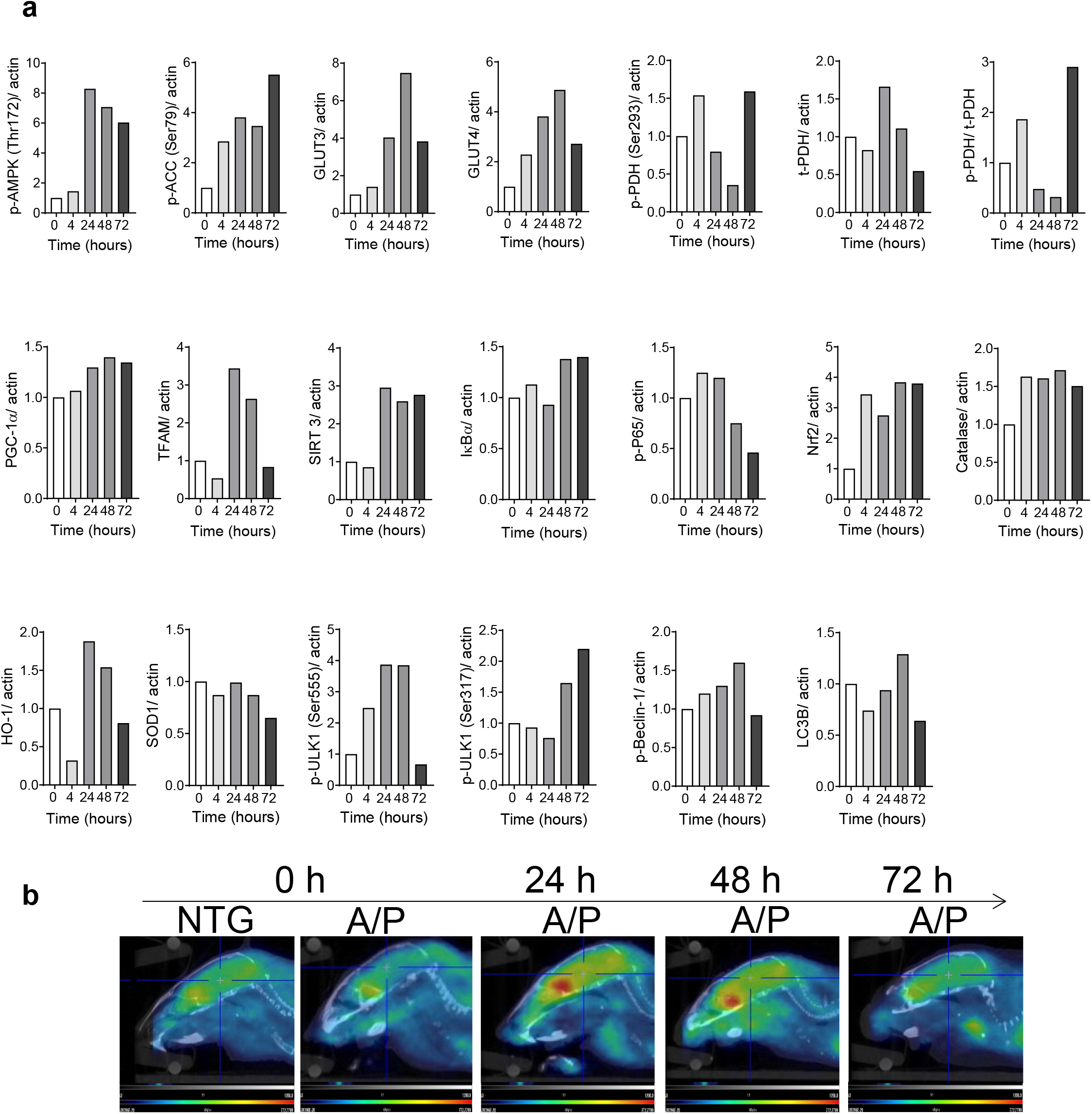
CP2 activates AMPK-dependent neuroprotective pathways in symptomatic APP/PS1 mice and improves glucose uptake in the brain. **a,** Quantification of time-dependent changes in levels of key proteins involved in multiple neuroprotective pathways in the brain of symptomatic APP/PS1 mice acutely gavaged with CP2 from experiments described in Extended Data Fig. 1e (25 mg/kg, *n* = 1 *per* time point) using Western blot analysis. **b**, Representative FDG-PET images collected from the independent cohort of APP/PS1 mice (9 – 10-month-old) before CP2 administration and 24, 48 and 72 h after a single dose via gavage (25 mg/kg, Extended Data Fig. 1e). NTG, untreated non-transgenic age- and sex-matched littermates. *n* = 3 female mice *per* group.

**Extended Data Fig. 3.**
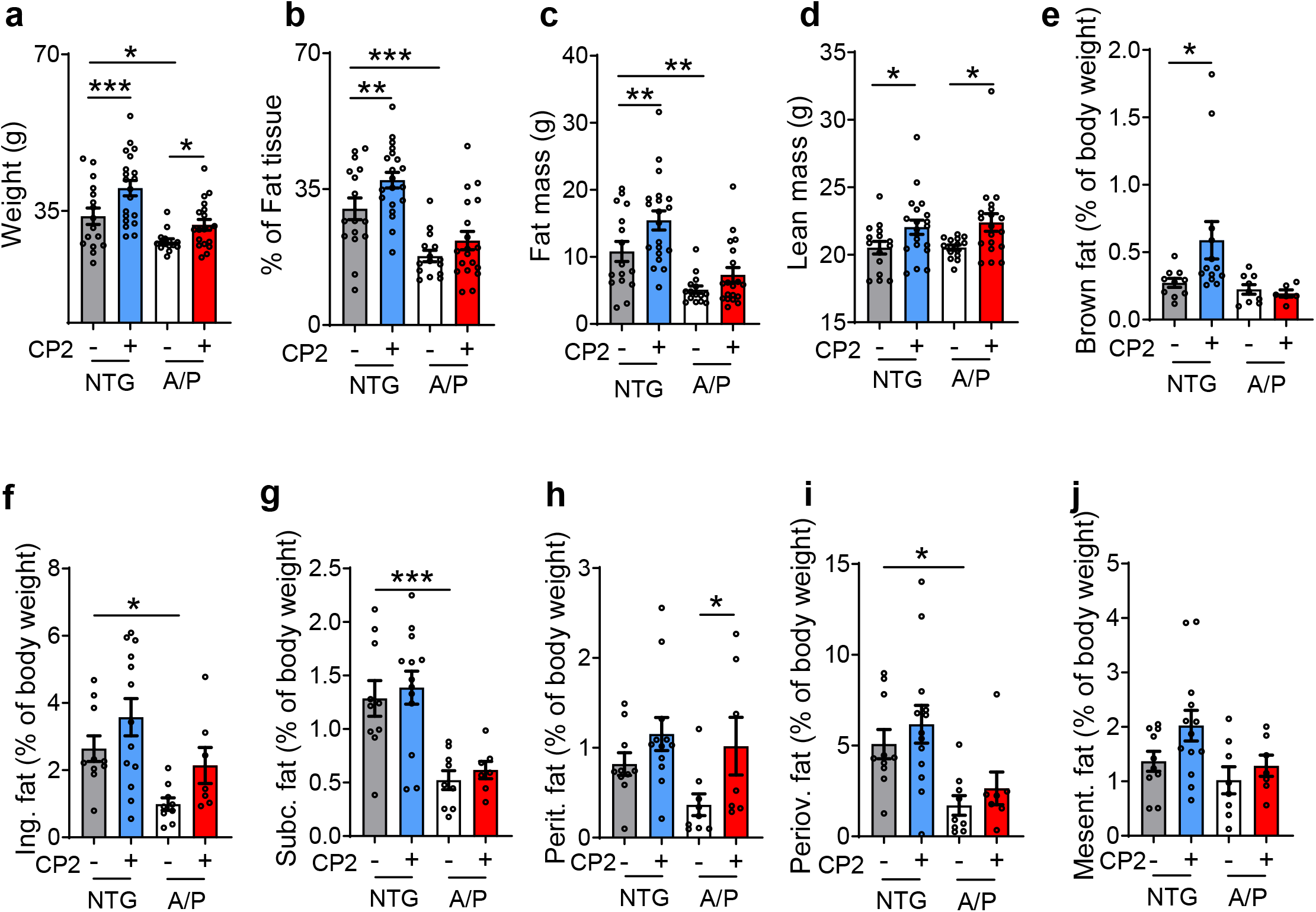
CP2 treatment ameliorates body weight loss in symptomatic APP/PS1 mice. **a,** Body weight was significantly increased in CP2-treated NTG and APP/PS1 mice relative to untreated counterparts. Vehicle-treated APP/PS1 mice were significantly lighter compared to untreated NTG mice. CP2-treated APP/PS1 mice did not differ significantly from vehicle-treated NTG mice. **b,c,** CP2 treatment significantly increased fat mass in NTG mice with a similar trend in APP/PS1 mice compared to vehicle-treated counterparts. **d,** CP2 treatment significantly increased lean mass in NTG and APP/PS1 mice. **e,f,** CP2 treatment increased brown fat in NTG mice (**e**), with a trend towards increased inguinal fat in both NTG and APP/PS1 mice (**f**). **g,** CP2 treatment did not affect subcutaneous fat mass in APP/PS1 or NTG mice. **h,i,** CP2 treatment significantly increased a percentage of peritoneal fat in APP/PS1 mice (**h**) with a similar trend for periovarian fat in APP/PS1 and NTG mice (**h,i**). **j,** CP2 did not affect the percentage of mesenteric fat in both NTG and APP/PS1 mice. All mice were 23-month-old treated with CP2 for 13 - 14 months. Significant changes were determined by two-way ANOVA with Fisher`s LSD *post-hoc* test. Data are mean ± S.E.M.; *n* = 15 - 20 mice *per* group. *P < 0.05, **P < 0.01, ***P < 0.001.

**Extended Data Fig. 4.**
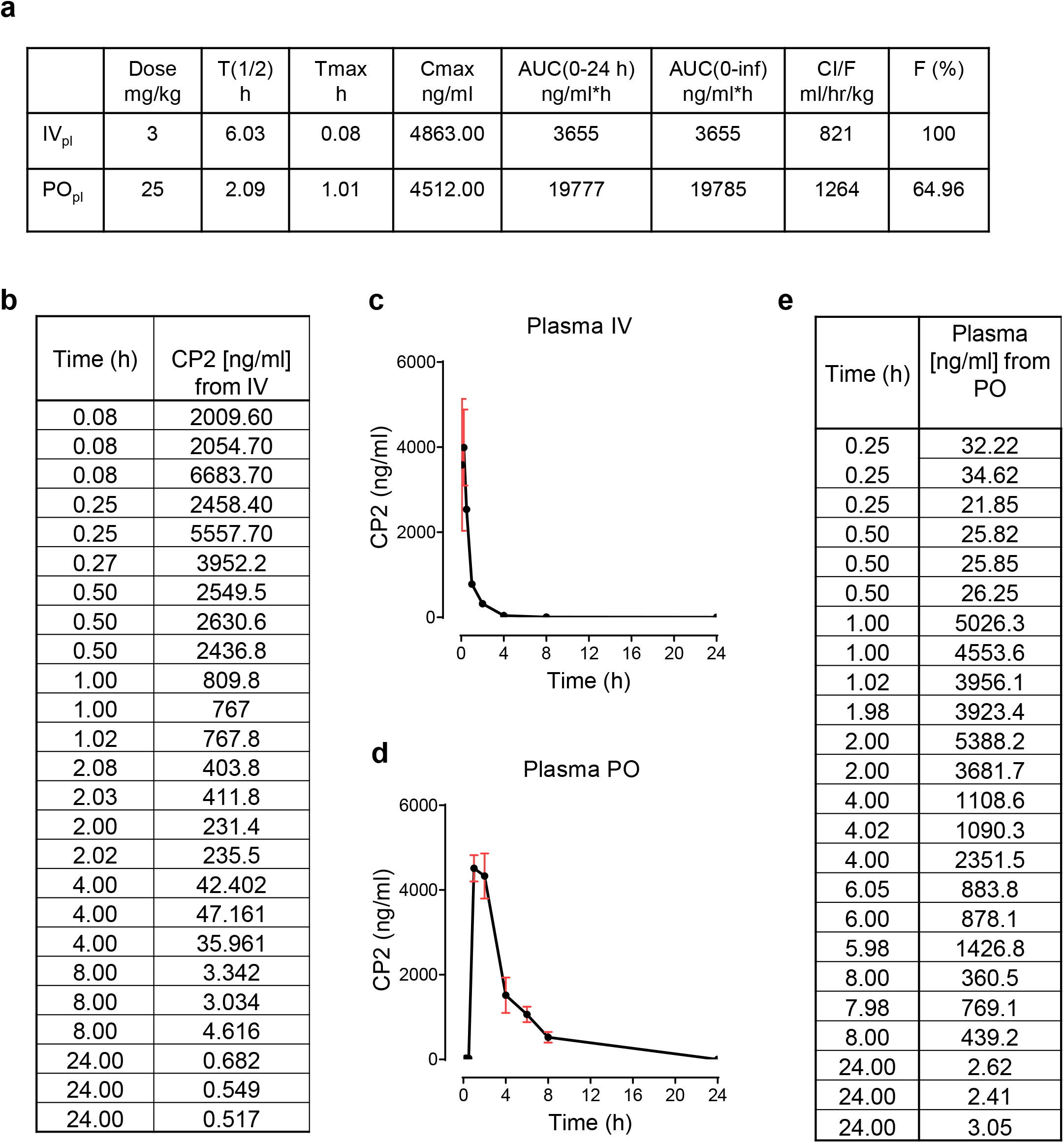
Pharmacokinetics (PK) of CP2 delivered via oral (PO) or intravenous (IV) administration. **a,** Summary of PK study of CP2 delivered to C57BL/6 female mice (*n* = 3 *per* time point) via IV route (3 mg/kg) or via gavage (PO route) (25 mg/kg). The maximal plasma concentration following PO route was 4,512 ng/ml. The T1/2 values following drug administration in plasma were around 6 and 2 h for IV and PO, respectively. The bioavailability (F) was estimated at 65%. AUC, areas under the curve**. b, c**, CP2 levels in plasma after the IV administration (3 mg/kg,). **d,e**, CP2 levels in plasma after the PO administration (25 mg/kg via gavage). Tmax, time to reach maximum (peak) plasma concentration following drug administration; Cmax, maximum (peak) plasma drug concentration; AUC(0-24 h), area under the plasma concentration-time curve over the last 24-h dosing interval; AUC(0-inf), area under the plasma concentration-time curve to infinity represents the total drug exposure across time; CI/F, apparent total clearance of the drug from plasma after oral administration.

**Extended Data Fig. 5.**
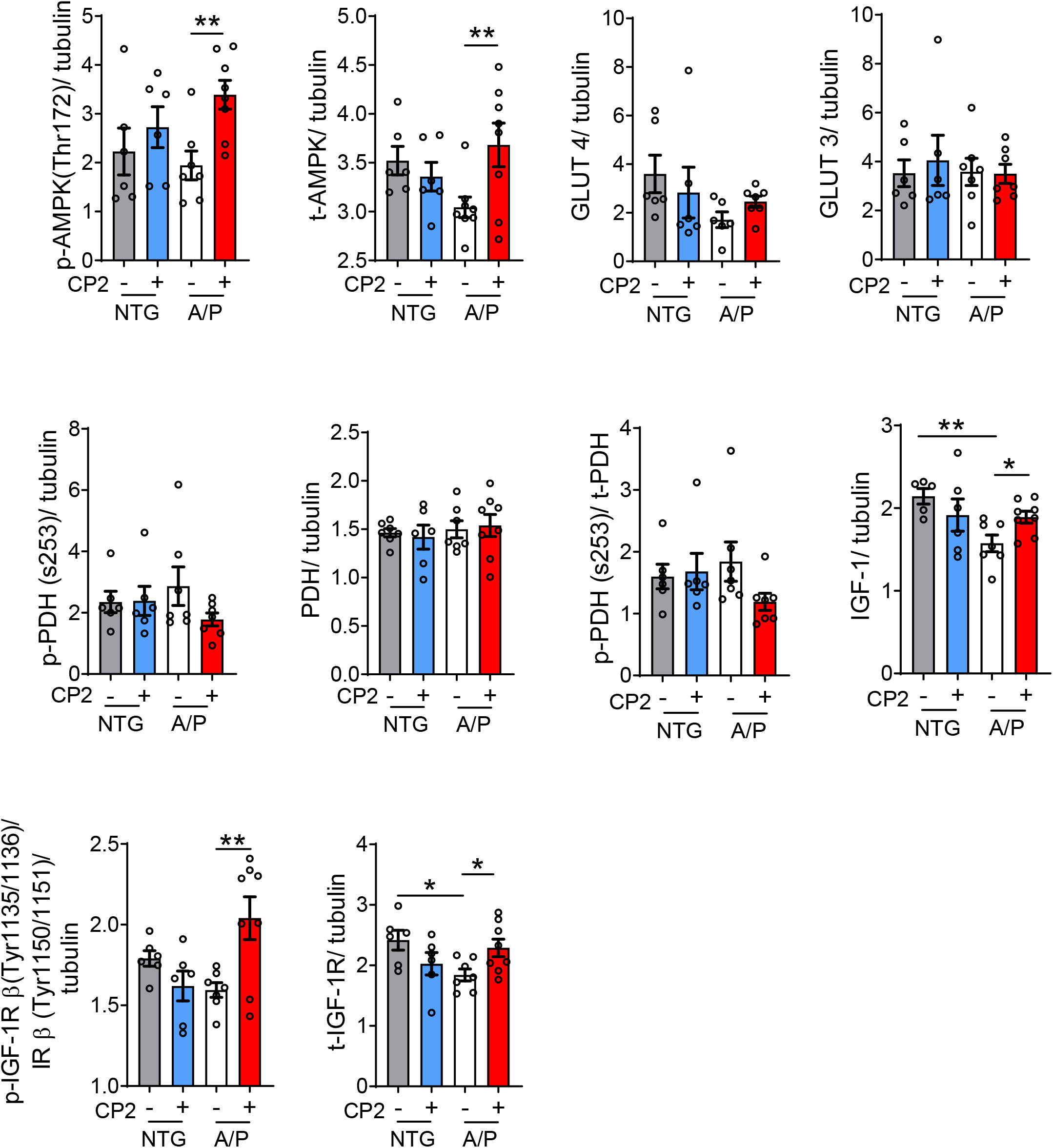
CP2 treatment improves glucose uptake and utilization in APP/PS1 mice. Western blot quantification of protein levels relevant to IGF-1 signaling pathway in the brain tissue of CP2- and vehicle-treated APP/PS1 and NTG mice after 13 - 14 months of treatment. Significance was determined by two-way ANOVA with Fisher`s LSD *post-hoc* test. A/P, APP/PS1; NTG, non-transgenic littermates. Data are presented as mean ± S.E.M. *P < 0.05; **P < 0.01; *n* = 6 - 8 mice *per* group.

**Extended Data Fig. 6.**
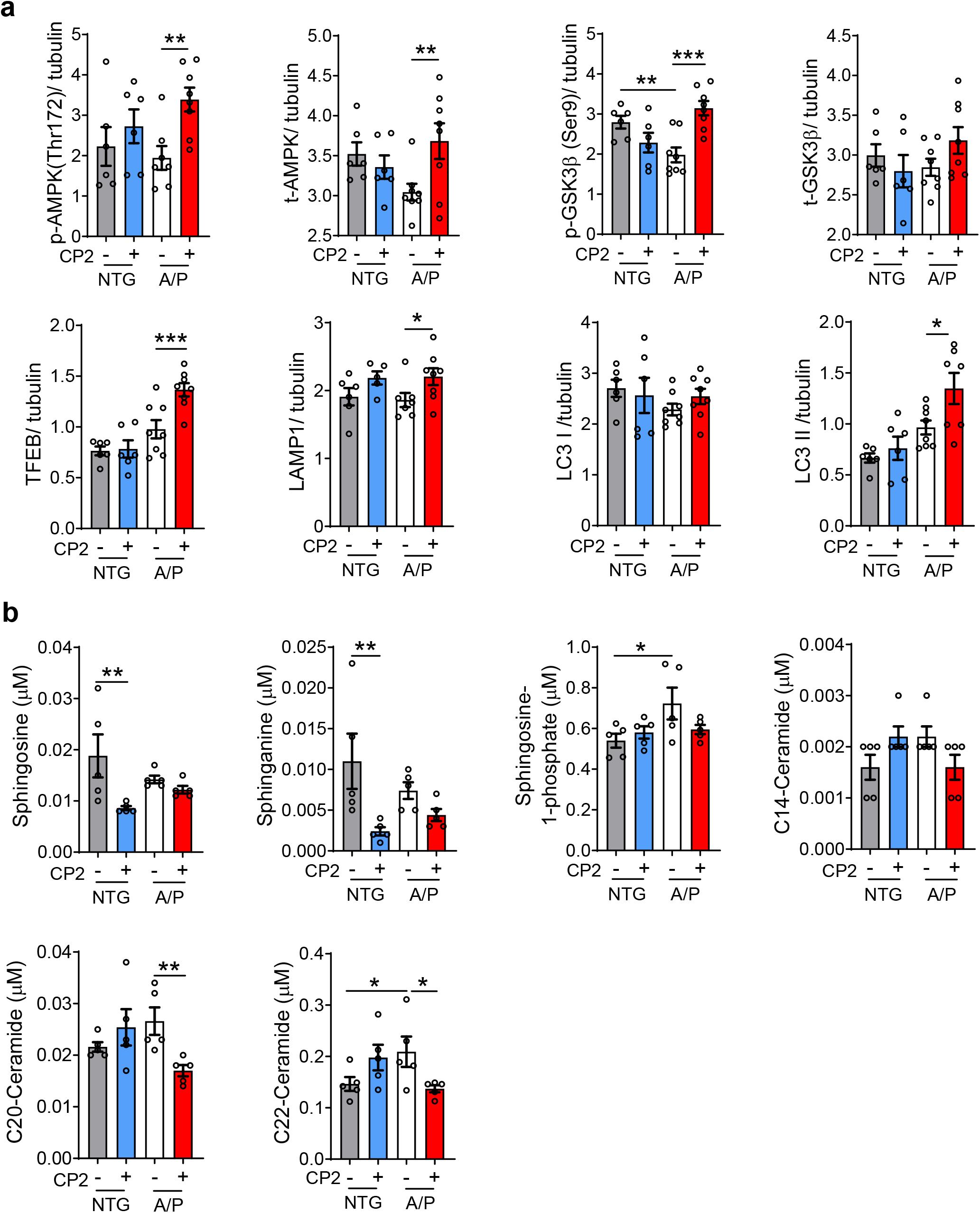
CP2 treatment induces autophagy in the brain and reduces levels of plasma ceramides in symptomatic APP/PS1 mice. **a,** CP2-dependent activation of AMPK and inhibition of GSK3β activates TFEB and increases autophagy in CP2-treated APP/PS1 mice. Quantification of Western blot analysis performed in the brain tissue of 23-month-old NTG and APP/PS1 mice treated with vehicle or CP2. *n* = 6 - 8 mice *per* group. **b,** Plasma from vehicle- and CP2-treated NTG and APP/PS1 mice 23-month-old (*n* = 5 *per* group) was obtained via retro-orbital bleeding. Ceramide concentrations were determined using targeted metabolomics approach and electrospray ionization mass spectrometry ESI/MS. Significance was determined by two-way ANOVA with Fisher`s LSD *post-hoc* test. A/P, APP/PS1; NTG, non-transgenic littermates. Data are presented as mean ± S.E.M. *P < 0.05; **P < 0.01; ***P < 0.001.

**Extended Data Fig. 7.**
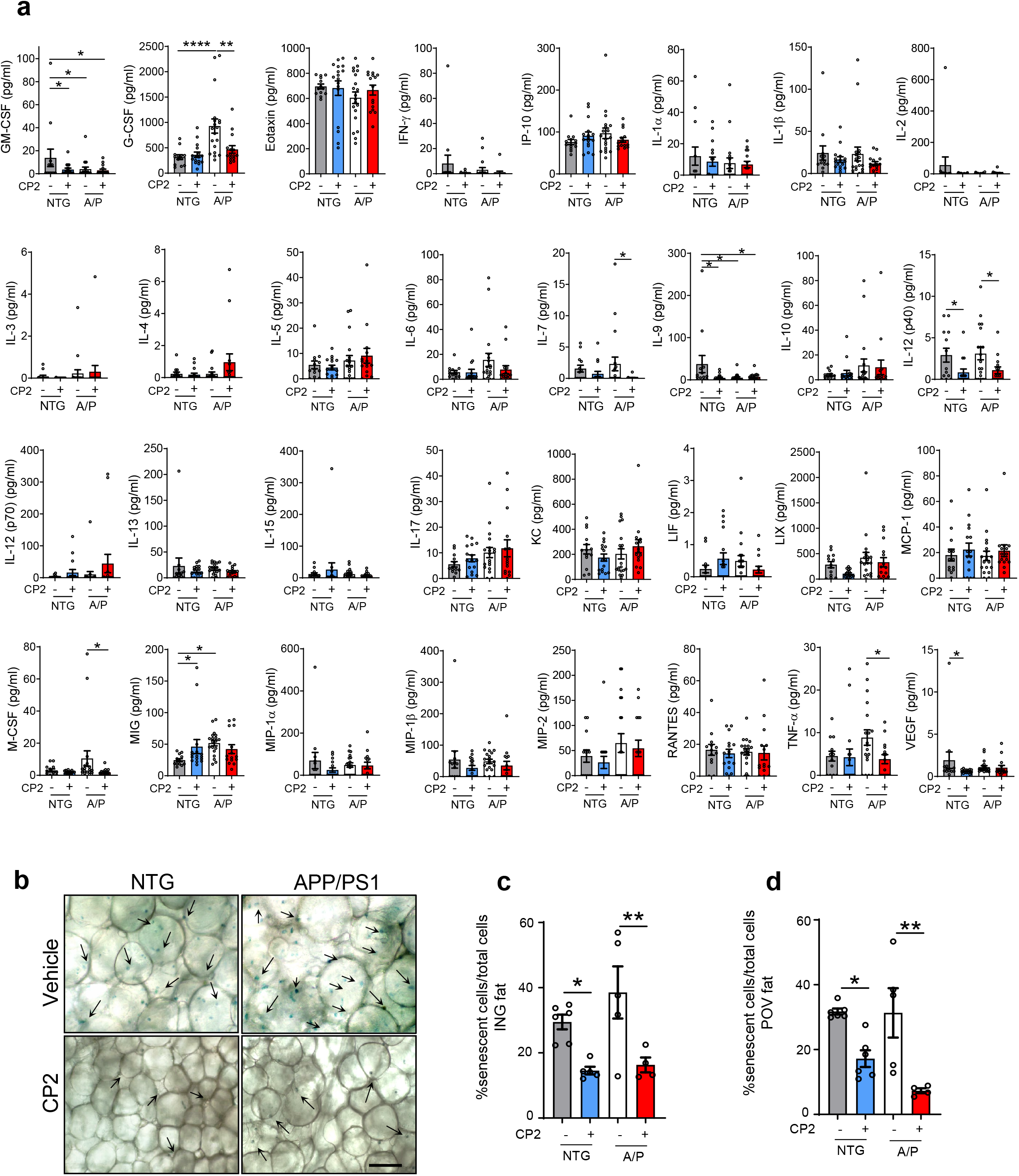
CP2 treatment reduces inflammatory markers in blood and level of senescence cells in adipose tissue of symptomatic APP/PS1 and NTG mice. **a,** A panel of 32 cytokines and chemokines measured in plasma from 23-month-old symptomatic APP/PS1 and NTG vehicle- and CP2-treated mice (*n* = 15 – 20 mice *per* group). Plasma was obtained via retro-orbital bleeding; ELISA was conducted by Eve Technologies. **b,** Representative images of β-Gal staining of senescence cells in periovarian (POV) and inguinal (ING) adipose tissue from 23-month-old vehicle and CP2-treated NTG and APP/PS1 mice. Scale bar, 100μm. **c,d,** Quantification of β-Gal staining in ING (**c**) and POV (**d**) fat depots from (**b**). *n* = 5 - 6 mice *per* group. A two-way ANOVA with Fisher`s LSD *post-hoc* test was used for statistical analysis. Data are presented as mean ± S.E.M. *P < 0.05; **P < 0.01; ****P < 0.0001.

**Extended Data Fig. 8.**
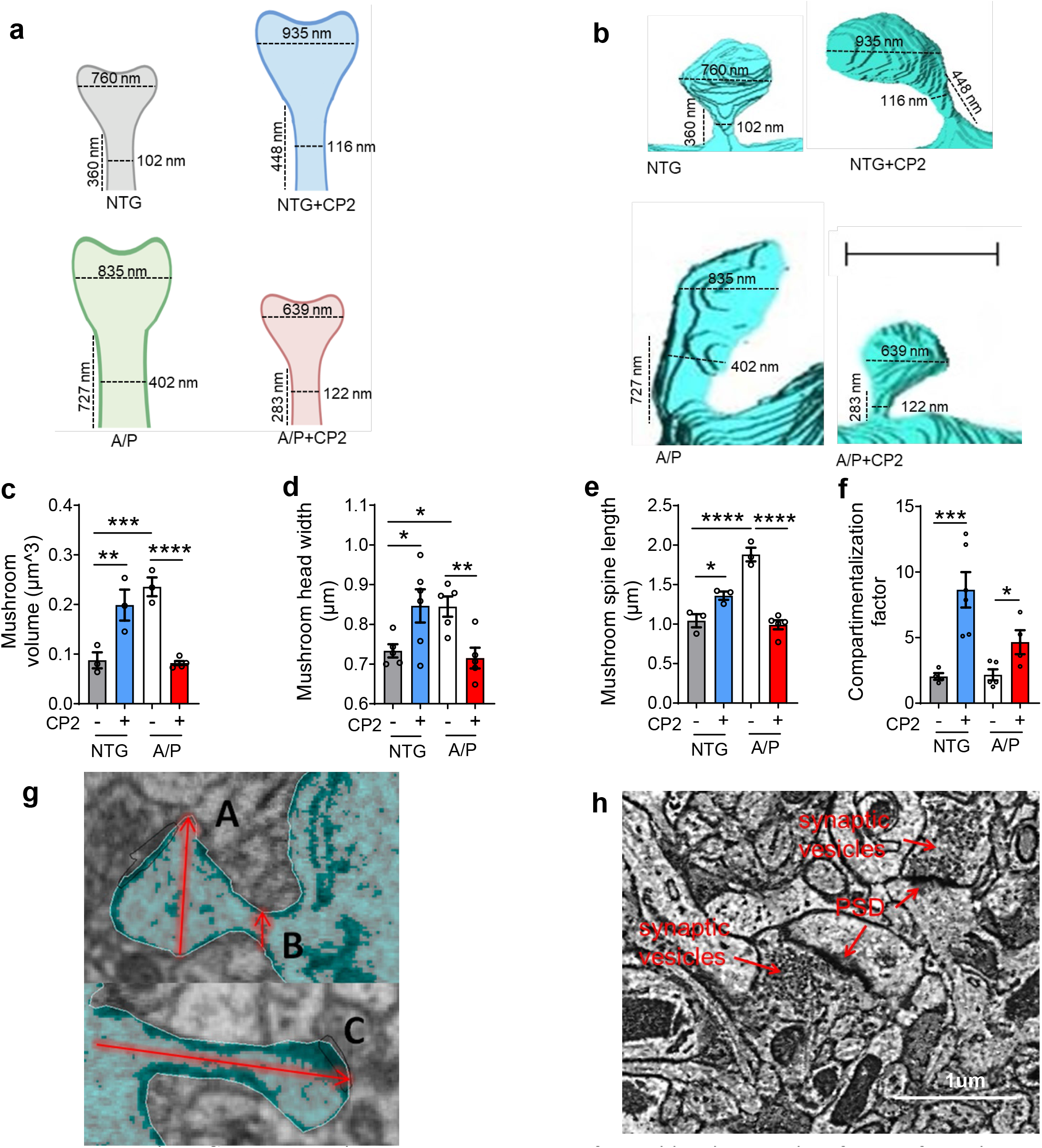
CP2 treatment improves the morphology of dendritic spines required for LTP formation and synaptic function. **a**, Schematic illustration of morphological alterations in dendritic spines observed in CP2-treated NTG and APP/PS1 mice. Numbers indicate mean values. **b**, Representative images of mushroom spines generated using 3DEM from brain tissue of vehicle- and CP2-treated NTG and APP/PS1 mice. Scale bar, 1 μm. APP/PS1 mice have larger spine heads and longer and wider spine necks, which is associated with reduced ability to maintain LTP. **c-e**, Multiple parameters of dendritic spine morphology depicted in (**a,b**) measured in vehicle- and CP2-treated NTG and APP/PS1 mice including the volume of mushroom spines (**c**); mushroom head width (**d**); and mushroom spine length (**e**). Increased spine maturation after CP2 treatment is supported by measurements of the overall spine volume and spine geometry, especially the length and the width of the neck and the head, which are essential parameters influencing the ability of spines to retain synaptic receptors. **f**, The compartmentalization factor, a measure of biochemical compartmentalization of spine synapses (see definition in the Method), calculated for each mushroom spine in vehicle *vs.* CP2-treated NTG and APP/PS1 mice. Significant increase in compartmentalization factor in CP2-treated mice indicates stronger ability to maintain LTP. **g**, Example of measurements of head diameter (A), neck width (B), and spine length from base to head (C). **h**, Representative EM micrograph of the CA1 hippocampal region utilized in the calculation of active synapses. Identification of synapses was done based on the presence of postsynaptic density (PSD) and synaptic vesicles. Scale bar, 1 μm. Data are presented as mean ± S.E.M. A two-way ANOVA with Fisher`s LSD *post-hoc* test was used for statistical analysis. *P < 0.05; **P < 0.01; ***P < 0.001; ****P < 0.0001.

**Extended Data Fig. 9.**
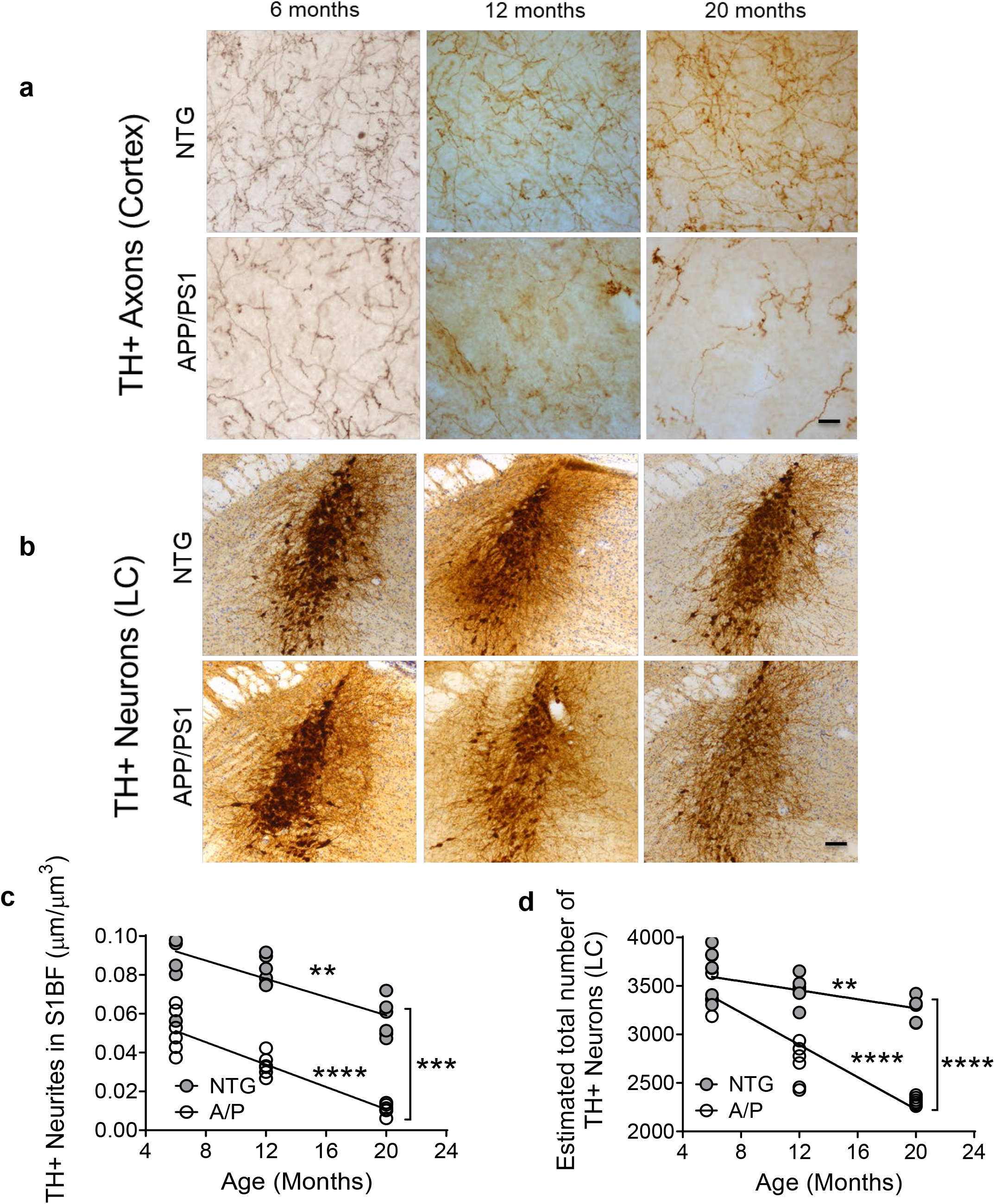
APP/PS1 mice display progressive neurodegeneration of TH+ neurons in cortex and *Locus Coeruleus* (LC). NTG and APP/PS1 mice were evaluated for the integrity of TH+ axons in S1BF (**a,c**) and TH+ neurons in LC (**b,d**) at 6, 12, and 20 months of age. Representative images of TH+ axons in S1BF (**a**) and TH+ neurons in LC (**b**). **c,** Progressive loss of TH+ axons was observed in both APP/PS1 (P < 0.0001) and NTG mice (P < 0.01). The rate of age-related decline of TH+ axons was similar between NTG and APP/PS1 (differences between slopes are not significant P = 0.52), but magnitude of TH+ axonal loss was greater in APP/PS1 mice (differences between the slopes elevation are significant P = 0.001). **d,** The age-related loss of TH+ neurons was observed starting from 6 months of age in both NTG (P < 0.01) and APP/PS1 mice (P < 0.0001), and additionally APP/PS1 mice showed greater age-related loss of TH+ neurons (P < 0.0001). *n* = 5 – 6 female mice *per* group in each test. Data are represented as mean ± S.E.M. A linear regression analysis was applied for statistical comparison between the groups. **P < 0.01, ***P < 0.001, ****P < 0.0001. Scale bars: **a**, 20 μm; **b**, 100 μm.

**Extended Data Fig. 10.**
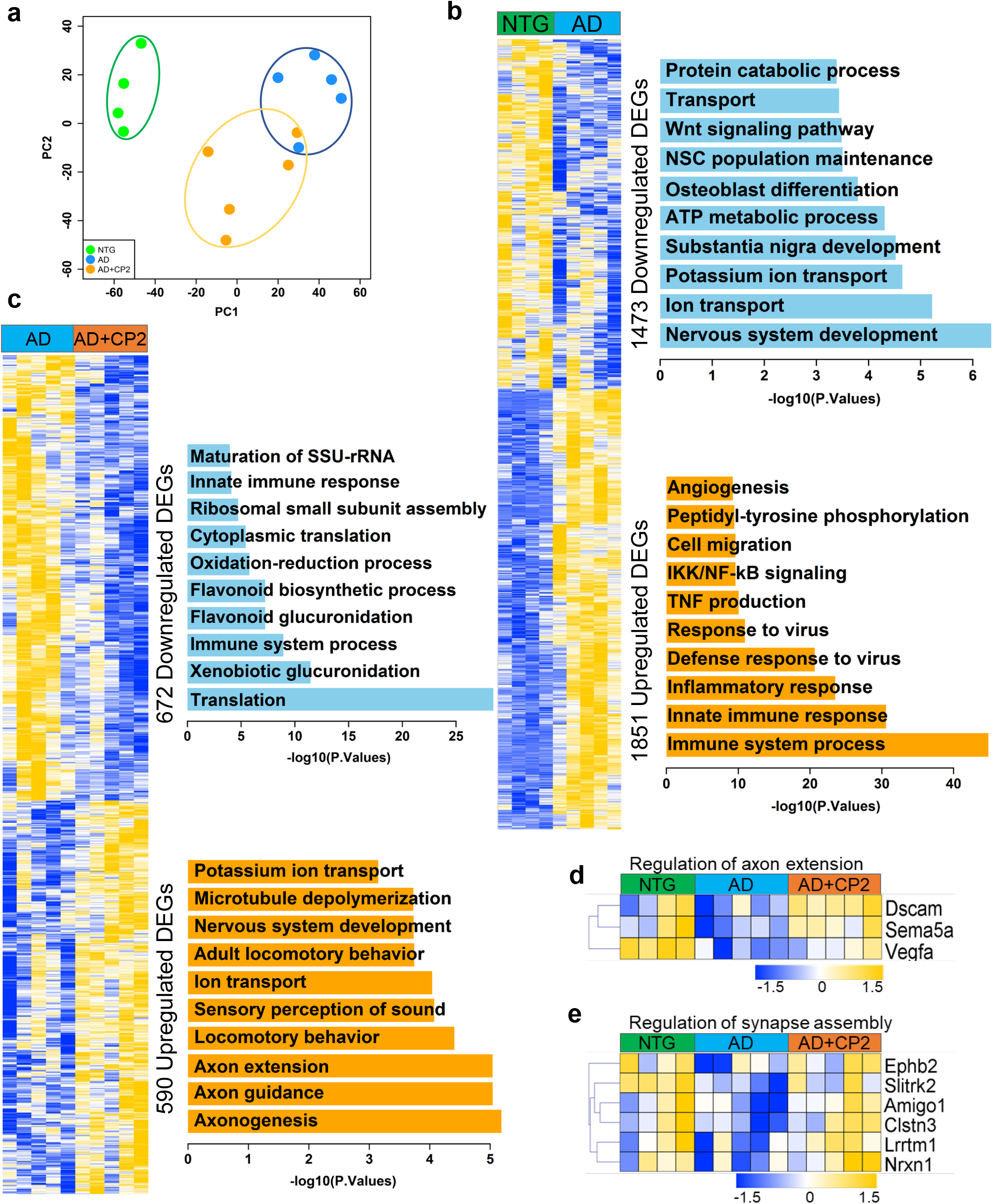
Transcriptomic changes in brain tissue from vehicle and CP2-treated NTG and APP/PS1 mice determined using RNA-seq. **a,** Principal Component Analysis (PCA) shows separated clusters of samples among 3 groups (NTG, green; APP/PS1, blue; APP/PS1+CP2, orange) where CP2 treatment produces specific changes allowing group separation. **b,** Differentially expressed genes in NTG *vs.* APP/PS1 mice and function enrichment analysis based on up- or down-regulated genes. **c,** Differentially expressed genes in APP/PS1 *vs.* APP/PS1+CP2 mice and function enrichment analysis based on up- or down-regulated genes. **d,** A heatmap shows changes in genes of pathway related to the positive regulation of axon extension, which were up-regulated after CP2 treatment in APP/PS1 mice. **e,** A heatmap shows changes in genes involved in synapse assembly that were up-regulated by CP2 treatment in APP/PS1 mice. All mice were 23 months of age treated with CP2 or vehicle for 13-14 months. *n* = 4 - 5 mice *per* group.

## Supplementary Information

**Supplementary Fig. 1. Original uncropped Western blot images for Extended Data Fig. 1e-l**. Time course of the expression of key proteins in each of the AMPK-dependent neuroprotective pathways depicted in Extended Data Fig. 1d in the brain tissue of symptomatic APP/PS1 mice 9 months of age acutely gavaged with 25 mg/kg of CP2. One mouse was taken for each time point. Blue arrows indicate the bands shown in Extended Data Fig. 1.

**Supplementary Fig. 2. Original uncropped Western blot images for Fig. 1r.** Cortico-hippocampal region of 6 - 8 mice *per* group was taken for Western blot analysis (NTG, *n* = 6; NTG+CP2, *n* = 6; APP/PS1, *n* = 8; APP/PS1+CP2, *n* = 8).

**Supplementary Fig. 3. Original uncropped Western blot images for Fig. 2g.** Cortico-hippocampal region of 6 - 8 mice *per* group was taken for Western blot analysis (NTG, *n*= 6; NTG+CP2, *n* = 6; APP/PS1, *n* = 8; APP/PS1+CP2, *n* = 8).

**Supplementary Fig. 4. Original uncropped Western blot images for Fig. 3k.** Cortico-hippocampal region of 6 - 8 mice *per* group was taken for Western blot analysis (NTG, *n* = 6; NTG+CP2, *n* = 6; APP/PS1, *n* = 8; APP/PS1+CP2, *n* = 8).

**Supplementary Table 1.** Results of the Eurofin Cerep Safety-Screen 44 Panel (receptors and ion channels).

**Supplementary Table 2.** Results of the Eurofin Cerep Safety-Screen 44 Panel (enzymes).

**Supplementary Table 3**. Results of kinome profiling for CP2 in the Nanosyn 250 Kinase panel.

**Supplementary Table 4.** CP2 levels in the brain tissue and plasma of NTG and APP/PS1 mice 22 – 24-month-old chronically treated from 9 months of age.

**Supplementary Table 5.** Effect of CP2 treatment on brain metabolite levels in APP/PS1 and NTG mice.

**Supplementary Table 6.** List of DEGs in APP/PS1 *vs.* NTG gene set comparison

**Supplementary Table 7.** The Gene Ontology enrichment analysis of down-regulated processes in APP/PS1 *vs.* NTG mice.

**Supplementary Table 8.** The Gene Ontology enrichment analysis of up-regulated processes in APP/PS1 *vs.* NTG mice.

**Supplementary Table 9.** List of DEGs in APP/PS1 v*s.* APP/PS1+CP2 gene set comparison.

**Supplementary Table 10.** The Gene Ontology enrichment analysis of down-regulated processes in APP/PS1 *vs.* APP/PS1+CP2 comparison.

**Supplementary Table 11.** The Gene Ontology enrichment analysis of up-regulated processes in APP/PS1 *vs.* APP/PS1+CP2 gene set comparison

**Supplementary Table 12.** Overlapped DEGs in APP/PS1+CP2 *vs.* APP/PS1 and APP/PS1 *vs.* NTG gene set comparison.

**Supplementary Table 13.** The Gene Ontology enrichment analysis of functional changes down-regulated by CP2 treatment in APP/PS1 mice to the levels detected in NTG mice.

**Supplementary Table 14.** The Gene Ontology enrichment analysis of functional changes up-regulated by CP2 treatment in APP/PS1 mice to the levels detected in NTG mice.

**Supplementary Table 15.** List of down-regulated and up-regulated genes identified in comparison of female AD patients and cognitively normal controls in the AMP-AD data set.

**Supplementary Table 16.** List of 294 down-regulated DEGs in comparison between NTG *vs.* APP/PS1 mice matched to down-regulated genes identified in the female AMP-AD cohort.

**Supplementary Table 17**. The Gene Ontology enrichment analysis of down-regulated functional processes in both APP/PS1 mice and females with AD in AMP-AD cohort.

**Supplementary Table 18**. List of 518 up-regulated DEGs in NTG vs. APP/PS1 mice matching to down-regulated genes identified in the female AMP-AD cohort.

**Supplementary Table 19.** The Gene Ontology enrichment analysis of up-regulated functional processes in both APP/PS1 mice and female AD patients.

**Supplementary Table 20.** List of 71 up-regulated DEGs in APP/PS1+CP2 *vs.* APP/PS1 and APP/PS1 *vs.* NTG gene set comparison matched to the up-regulated genes identified in the female AMP-AD cohort.

**Supplementary Table 21.** The Gene Ontology enrichment analysis of functional processes identified in APP/PS1+CP2 *vs.* APP/PS1 and APP/PS1 *vs.* NTG gene set comparison matched to the up-regulated genes identified in the female AMP-AD cohort.

**Supplementary Table 22.** List of 57 down-regulated DEGs in APP/PS1+CP2 *vs.* APP/PS1 and APP/PS1 *vs.* NTG gene set comparison, matched to down-regulated genes identified in the female AMP-AD cohort.

**Supplementary Table 23.** The Gene Ontology enrichment analysis of functional processes identified in APP/PS1+CP2 *vs.* APP/PS1 and APP/PS1 *vs.* NTG gene set comparison matched to the down-regulated genes identified in the female AMP-AD cohort.

