## Supplementary Figures for "Partial inhibition of mitochondrial complex I attenuates neurodegeneration and restores energy homeostasis and synaptic function in a symptomatic Alzheimer’s mouse model"

### Western blots for Extended data Fig. 1

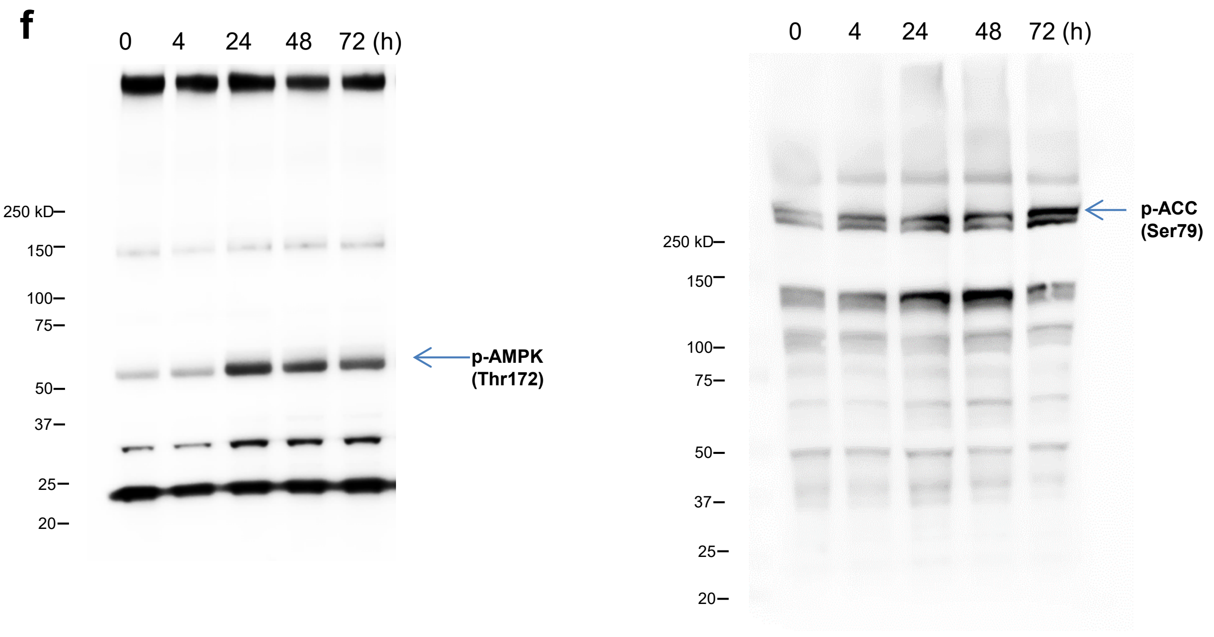


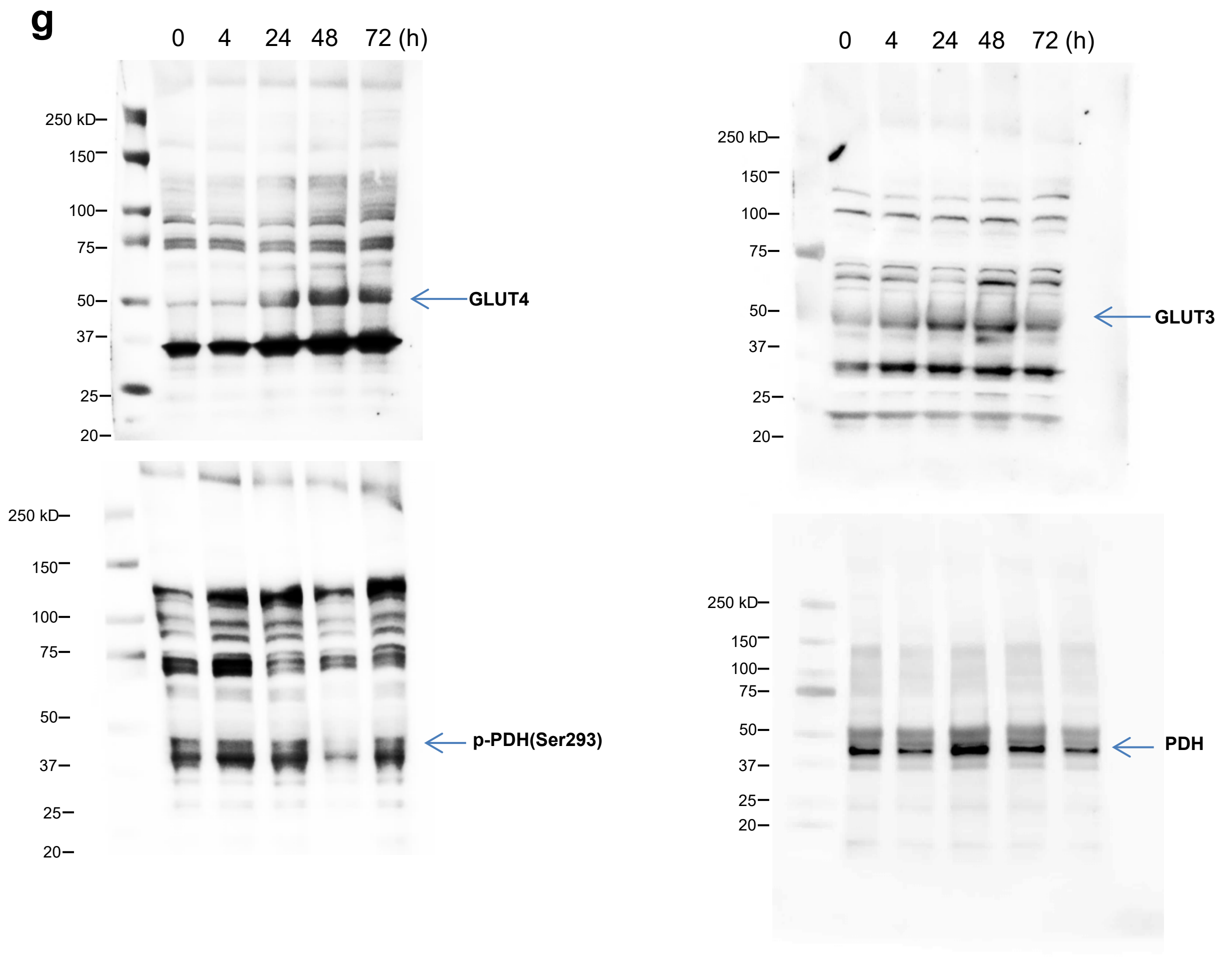

### Western blots for Extended data Fig. 1

i

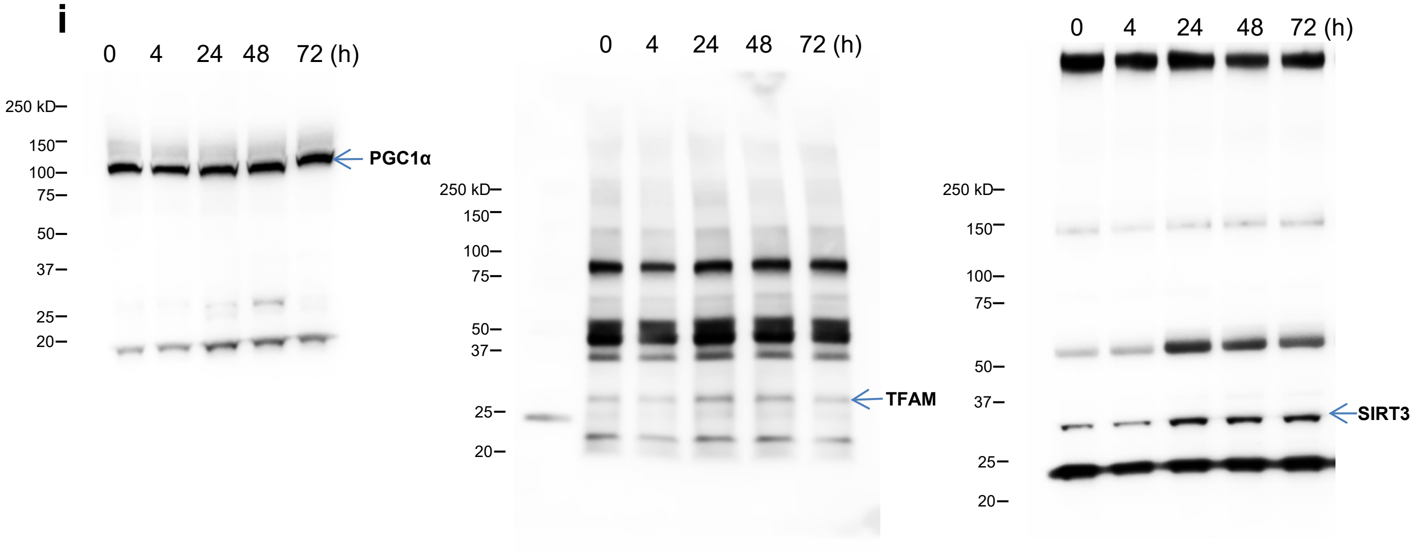


j

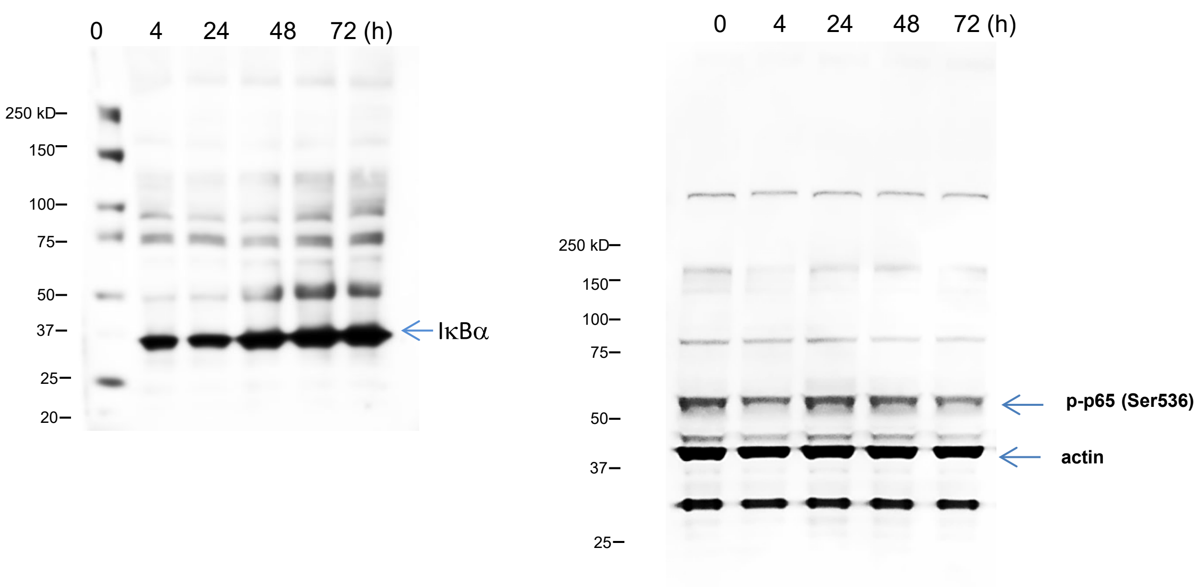


k

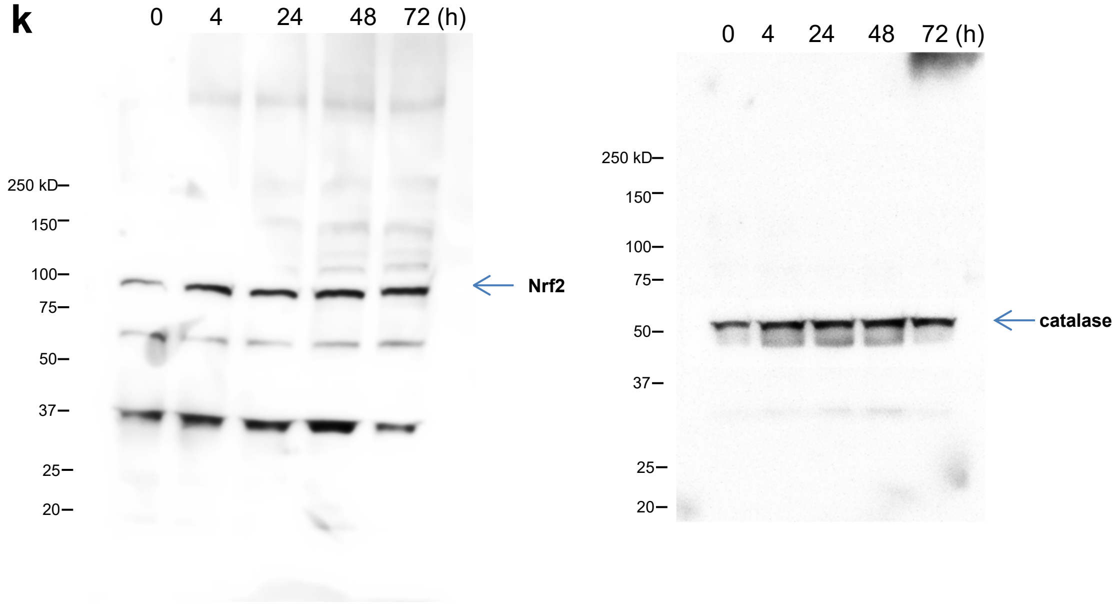

### Western blots for Extended data Fig. 1

**k**

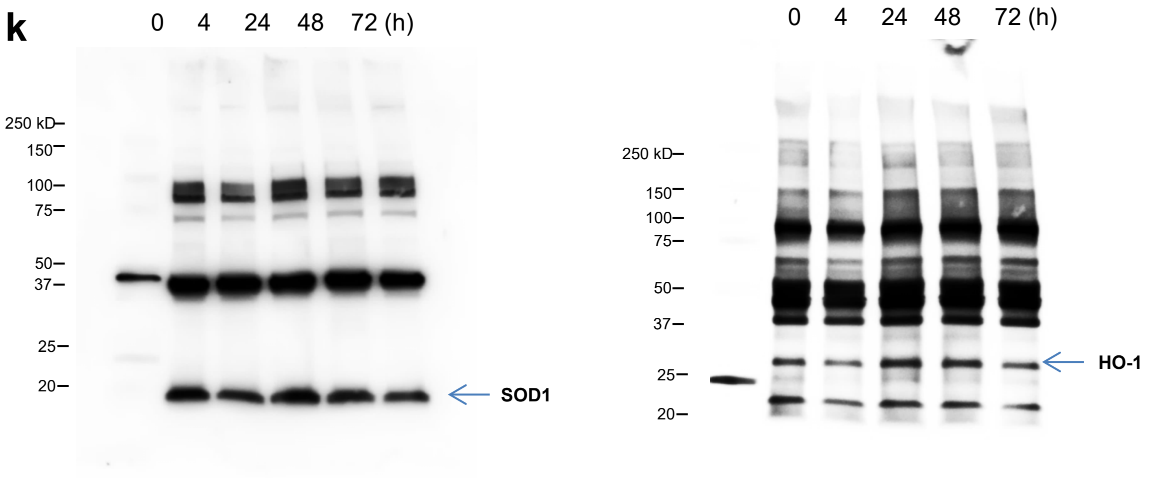


**l**

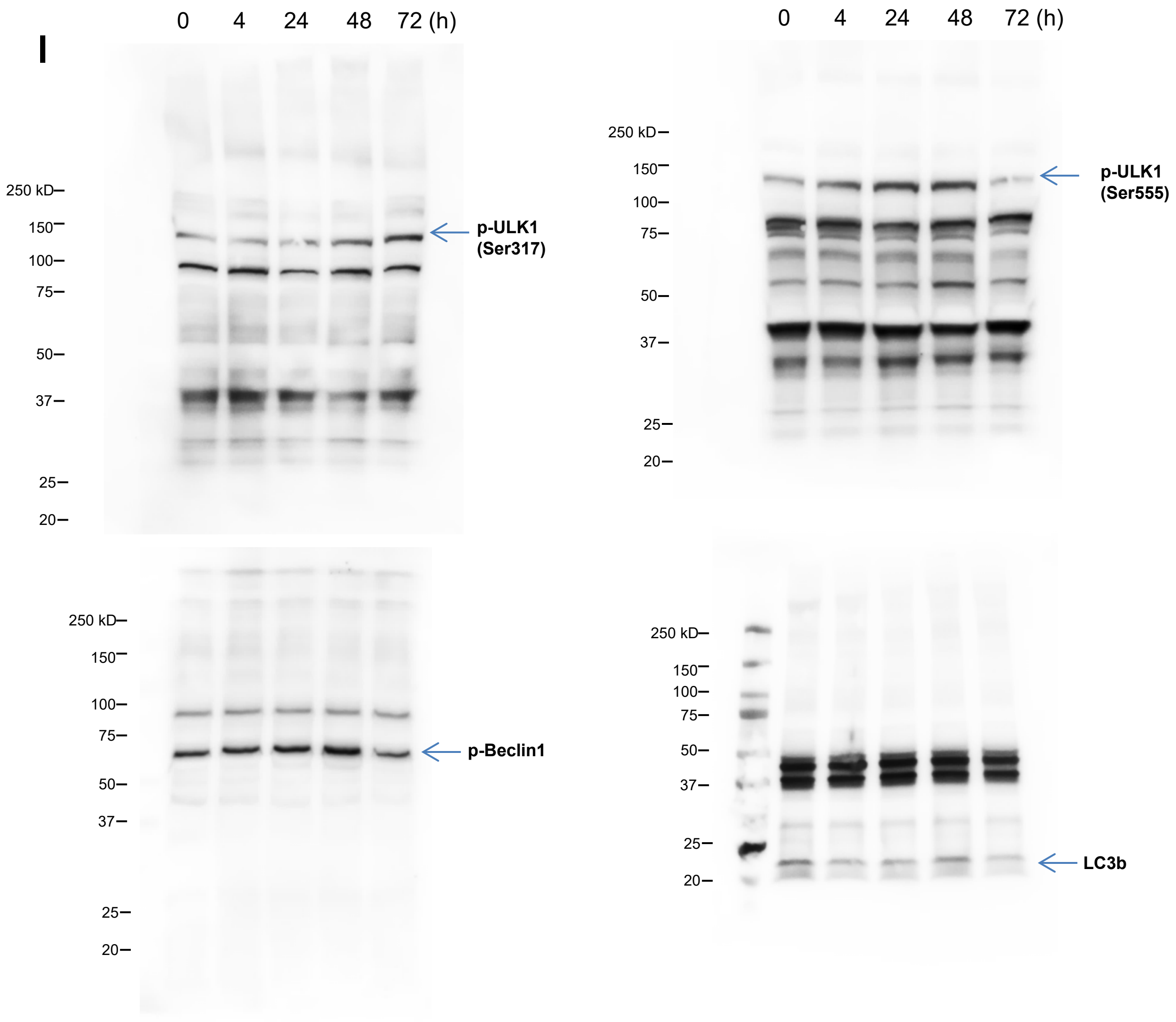


**Supplementary Fig. 1. Original uncropped Western blot images for Extended Data Fig. 1e-l.** Time course of the expression of key proteins in each of the AMPK-dependent neuroprotective pathways depicted in Extended Data Fig. 1d in the brain tissue of symptomatic APP/PS1 mice 9 months of age acutely gavaged with 25 mg/kg of CP2. One mouse was taken for each time point. Blue arrows indicate the bands shown in Extended Data Fig. 1

### Western blots for Fig. 1r

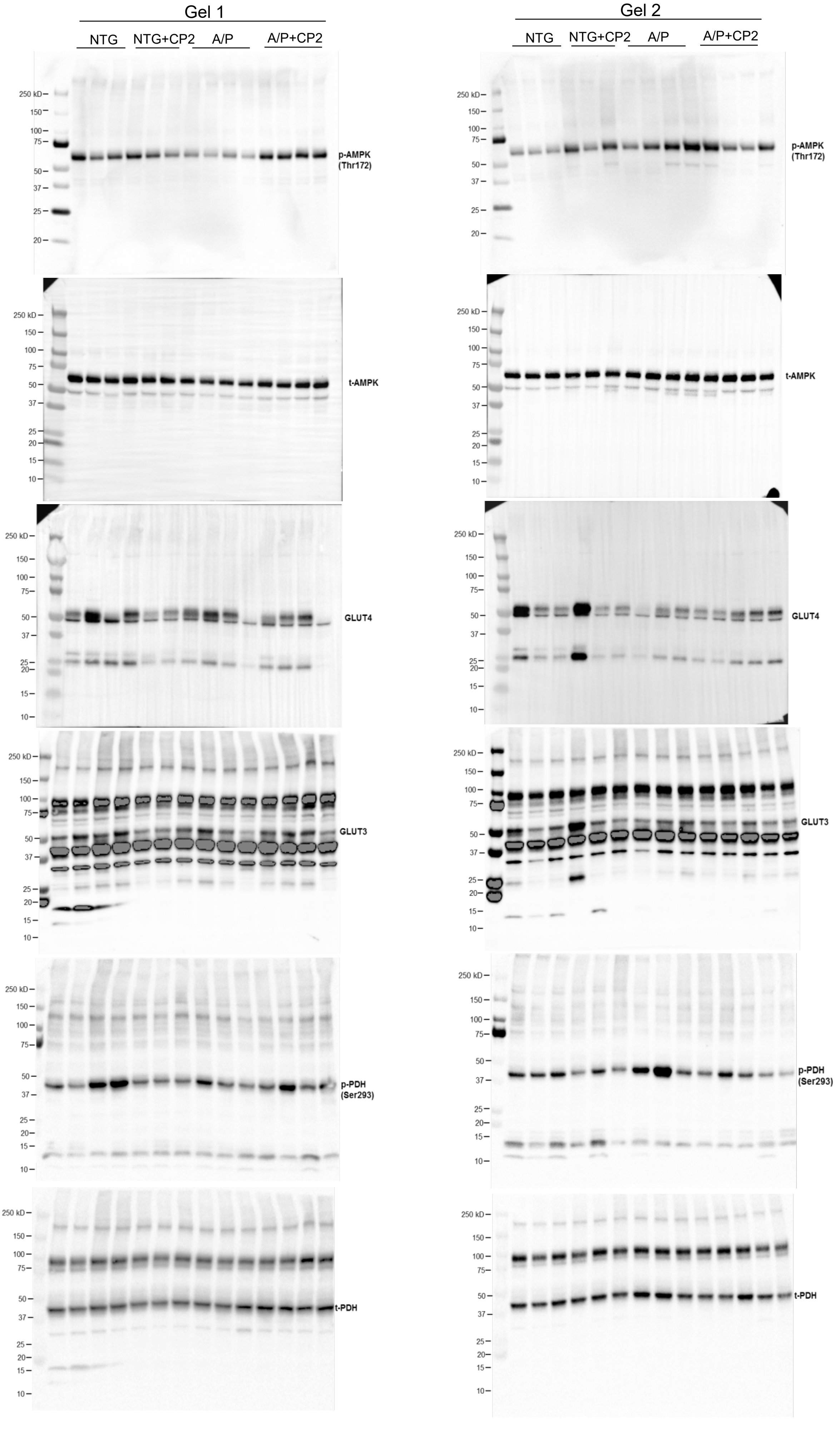

### Western blots for Fig. 1r

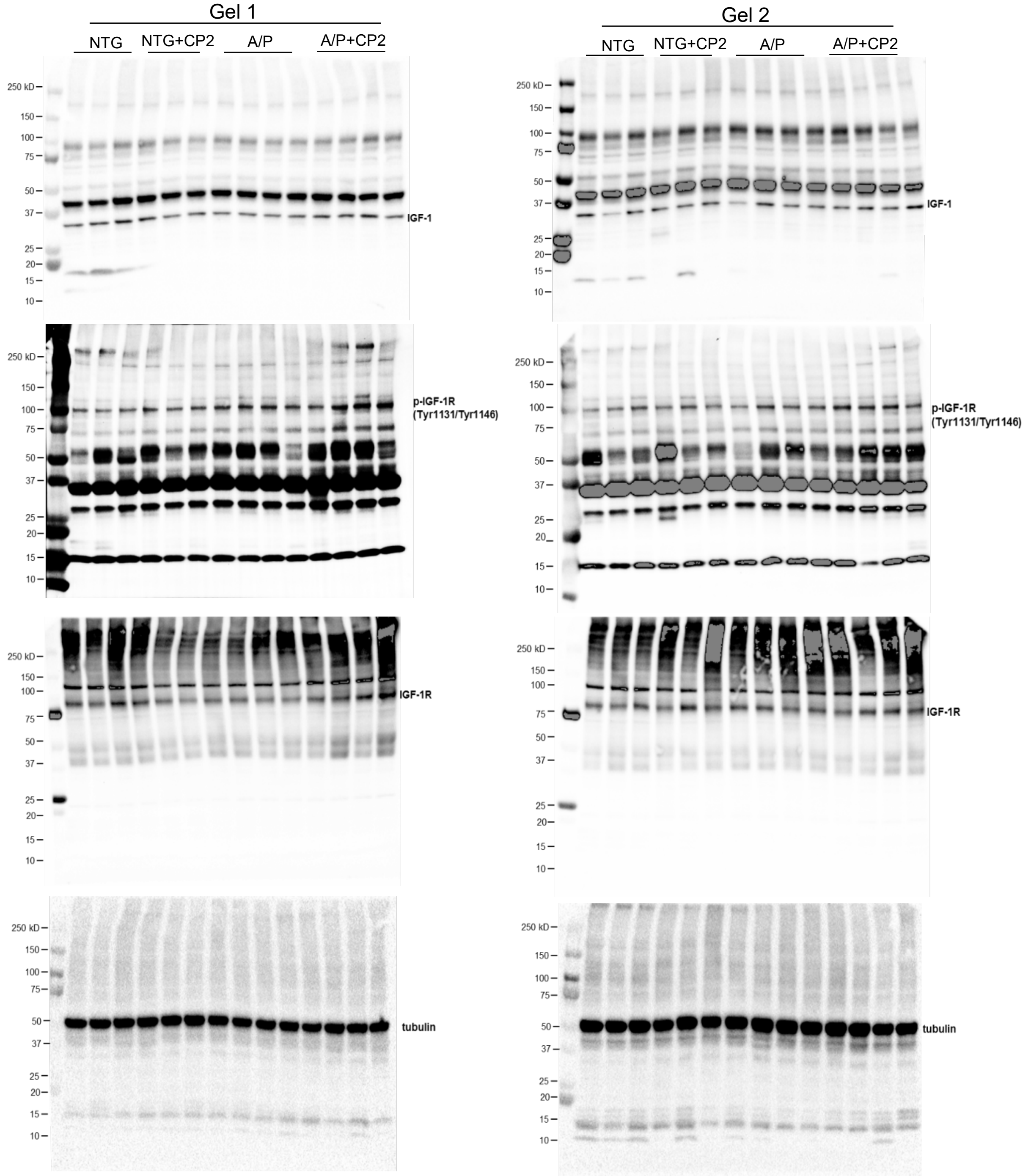


**Supplementary Fig. 2. Original uncropped Western blot images for Fig. 1r.** Cortico-hippocampal region of 6 - 8 mice *per* group was taken for Western blot analysis (NTG, *n* = 6; NTG+CP2, *n* = 6; APP/PS1, *n* = 8; APP/PS1+CP2, *n* = 8).

Western blots for Fig. 2g

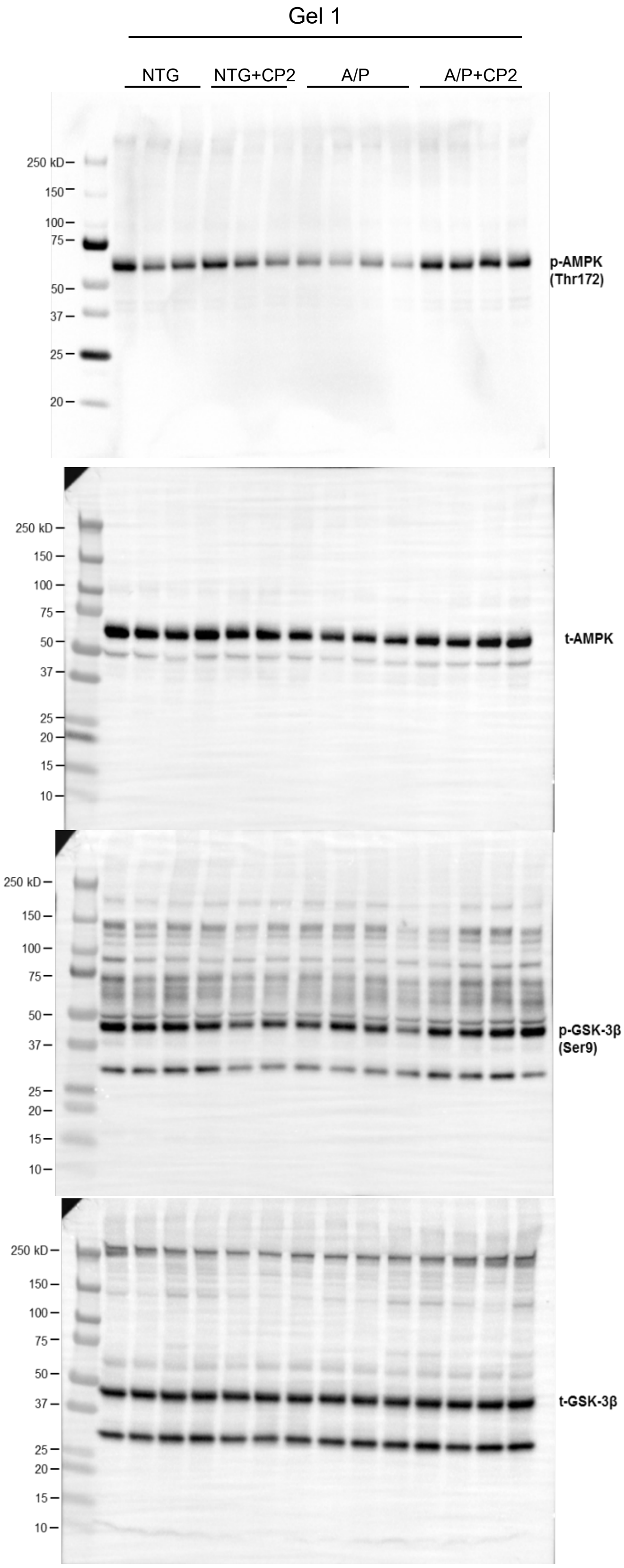


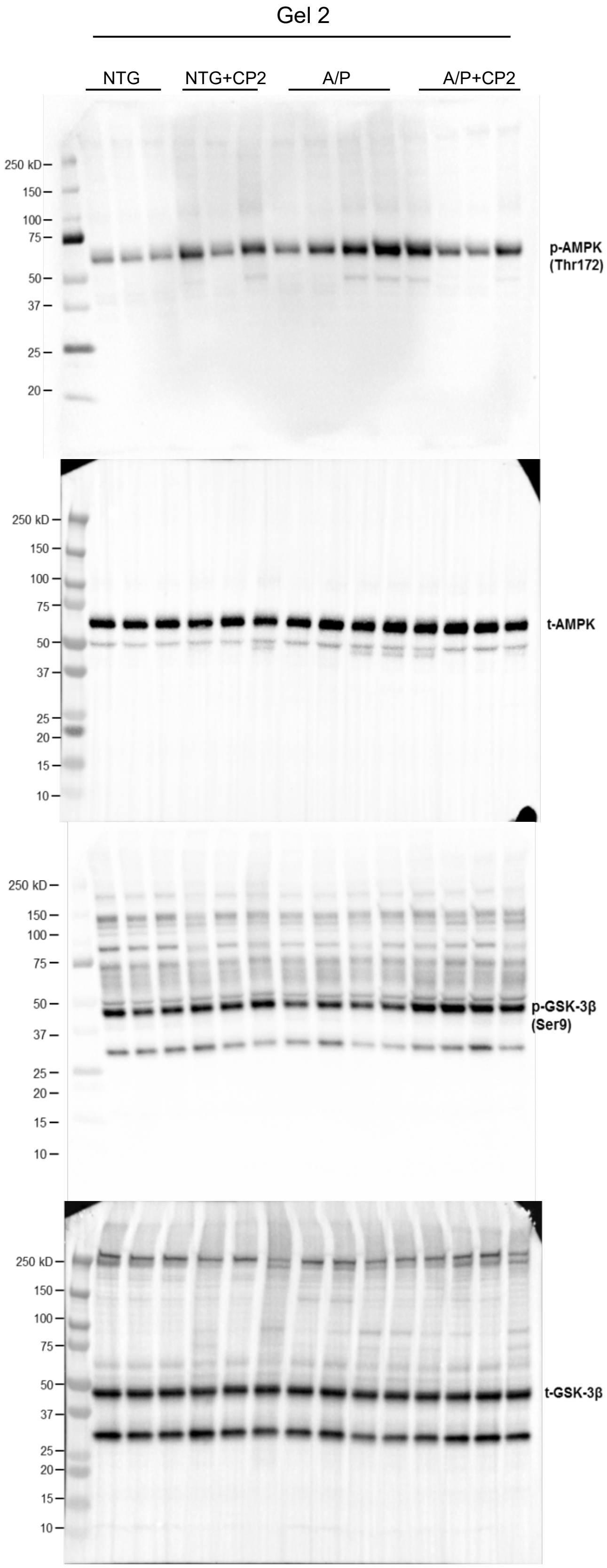

### Western blots for Fig. 2g

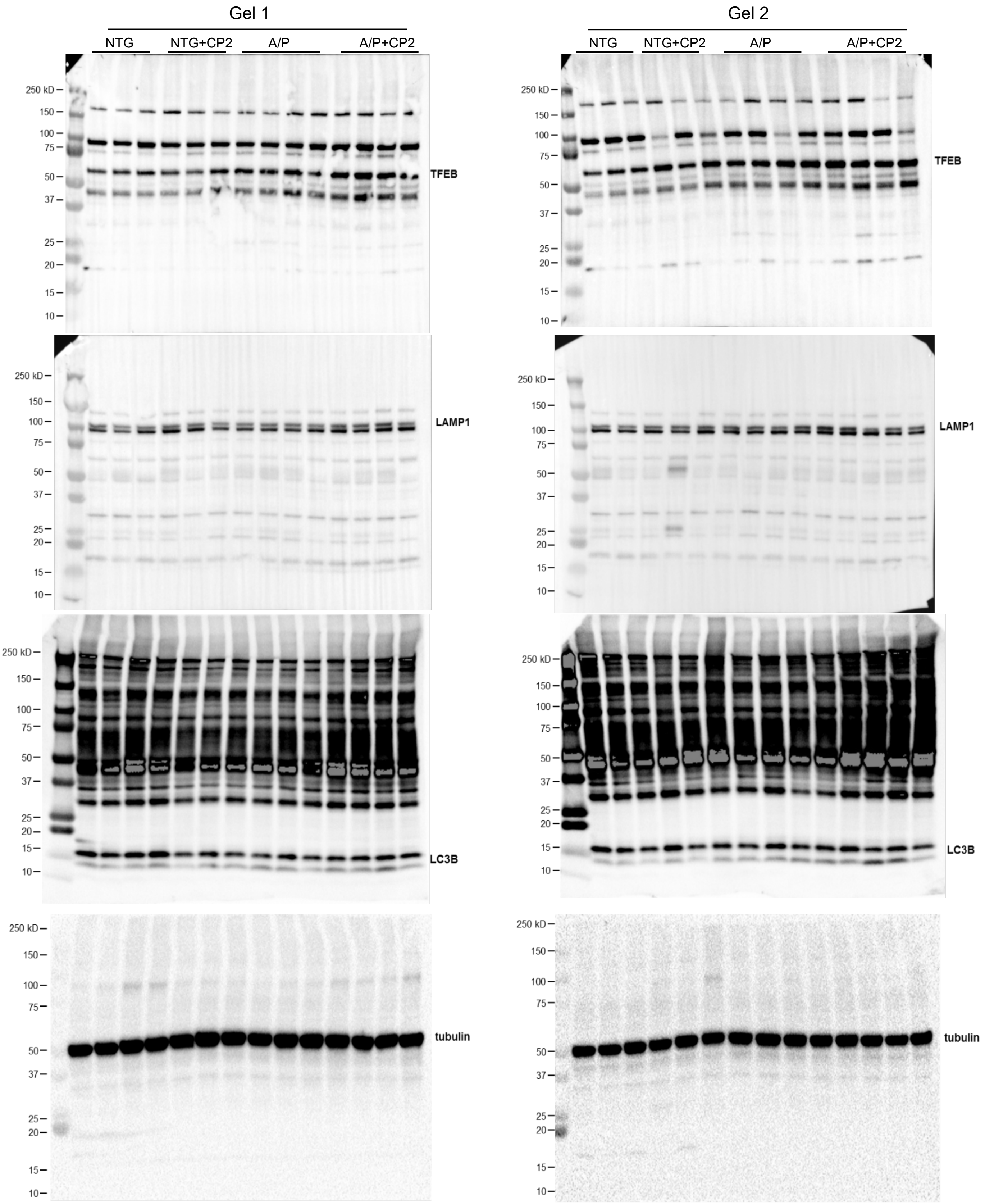


**Supplementary Fig. 3. Original uncropped Western blot images for Fig. 2g.** Cortico-hippocampal region of 6 - 8 mice *per* group was taken for Western blot analysis (NTG, *n*= 6; NTG+CP2, *n* = 6; APP/PS1, *n* = 8; APP/PS1+CP2, *n* = 8).

### Western blots for Fig. 3k

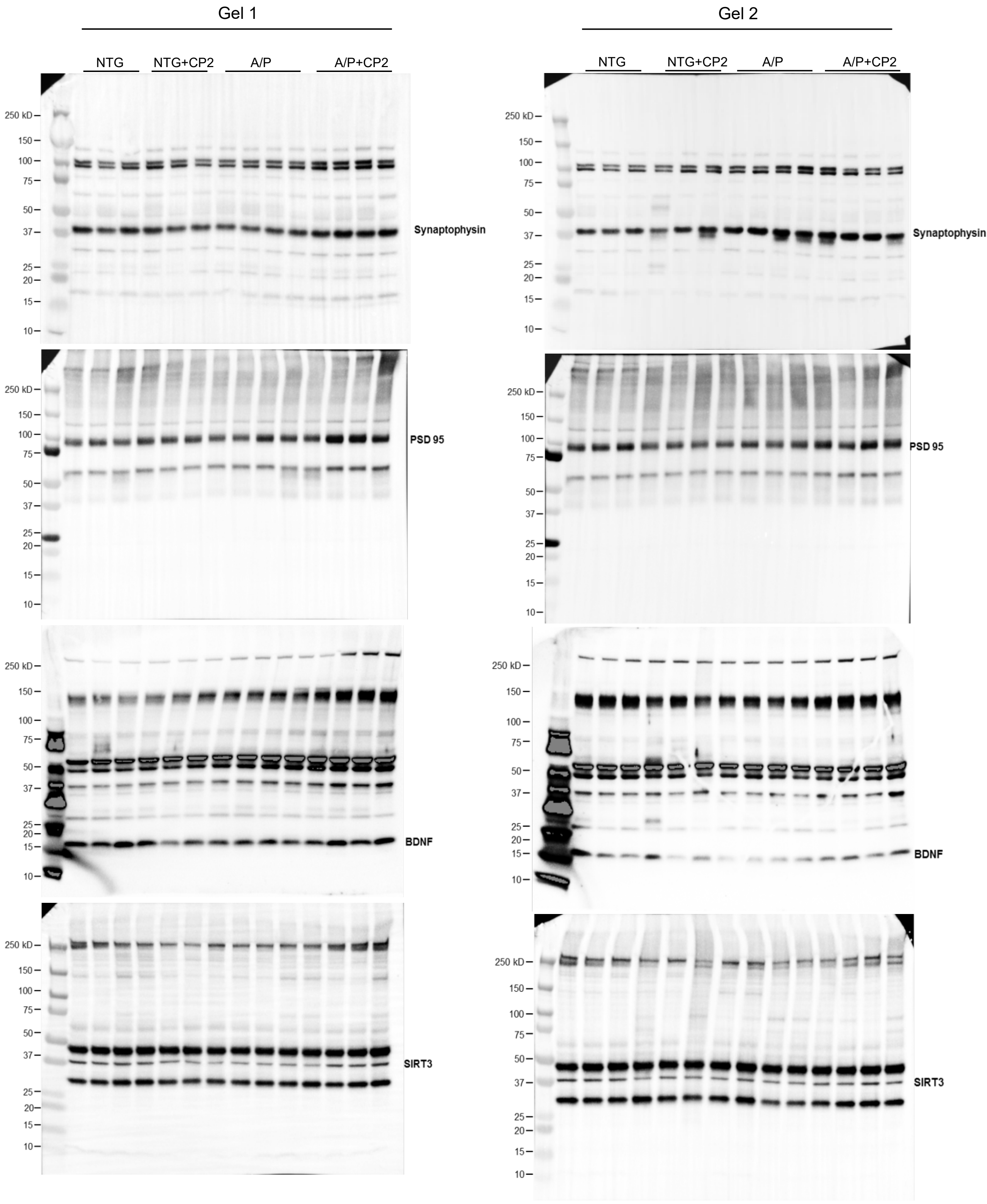

### Western blots for Fig. 3k

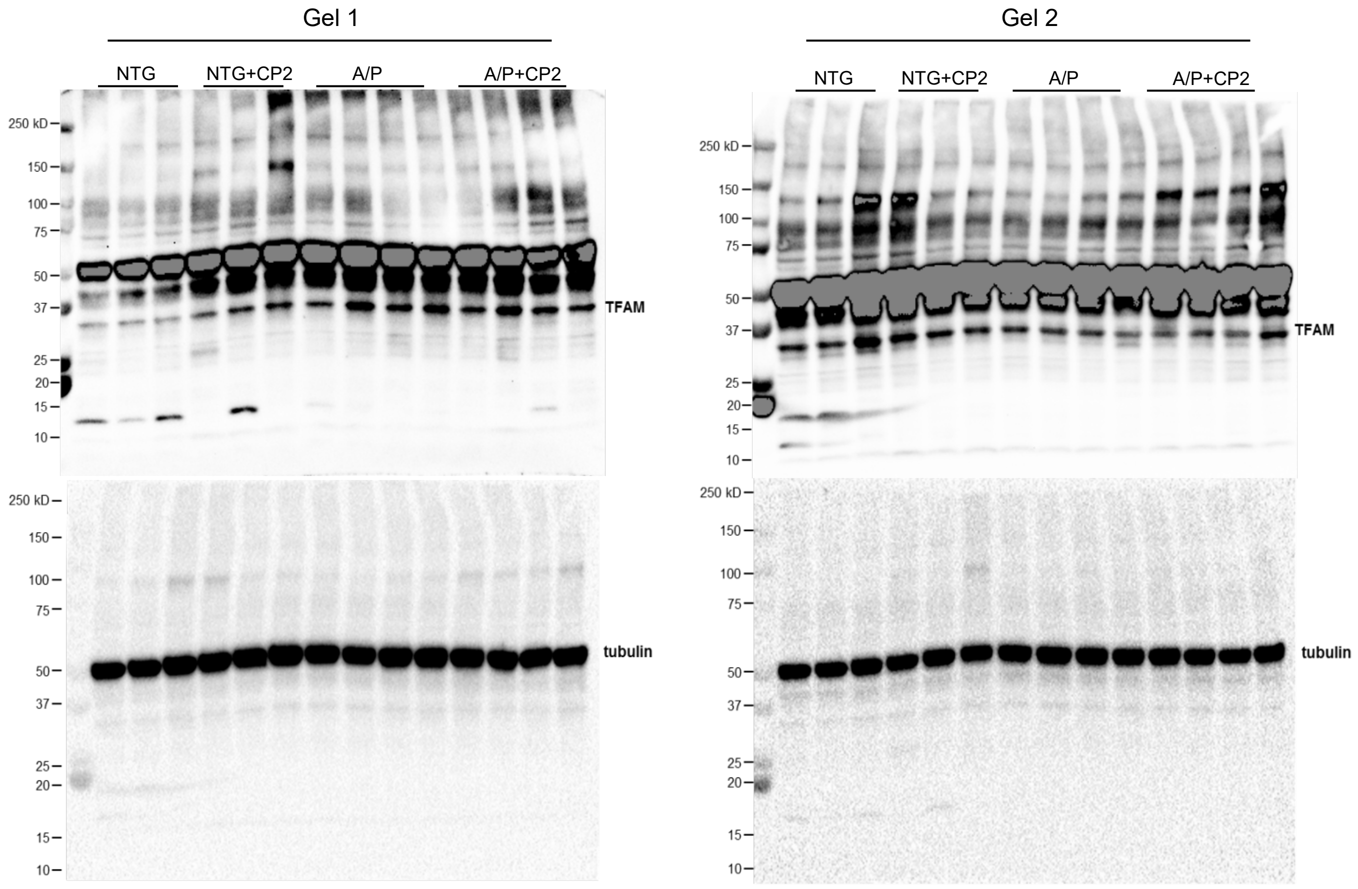


**Supplementary Fig. 4. Original uncropped Western blot images for Fig. 3k.** Cortico-hippocampal region of 6 - 8 mice *per* group was taken for Western blot analysis (NTG, *n* = 6; NTG+CP2, *n* = 6; APP/PS1, *n* = 8; APP/PS1+CP2, *n* = 8).
