## Supplementary Tables for "Partial inhibition of mitochondrial complex I attenuates neurodegeneration and restores energy homeostasis and synaptic function in a symptomatic Alzheimer’s mouse model"

**Supplementary Table 1.** Results of the Eurofin Cerep Safety-Screen 44 Panel (receptors and ion channels)

| Assay | Test concentration (M) | % Inhibition of Control Specific Binding |
|---|---|---|
| A2A (h) (agonist radioligand) | 1.0E-05 | 15 |
| alpha 1A (h) (antagonist radioligand) | 1.0E-05 | 12 |
| alpha 2A (h) (antagonist radioligand) | 1.0E-05 | -7 |
| beta 1 (h) (agonist radioligand) | 1.0E-05 | 5 |
| beta 2 (h) (agonist radioligand) | 1.0E-05 | -1 |
| BZD (central) (agonist radioligand) | 1.0E-05 | 11 |
| CB1 (h) (agonist radioligand) | 1.0E-05 | -14 |
| CB2 (h) (agonist radioligand) | 1.0E-05 | -6 |
| CCK1 (CCKA) (h) (agonist radioligand) | 1.0E-05 | -3 |
| D1 (h) (antagonist radioligand) | 1.0E-05 | -9 |
| D2S (h) (agonist radioligand) | 1.0E-05 | 0 |
| ETA (h) (agonist radioligand) | 1.0E-05 | -1 |
| NMDA (antagonist radioligand) | 1.0E-05 | 3 |
| H1 (h) (antagonist radioligand) | 1.0E-05 | 8 |
| H2 (h) (antagonist radioligand) | 1.0E-05 | -3 |
| MAO-A (antagonist radioligand) | 1.0E-05 | 25 |
| M1 (h) (antagonist radioligand) | 1.0E-05 | 35 |
| M2 (h) (antagonist radioligand) | 1.0E-05 | 50 |
| M3 (h) (antagonist radioligand) | 1.0E-05 | 30 |
| N neuronal alpha 4beta 2 (h) (agonist radioligand) | 1.0E-05 | 5 |
| delta 2 (DOP) (h) (agonist radioligand) | 1.0E-05 | 26 |
| kappa (KOP) (agonist radioligand) | 1.0E-05 | 3 |
| mu (MOP) (h) (agonist radioligand) | 1.0E-05 | 28 |
| 5-HT1A (h) (agonist radioligand) | 1.0E-05 | 0 |
| 5-HT1B (antagonist radioligand) | 1.0E-05 | -12 |
| 5-HT2A (h) (agonist radioligand) | 1.0E-05 | -2 |
| 5-HT2B (h) (agonist radioligand) | 1.0E-05 | 1 |
| 5-HT3 (h) (antagonist radioligand) | 1.0E-05 | 0 |
| GR (h) (agonist radioligand) | 1.0E-05 | 3 |
| AR (h) (agonist radioligand) | 1.0E-05 | 18 |
| V1a (h) (agonist radioligand) | 1.0E-05 | 10 |
| Ca2+ channel (L, dihydropyridine site) (antagonist radioligand) | 1.0E-05 | 8 |
| Potassium Channel hERG (human), [3H] Dofetilide | 1.0E-05 | 71 |
| KV channel (antagonist radioligand) | 1.0E-05 | -4 |
| Na+ channel (site 2) (antagonist radioligand) | 1.0E-05 | 34 |
| norepinephrine transporter (h) (antagonist radioligand) | 1.0E-05 | 8 |
| dopamine transporter (h) (antagonist radioligand) | 1.0E-05 | 32 |
| 5-HT transporter (h) (antagonist radioligand) | 1.0E-05 | 0 |

**In vitro pharmacological profiling and assessment of the potential for off-target interactions of CP2 in binding screens (Eurofins Cerep-Panlabs SafetyScreen 44).** The compound was screened at a 10 μM concentration in duplicate for its potential to interfere with the binding of native ligands of 44 different receptors, ion channels and enzymes. For receptor assays, compound binding was calculated as percent inhibition of the binding of a radioactively labeled ligand specific for each target. Tests were performed in duplicate and results are averages of those two tests. Results showing an inhibition (or stimulation for assays run in basal conditions) higher than 50% are considered to represent significant effect of the test compound. Results showing an inhibition (or stimulation) between 25% and 50% are indicative of weak to moderate effect. Results showing an inhibition (or stimulation) lower than 25% are not considered significant and mostly attributable to variability of the signal around the control level.

**Supplementary Table 2.** Results of the Eurofin Cerep Safety-Screen 44 Panel (enzymes)

| Assay | Test Concentration (M) | % Inhibition of Control Values |
|---|---|---|
| COX1 (h) | 1.0E-05 | 17 |
| COX2 (h) | 1.0E-05 | 17 |
| PDE3A (h) | 1.0E-05 | 13 |
| PDE4D2 (h) | 1.0E-05 | 28 |
| Lck kinase (h) | 1.0E-05 | -5 |
| acetylcholinesterase (h) | 1.0E-05 | 33 |

**In vitro pharmacological profiling and assessment of the potential for off-target interactions of CP2 in binding screens (Eurofins Cerep-Panlabs SafetyScreen 44).** The compound was screened at a 10 μM concentration in duplicate for its potential to interfere with the binding of native ligands of 44 different receptors, ion channels and enzymes. For enzyme assays, the inhibition effect was calculated as percent inhibition of control enzyme activity. Tests were performed in duplicate and results are averages of those two tests. Results showing an inhibition (or stimulation for assays run in basal conditions) higher than 50% are considered to represent significant effect of the test compound. Results showing an inhibition (or stimulation) between 25% and 50% are indicative of weak to moderate effect. Results showing an inhibition (or stimulation) lower than 25% are not considered significant and mostly attributable to variability of the signal around the control level.

**Supplementary Table 3**. Results of kinome profiling for CP2 in the Nanosyn 250 Kinase panel.
The number represents % inhibition at the concentration tested. Negative numbers represent increased enzyme activity. Numbers below 20 considered to be within the acceptable noise level within the plate.

| Kinase | Conc. Tested (μM) | CP2 |
|---|---|---|
| ABL1 | 1 | -4 |
| ABL1 | 10 | 4 |
| AKT1 | 1 | 3 |
| AKT1 | 10 | 6 |
| AKT2 | 1 | 1 |
| AKT2 | 10 | -4 |
| AKT3 | 1 | 0 |
| AKT3 | 10 | 1 |
| ALK | 1 | 2 |
| ALK | 10 | -2 |
| AMP-A1B1G1 | 1 | -3 |
| AMP-A1B1G1 | 10 | -13 |
| AMP-A2B1G1 | 1 | 3 |
| AMP-A2B1G1 | 10 | 1 |
| ARG | 1 | -1 |
| ARG | 10 | 2 |
| ARK5 | 1 | -3 |
| ARK5 | 10 | 17 |
| AURORA-A | 1 | 3 |
| AURORA-A | 10 | -4 |
| AURORA-B | 1 | 1 |
| AURORA-B | 10 | 7 |
| AURORA-C | 1 | 3 |
| AURORA-C | 10 | -7 |
| AXL | 1 | 2 |
| AXL | 10 | -7 |

| Kinase | Conc. Tested (μM) | CP2 |
|---|---|---|
| BLK | 1 | 36 |
| BLK | 10 | 34 |
| BMX | 1 | 3 |
| BMX | 10 | -6 |
| BRAF | 1 | 0 |
| BRAF | 10 | -4 |
| BRK | 1 | -3 |
| BRK | 10 | -15 |
| BRSK1 | 1 | -2 |
| BRSK1 | 10 | -4 |
| BRSK2 | 1 | -10 |
| BRSK2 | 10 | -14 |
| BTK | 1 | 1 |
| BTK | 10 | 1 |
| CAMK1A | 1 | 2 |
| CAMK1A | 10 | 1 |
| CAMK1D | 1 | 2 |
| CAMK1D | 10 | -4 |
| CAMK2A | 1 | -1 |
| CAMK2A | 10 | 0 |
| CAMK2B | 1 | 1 |
| CAMK2B | 10 | 4 |
| CAMK2D | 1 | -2 |
| CAMK2D | 10 | -1 |
| CAMK2G | 1 | -5 |
| CAMK2G | 10 | -7 |

| Kinase | Conc. Tested (μM) | CP2 |
|---|---|---|
| CAMK4 | 1 | 0 |
| CAMK4 | 10 | -6 |
| CDK1 | 1 | -4 |
| CDK1 | 10 | 0 |
| CDK2 | 1 | -3 |
| CDK2 | 10 | -4 |
| CDK2-CYCLINE | 1 | 4 |
| CDK2-CYCLINE | 10 | 1 |
| CDK3-CYCLINE | 1 | 1 |
| CDK3-CYCLINE | 10 | -2 |
| CDK4-CYCLIND | 1 | -1 |
| CDK4-CYCLIND | 10 | 3 |
| CDK5 | 1 | 0 |
| CDK5 | 10 | 3 |
| CDK5-P25 | 1 | 8 |
| CDK5-P25 | 10 | 2 |
| CDK6-CYCLIND3 | 1 | 0 |
| CDK6-CYCLIND3 | 10 | -3 |
| CDK7 | 1 | 4 |
| CDK7 | 10 | 11 |
| CDK9-CYCLINT1 | 1 | 5 |
| CDK9-CYCLINT1 | 10 | -26 |
| CHEK1 | 1 | -1 |
| CHEK1 | 10 | -10 |
| CHEK2 | 1 | 1 |
| CHEK2 | 10 | -1 |
| CK1 | 1 | 1 |
| CK1 | 10 | 19 |

| Kinase | Conc. Tested (μM) | CP2 |
|---|---|---|
| CK1-EPSILON | 1 | 2 |
| CK1-EPSILON | 10 | 30 |
| CK1-GAMMA1 | 1 | 1 |
| CK1-GAMMA1 | 10 | 19 |
| CK1-GAMMA2 | 1 | -3 |
| CK1-GAMMA2 | 10 | 10 |
| CK1-GAMMA3 | 1 | 1 |
| CK1-GAMMA3 | 10 | 15 |
| CLK1 | 1 | -6 |
| CLK1 | 10 | -6 |
| CLK2 | 1 | -2 |
| CLK2 | 10 | 3 |
| CLK3 | 1 | 1 |
| CLK3 | 10 | -2 |
| CLK4 | 1 | 5 |
| CLK4 | 10 | 24 |
| CRAF | 1 | -1 |
| CRAF | 10 | 0 |
| CSK | 1 | 0 |
| CSK | 10 | -5 |
| DAPK1 | 1 | 0 |
| DAPK1 | 10 | -1 |
| DAPK3 | 1 | -2 |
| DAPK3 | 10 | 0 |
| DCAMKL2 | 1 | -2 |
| DCAMKL2 | 10 | -5 |
| DDR1 | 1 | 3 |
| DDR1 | 10 | 16 |

**Supplementary Table 3**.

| Kinase | Conc. Tested (μM) | CP2 |
|---|---|---|
| DDR2 | 1 | 4 |
| DDR2 | 10 | 2 |
| DYRK1A | 1 | 4 |
| DYRK1A | 10 | 3 |
| DYRK1B | 1 | 2 |
| DYRK1B | 10 | 6 |
| DYRK2 | 1 | -7 |
| DYRK2 | 10 | 1 |
| DYRK3 | 1 | 1 |
| DYRK3 | 10 | 2 |
| DYRK4 | 1 | -2 |
| DYRK4 | 10 | -3 |
| EGFR | 1 | 5 |
| EGFR | 10 | -12 |
| EPH-A1 | 1 | -1 |
| EPH-A1 | 10 | -9 |
| EPH-A2 | 1 | 1 |
| EPH-A2 | 10 | 0 |
| EPH-A3 | 1 | -2 |
| EPH-A3 | 10 | -14 |
| EPH-A4 | 1 | 0 |
| EPH-A4 | 10 | 1 |
| EPH-A5 | 1 | -3 |
| EPH-A5 | 10 | -10 |
| EPH-A8 | 1 | 0 |
| EPH-A8 | 10 | -9 |
| EPH-B1 | 1 | -3 |
| EPH-B1 | 10 | -12 |
| EPH-B2 | 1 | -3 |

| Kinase | Conc. Tested (μM) | CP2 |
|---|---|---|
| EPH-B2 | 10 | 2 |
| EPH-B3 | 1 | -3 |
| EPH-B3 | 10 | -10 |
| EPH-B4 | 1 | -4 |
| EPH-B4 | 10 | -12 |
| ERB-B2 | 1 | -6 |
| ERB-B2 | 10 | -13 |
| ERB-B4 | 1 | 8 |
| ERB-B4 | 10 | -1 |
| FAK | 1 | 0 |
| FAK | 10 | -3 |
| FER | 1 | -1 |
| FER | 10 | 1 |
| FES | 1 | 5 |
| FES | 10 | -3 |
| FGFR1 | 1 | 5 |
| FGFR1 | 10 | -15 |
| FGFR2 | 1 | 1 |
| FGFR2 | 10 | -3 |
| FGFR4 | 1 | 0 |
| FGFR4 | 10 | -4 |
| FGR | 1 | 0 |
| FGR | 10 | -38 |
| FLT-1 | 1 | 4 |
| FLT-1 | 10 | -23 |
| FLT-3 | 1 | 0 |
| FLT-3 | 10 | 0 |
| FLT-4 | 1 | 1 |
| FLT-4 | 10 | 2 |

| Kinase | Conc. Tested (μM) | CP2 |
|---|---|---|
| FMS | 1 | 4 |
| FMS | 10 | -1 |
| FRAP1 | 1 | 2 |
| FRAP1 | 10 | 22 |
| FYN | 1 | 1 |
| FYN | 10 | 5 |
| GRK6 | 1 | -2 |
| GRK6 | 10 | -5 |
| GRK7 | 1 | 0 |
| GRK7 | 10 | -12 |
| GSK-3-ALPHA | 1 | 5 |
| GSK-3-ALPHA | 10 | 47 |
| GSK-3-BETA | 1 | -4 |
| GSK-3-BETA | 10 | 21 |
| HASPIN | 1 | 2 |
| HASPIN | 10 | -1 |
| HCK | 1 | 9 |
| HCK | 10 | -1 |
| HIPK1 | 1 | -6 |
| HIPK1 | 10 | 2 |
| HIPK2 | 1 | 0 |
| HIPK2 | 10 | 0 |
| HIPK3 | 1 | -3 |
| HIPK3 | 10 | -1 |
| HIPK4 | 1 | -2 |
| HIPK4 | 10 | -4 |
| IGF1R | 1 | 4 |
| IGF1R | 10 | -4 |
| IKK-ALPHA | 1 | 0 |

| Kinase | Conc. Tested (μM) | CP2 |
|---|---|---|
| IKK-ALPHA | 10 | -3 |
| IKK-BETA | 1 | 0 |
| IKK-BETA | 10 | 6 |
| IKK-EPSILON | 1 | -1 |
| IKK-EPSILON | 10 | 1 |
| INSR | 1 | -1 |
| INSR | 10 | 1 |
| IRAK1 | 1 | -4 |
| IRAK1 | 10 | -4 |
| IRAK4 | 1 | 0 |
| IRAK4 | 10 | 3 |
| IRR | 1 | 4 |
| IRR | 10 | -1 |
| ITK | 1 | 4 |
| ITK | 10 | -3 |
| JAK1 | 1 | -2 |
| JAK1 | 10 | 2 |
| JAK2 | 1 | 4 |
| JAK2 | 10 | 0 |
| JAK3 | 1 | -2 |
| JAK3 | 10 | -11 |
| JNK1 | 1 | 2 |
| JNK1 | 10 | 6 |
| JNK2 | 1 | 6 |
| JNK2 | 10 | -3 |
| JNK3 | 1 | 4 |
| JNK3 | 10 | 9 |
| KDR | 1 | 6 |
| KDR | 10 | -3 |

**Supplementary Table 3**.

| Kinase | Conc. Tested (μM) | CP2 |
|---|---|---|
| KIT | 1 | 0 |
| KIT | 10 | -6 |
| LATS1 | 1 | 0 |
| LATS1 | 10 | -9 |
| LATS2 | 1 | 0 |
| LATS2 | 10 | 0 |
| LCK | 1 | 2 |
| LCK | 10 | -10 |
| LOK | 1 | 4 |
| LOK | 10 | 14 |
| LRRK2-G2019S | 1 | 0 |
| LRRK2-G2019S | 10 | 3 |
| LTK | 1 | 2 |
| LTK | 10 | -6 |
| LYNA | 1 | 1 |
| LYNA | 10 | -2 |
| LYNB | 1 | 2 |
| LYNB | 10 | -18 |
| MAP4K2 | 1 | 3 |
| MAP4K2 | 10 | 2 |
| MAP4K4 | 1 | 4 |
| MAP4K4 | 10 | 1 |
| MAP4K5 | 1 | 1 |
| MAP4K5 | 10 | 4 |
| MAPK1 | 1 | -1 |
| MAPK1 | 10 | -10 |
| MAPK3 | 1 | -1 |
| MAPK3 | 10 | 3 |
| MAPKAPK-2 | 1 | -4 |

| Kinase | Conc. Tested (μM) | CP2 |
|---|---|---|
| MAPKAPK-2 | 10 | 2 |
| MAPKAPK-3 | 1 | -1 |
| MAPKAPK-3 | 10 | -3 |
| MARK1 | 1 | 2 |
| MARK1 | 10 | 3 |
| MARK3 | 1 | -4 |
| MARK3 | 10 | -4 |
| MARK4 | 1 | -3 |
| MARK4 | 10 | 0 |
| MEK1 | 1 | 2 |
| MEK1 | 10 | -1 |
| MEK2 | 1 | -1 |
| MEK2 | 10 | 6 |
| MEK3 | 1 | 0 |
| MEK3 | 10 | 2 |
| MELK | 1 | 6 |
| MELK | 10 | 3 |
| MER | 1 | 2 |
| MER | 10 | -1 |
| MET | 1 | 8 |
| MET | 10 | -9 |
| MINK | 1 | 0 |
| MINK | 10 | 10 |
| MKNK1 | 1 | 0 |
| MKNK1 | 10 | 5 |
| MNK2 | 1 | 5 |
| MNK2 | 10 | 5 |
| MRCK-ALPHA | 1 | -2 |
| MRCK-ALPHA | 10 | -3 |

| Kinase | Conc. Tested (μM) | CP2 |
|---|---|---|
| MRCK-BETA | 1 | 0 |
| MRCK-BETA | 10 | -2 |
| MSK1 | 1 | -1 |
| MSK1 | 10 | -13 |
| MSK2 | 1 | 1 |
| MSK2 | 10 | -11 |
| MSSK1 | 1 | -3 |
| MSSK1 | 10 | 2 |
| MST1 | 1 | 3 |
| MST1 | 10 | 22 |
| MST2 | 1 | 5 |
| MST2 | 10 | 10 |
| MST3 | 1 | -2 |
| MST3 | 10 | -3 |
| MST4 | 1 | -2 |
| MST4 | 10 | -5 |
| MUSK | 1 | 4 |
| MUSK | 10 | -6 |
| NDR2 | 1 | 1 |
| NDR2 | 10 | -4 |
| NDRG1 | 1 | 0 |
| NDRG1 | 10 | -2 |
| NEK1 | 1 | 3 |
| NEK1 | 10 | 7 |
| NEK2 | 1 | 0 |
| NEK2 | 10 | -1 |
| NEK6 | 1 | 4 |
| NEK6 | 10 | 1 |
| NEK7 | 1 | 3 |

| Kinase | Conc. Tested (μM) | CP2 |
|---|---|---|
| NEK7 | 10 | -20 |
| NEK9 | 1 | 2 |
| NEK9 | 10 | 5 |
| P38-ALPHA | 1 | 4 |
| P38-ALPHA | 10 | 1 |
| P38-BETA | 1 | 3 |
| P38-BETA | 10 | 1 |
| P38-DELTA | 1 | 0 |
| P38-DELTA | 10 | -2 |
| P38-GAMMA | 1 | 0 |
| P38-GAMMA | 10 | -7 |
| P70S6K1 | 1 | 3 |
| P70S6K1 | 10 | -1 |
| P70S6K2 | 1 | 1 |
| P70S6K2 | 10 | -14 |
| PAK1 | 1 | 0 |
| PAK1 | 10 | 2 |
| PAK2 | 1 | 0 |
| PAK2 | 10 | -2 |
| PAK3 | 1 | -1 |
| PAK3 | 10 | -3 |
| PAK4 | 1 | 5 |
| PAK4 | 10 | 10 |
| PAK5 | 1 | 1 |
| PAK5 | 10 | 3 |
| PAK6 | 1 | 4 |
| PAK6 | 10 | -3 |
| PAR-1B-ALPHA | 1 | -3 |
| PAR-1B-ALPHA | 10 | 7 |

**Supplementary Table 3**.

| Kinase | Conc. Tested (μM) | CP2 |
|---|---|---|
| PASK | 1 | -5 |
| PASK | 10 | 2 |
| PDGFR-ALPHA | 1 | 3 |
| PDGFR-ALPHA | 10 | 3 |
| PDGFR-BETA | 1 | 4 |
| PDGFR-BETA | 10 | -1 |
| PDK1 | 1 | -4 |
| PDK1 | 10 | 0 |
| PERK | 1 | 2 |
| PERK | 10 | -14 |
| PHK-GAMMA1 | 1 | 3 |
| PHK-GAMMA1 | 10 | 0 |
| PHK-GAMMA2 | 1 | 1 |
| PHK-GAMMA2 | 10 | 5 |
| PI3-KINASE-ALPHA | 1 | 2 |
| PI3-KINASE-ALPHA | 10 | 7 |
| PI4-K-BETA | 1 | 0 |
| PI4-K-BETA | 10 | -3 |
| PIM-1-KINASE | 1 | 0 |
| PIM-1-KINASE | 10 | 1 |
| PIM2 | 1 | 1 |
| PIM2 | 10 | -2 |
| PIM3 | 1 | -1 |
| PIM3 | 10 | 0 |
| PKA | 1 | 2 |
| PKA | 10 | -11 |
| PKACB | 1 | -1 |
| PKACB | 10 | 0 |
| PKC-ALPHA | 1 | 1 |
| PKC-ALPHA | 10 | 0 |
| PKC-BETA1 | 1 | 0 |
| PKC-BETA1 | 10 | 8 |
| PKC-BETA2 | 1 | 2 |
| PKC-BETA2 | 10 | 0 |
| PKC-ETA | 1 | -2 |
| PKC-ETA | 10 | 1 |
| PKC-GAMMA | 1 | 2 |
| PKC-GAMMA | 10 | 1 |
| PKC-IOTA | 1 | -6 |
| PKC-IOTA | 10 | 3 |
| PKC-THETA | 1 | -3 |
| PKC-THETA | 10 | -1 |
| PKC-ZETA | 1 | -1 |
| PKC-ZETA | 10 | -3 |
| PKN1 | 1 | -1 |
| PKN1 | 10 | -4 |
| PKN2 | 1 | 2 |
| PKN2 | 10 | 1 |
| PLK1 | 1 | -1 |
| PLK1 | 10 | -19 |
| PLK3 | 1 | -5 |
| PLK3 | 10 | 14 |
| PLK4 | 1 | -9 |
| PLK4 | 10 | -11 |
| PRAK | 1 | -3 |
| PRAK | 10 | -8 |
| PRKACA | 1 | -2 |
| PRKACA | 10 | -2 |
| PRKD1 | 1 | 7 |
| PRKD1 | 10 | -4 |
| PRKD2 | 1 | 0 |
| PRKD2 | 10 | 2 |
| PRKD3 | 1 | 5 |
| PRKD3 | 10 | -3 |
| PRKG1 | 1 | 2 |
| PRKG1 | 10 | 4 |
| PRKX | 1 | 0 |
| PRKX | 10 | -1 |
| PTK5 | 1 | -1 |
| PTK5 | 10 | -3 |
| PYK2 | 1 | 5 |
| PYK2 | 10 | -8 |
| RET | 1 | 3 |
| RET | 10 | -4 |
| RIPK2 | 1 | 0 |
| RIPK2 | 10 | -5 |
| ROCK1 | 1 | 5 |
| ROCK1 | 10 | -1 |
| ROCK2 | 1 | 7 |
| ROCK2 | 10 | 16 |
| RON | 1 | 8 |
| RON | 10 | -12 |
| ROS | 1 | 4 |
| ROS | 10 | -5 |
| RSK1 | 1 | 2 |
| RSK1 | 10 | 0 |
| RSK2 | 1 | -2 |
| RSK2 | 10 | -6 |
| RSK3 | 1 | -3 |
| RSK3 | 10 | -1 |
| RSK4 | 1 | -1 |
| RSK4 | 10 | -1 |
| SGK1 | 1 | 1 |
| SGK1 | 10 | 1 |
| SGK2 | 1 | -4 |
| SGK2 | 10 | 1 |
| SGK3 | 1 | 5 |
| SGK3 | 10 | -5 |
| SIK | 1 | 4 |
| SIK | 10 | 1 |
| SLK | 1 | -2 |
| SLK | 10 | 4 |
| SNF1LK2 | 1 | 4 |
| SNF1LK2 | 10 | 1 |
| SPHK1 | 1 | 3 |
| SPHK1 | 10 | 0 |
| SPHK2 | 1 | 4 |
| SPHK2 | 10 | -4 |
| SRC | 1 | 2 |
| SRC | 10 | -18 |
| SRMS | 1 | -6 |
| SRMS | 10 | -10 |
| SRPK1 | 1 | 3 |
| SRPK1 | 10 | -8 |
| SRPK2 | 1 | -3 |

**Supplementary Table 3**.

| Kinase | Conc. Tested (μM) | CP2 |
|---|---|---|
| SRPK2 | 10 | 0 |
| STK16 | 1 | 1 |
| STK16 | 10 | -5 |
| STK25 | 1 | -1 |
| STK25 | 10 | 2 |
| SYK | 1 | 6 |
| SYK | 10 | -15 |
| TAK1-TAB1 | 1 | -1 |
| TAK1-TAB1 | 10 | -1 |
| TAOK2 | 1 | -2 |
| TAOK2 | 10 | -1 |
| TAOK3 | 1 | 3 |
| TAOK3 | 10 | -2 |
| TBK1 | 1 | -2 |
| TBK1 | 10 | -7 |
| TEC | 1 | 2 |
| TEC | 10 | 1 |
| TIE2 | 1 | 0 |
| TIE2 | 10 | -9 |
| TNIK | 1 | 1 |
| TNIK | 10 | 1 |
| TNK2 | 1 | 3 |
| TNK2 | 10 | 2 |
| TRKA | 1 | 2 |
| TRKA | 10 | -5 |
| TRKB | 1 | 5 |
| TRKB | 10 | -3 |
| TRKC | 1 | 3 |
| TRKC | 10 | -3 |

| Kinase | Conc. Tested (μM) | CP2 |
|---|---|---|
| TSSK1 | 1 | -2 |
| TSSK1 | 10 | 7 |
| TSSK2 | 1 | -2 |
| TSSK2 | 10 | 1 |
| TTK | 1 | 2 |
| TTK | 10 | 4 |
| TXK | 1 | 6 |
| TXK | 10 | -5 |
| TYK2 | 1 | 4 |
| TYK2 | 10 | -7 |
| TYRO3 | 1 | 0 |
| TYRO3 | 10 | -4 |
| YES | 1 | -1 |
| YES | 10 | -4 |
| ZAP70 | 1 | -15 |
| ZAP70 | 10 | -3 |

**Supplementary Table 4.** CP2 levels in the brain tissue and plasma of NTG and APP/PS1 mice 22 – 24-month-old chronically treated from 9 months of age.

| Genotype | Brain (ng/g) | Brain (nM) | Plasma (ng/ml) | Plasma (nM) |
|---|---|---|---|---|
| NTG | 24.77 | 62.96 | 44.44 | 112.95 |
| NTG | 16.91 | 42.98 | 138.32 | 351.56 |
| NTG | 14.57 | 37.03 | 40.95 | 104.08 |
| NTG | 14.44 | 36.70 | 138.28 | 351.46 |
| NTG | 52.22 | 132.72 | 419.32 | 1,065.75 |
| NTG | 110.37 | 280.52 | 66.7 | 169.53 |
| NTG | 9.97 | 25.34 | 21.96 | 55.81 |
| APP/PS1 | 7.34 | 18.66 | 187.06 | 475.44 |
| APP/PS1 | 44.37 | 112.77 | 349.88 | 889.26 |
| APP/PS1 | 4.52 | 11.49 | 67.44 | 171.41 |
| APP/PS1 | 10.36 | 26.33 | 161.3 | 409.96 |
| APP/PS1 | 50.23 | 127.67 | 89.664 | 227.89 |
| APP/PS1 | 16.36 | 41.58 | 174.72 | 444.07 |
| APP/PS1 | 11.99 | 30.47 | 32.42 | 82.40 |
| APP/PS1 | 16.37 | 41.61 | 28.31 | 71.95 |
| APP/PS1 | 14.23 | 36.17 | 10.69 | 27.17 |
| APP/PS1 | 9.15 | 23.26 | 67.22 | 170.85 |
| APP/PS1 | 11.51 | 29.25 | 52.76 | 134.10 |

CP2 levels were measured in cerebellum. Blood for this analysis was collected at the end of the study at the time of sacrifice.

**Supplementary Table 5.** Effect of CP2 treatment on brain metabolite levels in APP/PS1 and NTG mice.

| | APP/PS1 | | APP/PS1 (CP2) | | t-test APP/PS1-CP2 | NTG | | NTG(CP2) | | t-test NTG-CP2 |
|---|---|---|---|---|---|---|---|---|---|---|
| **BRAIN** | **X̄** | **±SE** | **X̄** | **±SE** | **P values** | **X̄** | **±SE** | **X̄** | **±SE** | **P values** |
| **2-Hydroxyglutarate** | 1.28 | 0.10 | 1.19 | 0.11 | 0.307 | 0.76 | 0.09 | 0.93 | 0.20 | 0.265 |
| **4-Aminobutyrate** | 0.89 | 0.05 | 1.07 | 0.07 | 0.062 | 0.98 | 0.15 | 0.96 | 0.04 | 0.448 |
| **Adenosine** | 1.25 | 0.09 | 0.97 | 0.16 | 0.119 | 0.96 | 0.18 | 0.76 | 0.06 | 0.195 |
| **ADP** | 0.89 | 0.08 | 1.08 | 0.06 | 0.078 | 1.03 | 0.09 | 1.01 | 0.08 | 0.458 |
| **Alanine** | 0.94 | 0.02 | 1.27 | 0.11 | ***0.033*** | 0.96 | 0.07 | 0.94 | 0.07 | 0.433 |
| **AMP** | 0.97 | 0.04 | 1.20 | 0.02 | ***0.003*** | 0.89 | 0.04 | 0.91 | 0.05 | 0.395 |
| **Ascorbate** | 0.95 | 0.14 | 1.59 | 0.22 | ***0.039*** | 0.76 | 0.18 | 0.74 | 0.20 | 0.472 |
| **Aspartate** | 0.52 | 0.05 | 0.90 | 0.22 | 0.107 | 0.94 | 0.18 | 1.35 | 0.16 | 0.088 |
| **ATP** | 0.78 | 0.15 | 1.15 | 0.23 | 0.140 | 1.10 | 0.18 | 1.02 | 0.14 | 0.387 |
| **β-Alanine** | 0.30 | 0.11 | 1.11 | 0.08 | ***0.001*** | 0.88 | 0.22 | 0.86 | 0.10 | 0.469 |
| **Cholestrol** | 0.84 | 0.20 | 0.69 | 0.29 | 0.359 | 1.18 | 0.38 | 1.46 | 0.51 | 0.351 |
| **Citrate** | 0.91 | 0.04 | 1.04 | 0.01 | ***0.029*** | 1.07 | 0.04 | 0.98 | 0.09 | 0.238 |
| **Creatinine** | 1.02 | 0.11 | 1.13 | 0.03 | 0.216 | 1.04 | 0.08 | 0.81 | 0.08 | 0.058 |
| **Dehydroascorbate** | 0.70 | 0.06 | 1.61 | 0.26 | ***0.025*** | 0.83 | 0.17 | 1.01 | 0.21 | 0.294 |
| **Ethanolamine** | 0.89 | 0.16 | 1.17 | 0.07 | 0.118 | 1.07 | 0.11 | 1.13 | 0.07 | 0.333 |
| **Fumarate** | 0.67 | 0.32 | 1.13 | 0.26 | 0.181 | 0.56 | 0.23 | 1.17 | 0.21 | 0.064 |
| **Galactose** | 0.95 | 0.12 | 1.03 | 0.24 | 0.395 | 0.84 | 0.09 | 1.15 | 0.18 | 0.123 |
| **GDP** | 0.84 | 0.04 | 1.13 | 0.03 | ***0.002*** | 0.92 | 0.13 | 1.04 | 0.05 | 0.237 |
| **Glucopyranose** | 1.56 | 0.17 | 1.08 | 0.13 | ***0.045*** | 0.89 | 0.13 | 0.42 | 0.06 | ***0.019*** |
| **Glutamate** | 0.99 | 0.09 | 1.15 | 0.15 | 0.210 | 1.05 | 0.10 | 0.91 | 0.25 | 0.337 |
| **Glycerate** | 1.03 | 0.15 | 0.78 | 0.06 | 0.117 | 1.19 | 0.09 | 0.74 | 0.06 | ***0.005*** |
| **Glycerol -1-P** | 1.00 | 0.05 | 1.05 | 0.06 | 0.267 | 1.00 | 0.11 | 1.02 | 0.10 | 0.461 |
| **Glycine** | 0.69 | 0.04 | 0.88 | 0.08 | 0.053 | 0.97 | 0.10 | 1.10 | 0.03 | 0.159 |
| **Glycolate** | 0.56 | 0.21 | 0.17 | 0.00 | 0.094 | 0.69 | 0.12 | 1.00 | 0.12 | 0.081 |
| **GTP** | 0.88 | 0.05 | 1.19 | 0.07 | ***0.012*** | 0.98 | 0.07 | 0.96 | 0.08 | 0.441 |
| **Iminodiacetate** | 1.13 | 0.08 | 0.86 | 0.21 | 0.172 | 0.91 | 0.24 | 0.55 | 0.14 | 0.153 |
| **Lactate** | 1.10 | 0.03 | 1.04 | 0.01 | 0.057 | 0.93 | 0.06 | 0.95 | 0.01 | 0.393 |
| **Malate** | 0.96 | 0.09 | 0.98 | 0.07 | 0.448 | 1.02 | 0.02 | 1.09 | 0.11 | 0.319 |
| **Myo-Inositol** | 0.91 | 0.03 | 1.04 | 0.06 | 0.072 | 1.00 | 0.05 | 0.98 | 0.03 | 0.364 |
| **NAA** | 0.94 | 0.02 | 1.10 | 0.03 | ***0.004*** | 1.00 | 0.06 | 1.04 | 0.09 | 0.369 |
| **Phosphocolamine** | 0.84 | 0.07 | 1.16 | 0.13 | 0.061 | 0.90 | 0.07 | 1.11 | 0.05 | ***0.036*** |
| **Pi** | 0.86 | 0.07 | 0.98 | 0.02 | 0.119 | 1.00 | 0.04 | 1.07 | 0.04 | 0.152 |
| **Pyroglutamate** | 0.52 | 0.07 | 0.84 | 0.07 | ***0.011*** | 0.94 | 0.15 | 1.28 | 0.07 | 0.070 |
| **Serine** | 0.65 | 0.05 | 0.97 | 0.05 | ***0.003*** | 0.91 | 0.10 | 1.20 | 0.09 | ***0.050*** |
| **Stearate** | 1.03 | 0.04 | 0.93 | 0.05 | 0.111 | 0.96 | 0.16 | 1.15 | 0.14 | 0.225 |
| **Succinate** | 1.08 | 0.10 | 1.07 | 0.13 | 0.489 | 0.98 | 0.06 | 0.91 | 0.13 | 0.352 |
| **Threonate** | 0.89 | 0.07 | 1.03 | 0.14 | 0.225 | 1.17 | 0.10 | 1.10 | 0.13 | 0.375 |
| **Urea** | 1.04 | 0.08 | 1.20 | 0.03 | 0.092 | 0.94 | 0.08 | 0.71 | 0.04 | ***0.037*** |
| **Valine** | 0.47 | 0.11 | 0.96 | 0.10 | ***0.013*** | 0.79 | 0.14 | 0.43 | 0.08 | ***0.050*** |

Metabolomics was conducted in mice treated with CP2 or vehicle for 6 months. 5 mice were included in each group. Data were analyzed by unpaired Student *t*-test. $P < 0.05$ was considered significant.
